# Chromosome-length genome assembly of *Uta stansburiana* and gene expression data reveal fast pace-of-life comes with environmental stability

**DOI:** 10.1101/2025.05.28.656178

**Authors:** Sam R. Fellows, Raúl Araya-Donoso, Elizabeth Dávalos-Dehullu, Ruqayya Khan, David Weisz, Olga Dudchenko, Erez Lieberman Aiden, Adrián Munguía-Vega, Greer A. Dolby, Danielle L. Edwards

**Author notes:** Correspondence: SRF, GAD.

## Abstract

*Uta stansburiana* are an emerging model system for testing hypotheses regarding the evolution of pace-of-life syndromes (POLS) across its variable environments and wide latitudinal gradient. POLS are suites of traits related to variation of life history along a slow maturing-fast maturing continuum. We present a high-quality chromosome-level reference genome for *U. stansburiana* and use RNA-seq gene expression data to test for molecular correlates for pace-of-life differences between locations with higher and lower climate seasonality, UV differences, and sexual size dimorphism (SSD). Our assembly is 2.1 Gbp, has scaffold N50 of 320 Mbp, includes 104 scaffolds, and has an L50 of 3. The assembly comprises six macrochromosomes and 11 microchromosomes. We annotated 20,350 genes for the assembly and found a repeat element composition of 49.23%, similar to work in other phrynosomatid lizards. RNA-seq data reveal differential expression in genes consistent with pace-of-life differences and physiological variation including those related to stress, sexual reproduction, and cell proliferation/carcinogenesis between distinctive environments. Our results provide genes potentially underlying the molecular bases of POLS differences in a wild lizard.

## 1. Introduction

The pace-of-life syndrome (hereafter, POLS; Ricklefs and Wikelski 2002) hypothesis invokes balancing selection to explain how life history variation is maintained within and among populations (Royauté et al. 2018). The POLS hypothesis posits that life history, behavioral, and physiological (especially metabolism, immunity, and stress responses) traits coevolve along a “slow-fast” continuum which can create life history variation within or among populations (Ricklefs and Wikelski 2002; Réale et al. 2007, 2010). Fast POLSs are often characterized by fast metabolic rates, high stress responses, rapid development and maturation rates, bold exploratory behaviors or high aggressiveness, high reproductive output, and are short-lived (Réale et al. 2010; Dammhahn et al. 2018; Montiglio et al. 2018; but see also Royauté et al. 2018). Differences amongst POLS are likely exacerbated by variation in population density, e.g., lineages with high variation in population density tend to have the starkest differences in POLS (Wright et al 2018). This suggests that the role of the environment in the formation and maintenance of POLS is foundational (Dammhahn et al. 2018; Vasilieva 2022). As genomic and gene expression data become more readily available, it becomes easier to test and generate hypotheses regarding behavior (Hu et al. 2022), ecology (Xue et al. 2024), niche (Mellor et al. 2025), and life history (Narum et al. 2018). Immonen et al (2018) reviewed POLS literature and identified diverse molecular pathways which, when differentially expressed, may identify POLS differences including stress, immunity, metabolism, life span, and sexual signals among others. Differences in POLS among populations can also contribute to genetic divergence (Bolstad et al. 2014; Wright et al. 2019) To date, gene expression evidence for POLS has only been identified in house mice (Prabh et al. 2023) where faster pace-of-life is associated with genes related to metabolism, resulting in shorter generation time and more exploratory behavior. However, on a molecular level stress brought on by a “fast” pace-of-life may shorten telomeres, results in cellular damage due to oxidative stress, and often increases cancer risk (Giraudeau et al. 2019; Ujvari et al. 2022). Despite this, studies connecting the molecular mechanisms underlying POLS-related genes to such phenotypes are rare. Linking key pace-of-life processes to gene expression differences in genes associated with life history, behavior, physiology, and across environments using high-quality reference genomes presents an unprecedented opportunity to detect evidence for the POLS hypothesis in wild settings.

Common side-blotched lizards (*Uta stansburiana*, Baird and Girard 1852) are small, wide-spread phrynosomatids native to the southwestern United States and northwestern Mexico, whose range encompasses approximately 25 degrees latitude and 22 degrees longitude (Figure 1; Stebbins 2003). Side-blotched lizards are small-bodied (up to 8 g and 60 mm snout-vent length) and typically found amongst rocky scrub. They comprise a widespread species complex with broad variation in morphological and life history characteristics across the range including differences in sexual size dimorphism (SSD; Corl et al. 2012; Chelini et al. 2021), dorsal pattern (McKinney 1971), genetically-determined mating tactics and associated throat color polymorphism (Sinervo and Lively 1996; Sinervo et al. 2000; Corl et al. 2010), longevity (Zani and Stein 2018; Smith et al. 2019), and reproductive investment (Alonzo and Sinervo 2001; Smith et al. 2019). Such variation has been interpreted in the context of pace-of-life processes operating as an evolutionary mechanism within this species complex (Wright et al. 2019). Further, such patterns are relevant to studies of sexual selection (Schlippe Justicia et al. 2024) and speciation (Corl et al. 2012), so recovering the transcriptomic differences associated with these patterns can offer preliminary insights into POLS in an ideal, emerging model system.

**Figure 1.**
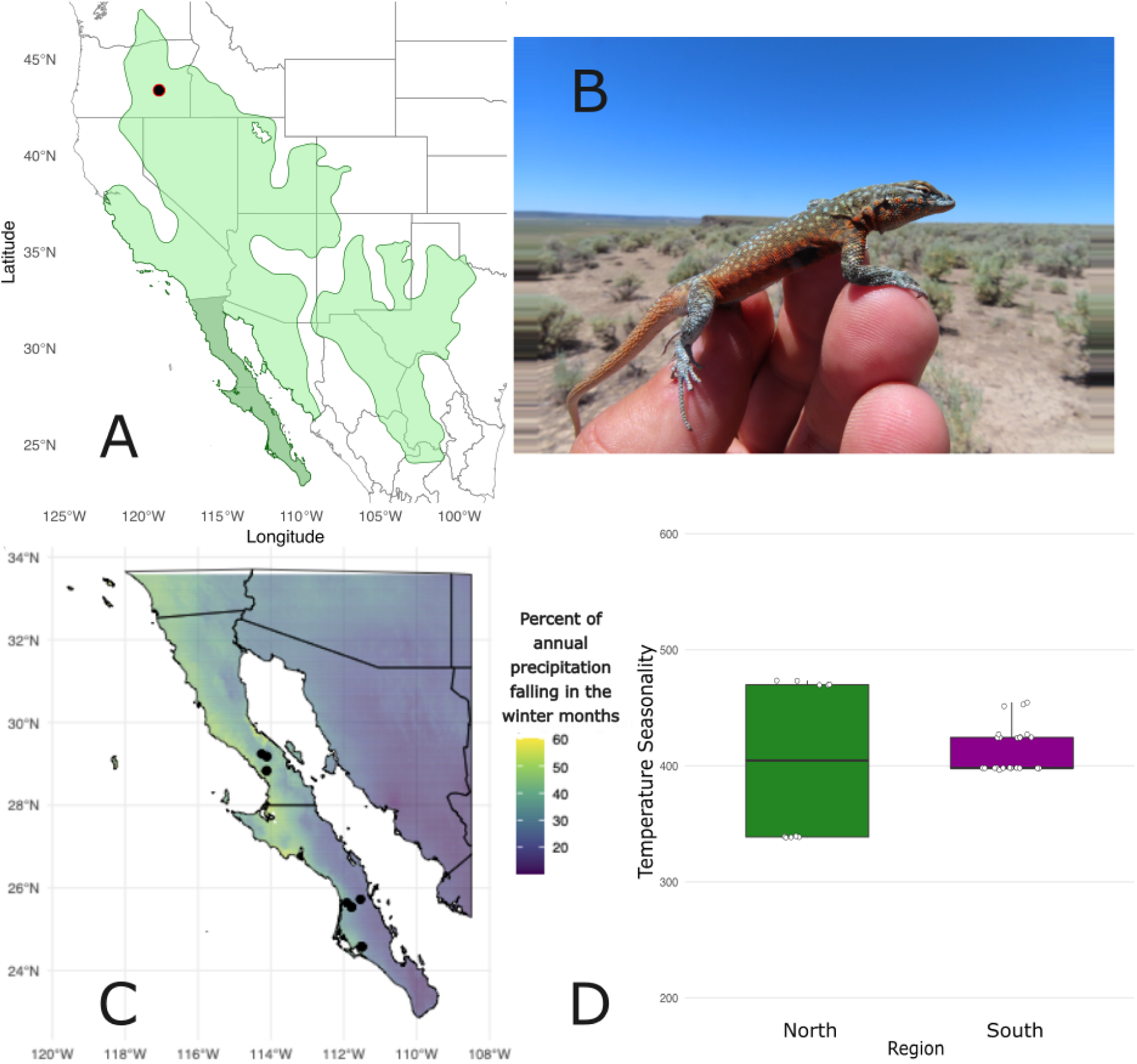
**A)** Map of full range of *Uta stansburiana* (IUCN Redlist) with collection locality of specimen from which the reference genome was sequenced (DLE 3124) and the study area in Baja California, MX is highlighted. **B)** Photo of the individual male *Uta stansburiana nevadensis* (DLE 3124) from whom the reference genome was generated (photo courtesy Dr. Pete Zani). **C)** Map of samples collected for RNA with percentage of average total annual precipitation occurring in the winter (December-February; Fick and Hijmans 2017). Samples represent 33 specimens collected across seven sites. Sites were divided into those north (3 populations, 9 samples) and south (4 populations, 24 samples) of the Vizcaíno desert. **D)** Differences in temperature seasonality (standard deviation of monthly temperatures x 100, BIO4; Fick and Hijmans 2017) between northern and southern sites.

Side-blotched lizards demonstrate strong life history trade-offs with respect to latitude across their range. Individuals from the southern portions of the species distribution in the southern United States and northern Mexico tend to be shorter-lived (up to two years, but usually one or less; Wilson 1991; Smith et al. 2019) and lay many smaller eggs relative to longer-lived individuals in the northern portions of the species range (i.e., up to 7 years in Oregon; Zani and Stein 2018) which lay fewer, larger eggs (Smith et al 2019). This transition in longevity exists alongside a range-wide shift from populations in the southern range with trimorphic throat colors to populations in the northern distribution that are monomorphic orange-throated (Corl et al. 2010), mediated by differences in seasonality and annual precipitation (Chelini et al. 2021). There have been suggestions that these dynamics are primarily underlain by sexual selection dynamics and population densities given evolutionary correlations between SSD and the degree of polymorphism (Corl et al. 2010b). However, recent evidence suggests that environments independently generate variation in polymorphism and SSD, and the variation in the latter occurs through sexually dimorphic growth rates in side-blotched lizards in alternate environmental conditions (Chelini et al. 2021). Sexually dimorphic growth (and variation in its degree) that correlates evolutionarily to alternate environmental conditions in different parts of the side-blotched lizard range indicate alternate sex-specific optima in POLS, and such variation may provide the mechanism connecting variation in SSD and polymorphism (Wright et al. 2019; Arnqvist and Rowe 2023) by altering sexual selection dynamics to reinforce divergence among populations in this system (Butlin and Smadja 2018). Furthermore, environmental conditions are known to influence metabolic and immunity trait differences associated with alternate growth rates in *U. stansburiana* (Smith et al. 2017) consistent with the POLS hypothesis (Dammhahn et al. 2018). We test for evidence of environmentally-mediated POLS differences across two genetically distinct groups of in *U. stansburiana* (Araya-Donoso 2024; Raul Araya-Donoso et al. 2025) using a chromosome-level reference genome and annotation, paired with gene expression data collected north and south of the Vizcaíno Desert, a major biogeographic structuring feature for many species (Dolby et al. 2015), at the center of the Baja California peninsula. All side-blotched lizard populations in this region are trimorphic for throat color, mitigating this as a confounding variable, while environmental conditions such as temperature, precipitation seasonality, and vegetation composition vary due to summer monsoons and tropical cyclones that sometimes bring large “off-season” precipitation events (Araya-Donoso et al. 2024; Figure 1C). Northern populations incur strong seasonal variation in temperature between extremes, while the majority of rainfall occurs in the winter. Southern populations encounter higher, more consistent temperatures through the year with precipitation falling during the summer driven by the North American Monsoons (Figure 1C). These climatic differences lead to different environmental dynamics for *U. stansburiana*, which is predominantly annual on the peninsula. Northern environments are seasonally disparate with relatively lower productivity given mismatches between lizard activity in the spring/summer and rainfall, where southern environments are comparatively climatically stable throughout the lifetime of *U. stansburiana* with precipitation coinciding with times when lizard activity is at its peak. These differences in alignment between peak environmental productivity and activity, and climatic consistency which influences growing season length likely impact life history variation relevant to divergence in pace-of-life processes. Seasonality is known to underlie variation in environmentally-mediated SSD and differences in growth in side-blotched lizards (Chelini et al. 2021) and may exacerbate POLS differences, leading to variation in SSD among these regions. Growing season differences between the northern and southern Baja California peninsula areas we sampled may exacerbate POLS differences (Cabezas-Cartes et al. 2018), and lead to variation in SSD among these regions (Chelini et al. 2021).

In this study we present a high-quality chromosome-level reference genome for the common side-blotched lizard (*Uta stansburiana*). This is the first genome assembly for the genus *Uta* and the seventh chromosome-level published phrynosomatid genome, enabling comprehensive comparative genomic analyses. The genome assembly is chromosome-scale (L90 = 8, N50 = 320 Mb), has a diploid number 2n = 34 consistent with the hypothesized ancestral karyotype of the family phrynosomatidae (Leaché and Sites, Jr. 2009), and reveals many microchromosomes consistent with previous karyotyping work (Pennock et al. (1968, 1969); but see Pinto et al. (2023)). We demonstrate the utility of this genome through analysis of RNA-seq data collected from wild lizards in seasonally disparate environments. Our approach explores how life history features (i.e., morphs and sexes) and environmental divergence (with impacts for population density, SSD) associate with the molecular mechanisms that could be relevant to pace-of-life-related trait variation to develop hypotheses about the evolution of POLS within *U. stansburiana*. If POLS-related processes influence these populations, we expect POLS divergence to present as differential expression in genes related to metabolism, immune function, senescence, stress, regulatory/transcription pathways, and osmotic regulation. We then explore whether these transcriptomic patterns are consistent with POLS-related expectations across environmental differences between northern and southern populations on the Baja California peninsula, a site of extensive cross-species divergence (Hämäläinen et al. 2021; Dolby et al. 2022).

## 2. Materials and Methods

### 2.1 Sample collection: Genome

Fresh liver tissue was collected from an adult male *Uta stansburiana nevadensis* collected by Dr. Pete Zani from Wrights Point, Harney County, OR, USA (permit ODFW024-20; Fig. 1a). This site comprises a large population of *U. stansburiana nevadensis* living on a lava flow surrounded by the Malheur wetlands and has been the subject of an 18-year mark-recapture study. *U. stansburiana nevadensis* found at this site are known to live for up to seven years (Zani and Stein 2018). The lizard was euthanized following IACUC protocol (University of California Merced AUP21-0001) and liver tissue was immediately removed. Half of the tissue was frozen and stored in 100% ethanol at −80 C, and the other half was used in immediate DNA extraction. The remains of the individual were formalin-fixed and prepared for museum donation (to be deposited at the Museum of Vertebrate Zoology, DLE3124).

Genomic DNA was extracted using a modified buffered lysis and detergent salting-out protocol from Moore et al (2004) and Macdonald et al (2011). We assessed sample purity using a NanoDrop 1.0 and quantified samples using the Qubit 2.0 high sensitivity kit. The extracted DNA was sent to the University of California Davis DNA Technologies and Expression Analysis core and sequenced using two Pacific Biosciences Sequel II SMRT-cells run in CCS mode. The remaining frozen liver tissue was sent to DNA Zoo (dnazoo.org) for *in situ* Hi-C library preparation and sequencing (Rao et al. 2014; Dudchenko et al. 2017, 2018).

### 2.2 Sample collection: RNA-seq

Adult *U. stansburiana elegans* were sampled by visual encounter survey and lassoing at seven sites on the Baja California peninsula from northern and southern regions (Collecting permit: 13439/19 DGVS Mexico, IACUC Protocol Arizona State University 20-1737R; IACUC Protocol University of Arizona 20-627 PHS-CDC import permit 20220221-0651A; FWS import # 2022CX2908360; Figure 1b) known to be genetically distinct lineages (Upton and Murphy 1997; Hollingsworth 1999). Sampling was undertaken at the same set of northern and southern sites in the fall and spring during daylight hours. These sampling sites are in the ranges of two reciprocally monophyletic groups located North and South of the center of the Peninsula (Upton and Murphy 1997; Hollingsworth 1999; Araya-Donoso et al. 2024). Capture locations were flagged with tape and morphological measurements were taken of the individuals, including body size (snout-vent length, SVL; Table S1), and throat color photos. Tail tissue samples were collected by autotomization and preserved in RNAlater on wet ice; tail tissue can be sampled with minimal-to-no harm to individuals and is vascularized. As such, it is suitable for the scope of these questions because of the wide array of RNAs found in even small reptilian blood samples (Waits et al. 2020). A broader SVL dataset was used to calculate SSD as described in Chelini et al. (2021) to show that northern sites (mean SSD = 1.04) were sexually monomorphic in body size while southern sites were male-biased (mean SSD = 1.14) and these two regions were significantly different in their degree of SSD (paired *t*-test; df = 2; *t*-stat = −9.04, p _¿_ 0.05). SVL data were collected from three populations in the northern and southern regions during the spring breeding season only, using 2-8 individuals per sex, to estimate average adult male and female SVL. We selected 33 individuals (14 male, 19 female) across seven sites (9 individuals across three sites in the north, 24 individuals across four sites in the south) and roughly equal throat colors (7OO, 4BB, 4YY, 4OB, 1BY, 2OY, 11 unassessable; Sinervo et al. 2001) for RNA sequencing. The full metadata are available in Table S2. Preserved tail tips were sent to the Yale Center for Genome Analysis for RNA isolation, poly-A mRNA library preparation, and sequencing on the Illumina NovaSeq 6000 platform with 150 bp PE reads.

### 2.3 Bioinformatics

#### 2.3.1 Genome assembly

We prepared the draft genome assembly using hifiasm v0.16.1-r375 (Cheng et al. 2021) and dropped k-mers that occurred more than 10 times the homozygous read coverage to improve resolution in repetitive areas, but otherwise assembled the data under default parameters. Hi-C reads were aligned to the hifiasm draft genome assembly using Juicer (Durand et al. 2016). The alignments were then used to scaffold the draft to chromosome-length following Dudchenko et al (2017, 2018) using the 3D-DNA pipeline and Juicebox Assembly Tools. To assess overall depth of the assembled reads, we mapped the filtered and deduplicated reads onto the reference assembly using minimap2 v2.28-r1209 (Li 2018) for long reads and bwa v0.7.17-r1188 (Li 2013) for Hi-C reads. We then calculated average read depth per 100 kb window using bedtools v2.27.1 (Quinlan and Hall 2010).

#### 2.3.2 Genome annotation

We generated an annotation for the chromosome-length genome assembly of *U. stansburiana* as follows. We renamed all HiC-scaffolds to simply “scaffold,” and sorted the chromosome-length scaffolds by decreasing length resulting in shuffling HiC-scaffold_14 through HiC-scaffold_17 (for a full list connecting HiC-scaffold names to the renamed scaffold names, see Table S13). On the genome assembly, repeats were identified with RepeatModeler v2.0.1 (Smit et al. 2015a) and soft-masked with RepeatMasker v4.1.1 (Smit et al. 2015b) Then, we ran multiple iterations of Maker v3.01.03 (Campbell et al. 2014) to annotate the genome. A first round of Maker was run to map and align evidence, which included the transcripts and protein-coding sequences from the phrynosomatids *Urosaurus nigricaudus* (Davalos-Dehullu et al. 2023) and *Phrynosoma platyrhinos* (Koochekian et al. 2022). Then, we ran two rounds of *ab initio* gene model prediction using SNAP v2006-07-28 (Korf 2004) and Augustus v3.4.0 (Stanke et al. 2006). After each round of Maker, we recorded the Annotation Edit Distance (AED) and assessed the annotation completeness by using BUSCO v5.4.2 (Simão et al. 2015) on the predicted transcripts obtained from Maker using the eukaryote and sauropsid gene datasets.

#### 2.3.3 RNA-seq bioinformatics and differential expression analysis

The RNA-seq data were assembled, quality-assessed, trimmed, aligned, and quantified under the nf-core/rnaseq pipeline v3.14.0 (Patel et al. 2023), one of the nf-core workflows (Ewels et al. 2020) which uses reproducible software from Bioconda (Grüning et al. 2018) and BioContainers (Da Veiga Leprevost et al. 2017) projects. We ran this pipeline using Trim Galore v0.6.7 (Krueger et al. 2021) to remove reads below 20 bp or with Q-values below 35, STAR v2.7.9a (Dobin et al. 2013) to align reads to the reference genome, and quantified reads using salmon v1.10.1 (Patro et al. 2017) under default parameters. All other processes in the pipeline were run under default parameters.

We used the nf-core/rnaseq output files and tximport v1.32.0 (Soneson et al. 2015) to prepare a matrix of non-normalized read counts for DESeq2 v1.44.0 (Love et al. 2014). We conducted two different differential expression models: 1) a group-means parameterization of the binary variables region (north and south), sex (male and female), and season (spring and fall) with the categories north, female, and fall set as reference levels for contrasts, and 2) a group-means parameterization of putatively homozygous throat colors (Sinervo et al. 2000, 2001) against the other homozygous throat colors without respect to other variables. We extracted contrasts for region (controlling for season and sex), sex (controlling for sex and region), orange throats (controlling for blue and yellow throats), blue throats (controlling for orange and yellow throats), and yellow throats (controlling for orange and blue throats). We calculated log2-fold changes between significantly differentially expressed genes (Bonferroni-corrected p < 0.025 because we tested two differential expression models).

### 2.4 Gene Ontology term analyses and functional enrichment

We used a two-pronged approach to determine which biological functions were associated with high levels of differential expression between northern and southern populations, controlling for season. First, we used g:Profiler v.e111_eg58_p18_f463989d (Raudvere et al. 2019) on the full list of differentially-expressed genes to determine major enriched biological processes, KEGG pathways, and transcription factors (Tables S3-S7). For this analysis, we used *Mus musculus* as reference because of its completeness in the database. We then visualized enriched biological processes via semantic clustering as a treemap using REVIGO v.1.8.1 (Supek et al. 2011; Fig. S6, S7).

To supplement these results, we used STRING Interaction Database (Szklarczyk et al. 2023) to generate a network to visualize known interactions among the final gene list. Because of the completeness of its functional annotation, we used human as the reference, and we used medium confidence (0.4) as the minimum threshold. We then analyzed the gene networks in Cytoscape (Shannon et al. 2003) to identify subnetworks and “hub” genes with high degrees of connectivity.

### 2.5 Synteny evolution in Phrynosomatidae

We investigated synteny of protein-coding genes in *U. stansburiana* by comparative analysis with other available phrynosomatid chromosome-level assemblies. We compared *U. stansburiana* against *Phrynosoma platyrhinos* (Koochekian et al. 2022), *Sceloporus undulatus* (Westfall et al. 2021), *Urosaurus nigricaudus* (Davalos-Dehullu et al. 2023) and the dactyloid lizard *Anolis carolinensis* (Sampson et al. 2025; Gambón-Deza 2023) using GENESPACE v1.3.1 (Lovell et al. 2022). Because the *A. carolinensis* assembly did not explicitly include microchromosomes, we considered synteny across all scaffolds in all assemblies if they contained syntenic blocks with a minimum of five hits.

## 3. Results

### 3.1 Genome assembly and annotation

PacBio HiFi long read sequencing resulted in 55.9 Gb of consensus sequences assembled into 204 contigs using hifiasm. The contig N50 was 93 Mb, and the longest contig was 170 Mb. The Hi-C guided genome assembly for *U. stansburiana* (Table 1) was 2.10 Gb total length spanning 17 chromosome-length scaffolds (N50: 320 Mb, L90: 8) and high completeness (BUSCO eukaryotic completeness of 99.6%). The chromosome-length scaffolds in the Hi-C-guided assembly were arranged in descending order except for scaffold_14 (for associated HiC_scaffold numbers, see Table S13) likely representing the sex chromosome based on a pseudoautosomal-compatible region. The following scaffold, scaffold_18, is potentially a Y chromosome for the same reason and further analysis is required to definitively identify these as sex chromosomes. Consistent with previous karyotype analysis (Pennock et al. 1968), the final assembly comprises six macrochromosomes ranging from 426 Mb to 146 Mb and 11 microchromosomes ranging from 38 Mb to 16 Mb (Fig. S8, S9). Roughly 50% of the genome corresponded to repeat elements, comparable to the other phrynosomatids (Table 2). A total of 20,350 protein-coding genes were annotated with an average gene length of 20,561 bp, corresponding to a 92.6% completeness of all eukaryote BUSCO (either complete or fragmented; Table 2). Of the 49.29% of the genome that comprises repeat elements (Table 3), the most abundant were unclassified (31.17%) followed by LINEs (15.44%). Over 2/3 of the LINEs (10.26% of the genome) belonged to the L2/CR1/Rex families (Table 3). RepeatModeler did not detect any SINEs, which may account for the relatively high (31.71%) proportion of unclassified repeats.

**Table 1.** Genome assembly statistics for *U. stansburiana* with three other phrynosomatid reference genomes for comparison.

|  | Species |  |  |  |
| --- | --- | --- | --- | --- |
|  | *Uta stansburiana* | *Urosaurus nigricaudus* | *Sceloporus undulatus* | *Phrynosoma platyrhinos* |
| **Genome version** | UCM_Usta_1 | ASU_Unig_1 | SceUnd_v1.1 | MUOH_PhPlat_1.1 |
| **Assembly length (Gbp)** | 2.1 | 1.87 | 1.91 | 1.9 |
| **N50 (Mbp)** | 320 | 279 | 276 | 273 |
| **L50 (n°)** | 3 | 3 | 3 | 3 |
| **Largest scaffold (bp)** | 426,646,591 | 380,405,096 |  |  |
| **Number scaffolds** | 104 | 327 |  |  |
| **Eukaryote BUSCO Assembly (%)** | 99.6 | 99.2 | 99.6 | 98.8 |

**Table 2.** Genome annotation statistics for *U. stansburiana* with three other phrynosomatid reference genomes for comparison.

|  | **Species** |  |  |  |
| --- | --- | --- | --- | --- |
|  | *Uta stansburiana* | *Urosaurus nigricaudus* | *Sceloporus undulatus* | *Phrynosoma platyrhinos* |
| **Genome version** | UCM_Usta_1 | ASU_Unig_1 | SceUnd_v1.1 | MUOH_PhPlat_1.1 |
| **% Repeats** | 49.23 | 45.42 | 41.33 | 44.45 |
| **N° genes** | 20,350 | 17,902 | 15,472 | 17,864 |
| **Average gene length (bp)** | 20,560.80 | 18,191.50 | 22,998.8 | 21,769.8 |
| **Eukaryote BUSCO Annotation (%)** | C + F: 92.6 | C + F: 94.9 | C + F: 96.1 | C + F: 92.5 |
| **Reference** | Dudchenko et al., 2017; Dudchenko et al., 2018; This study | Dávalos-Dehullu et al., 2023 | Westfall et al. 2021 | Koochekian et al., 2022 |

**Table 3.**
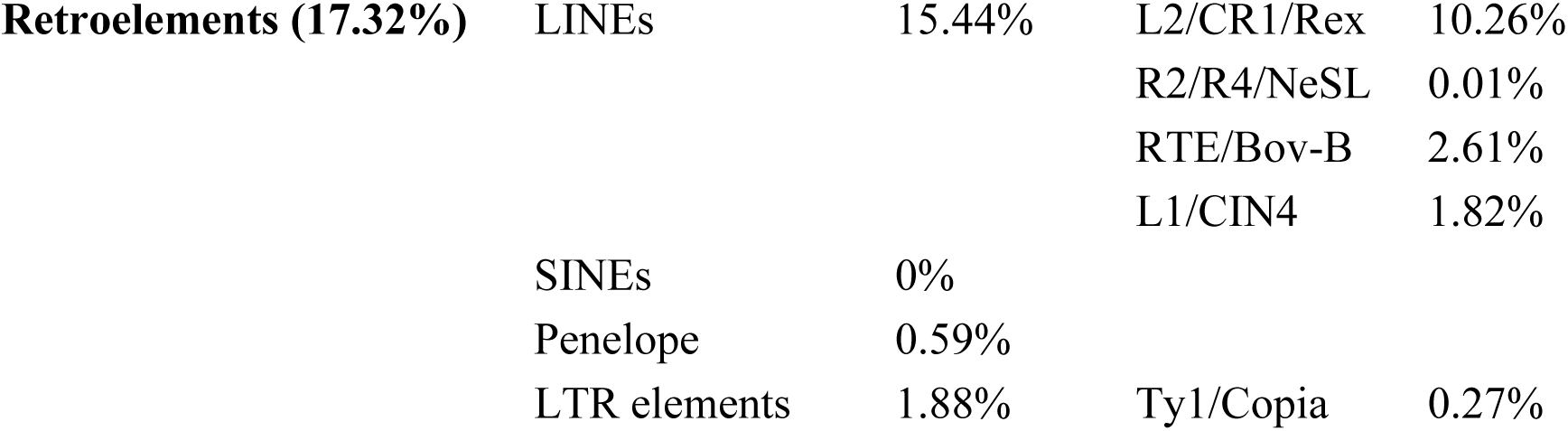
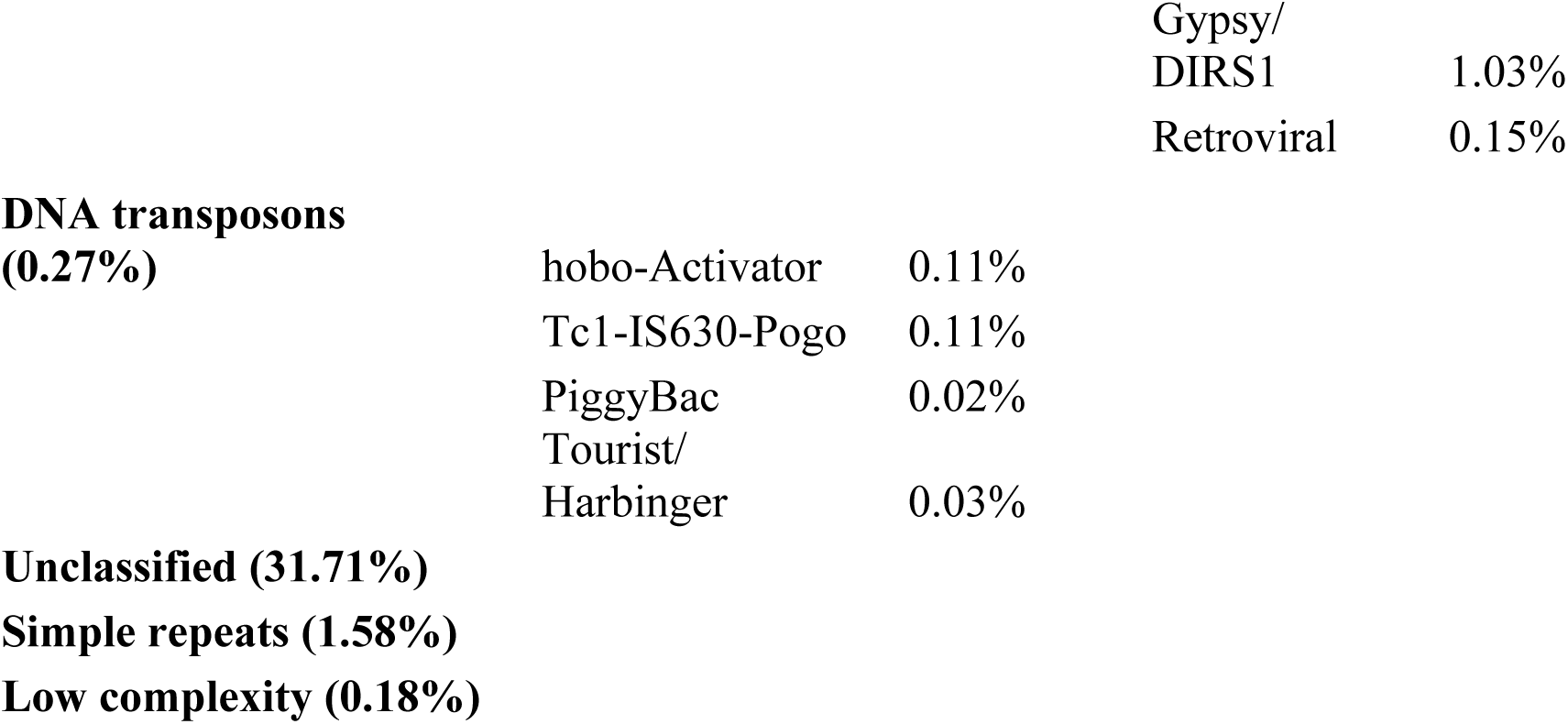
Repeat element statistics for *U. stansburiana* genome from RepeatModeler v2.0.1 (Smit et al., 2015a) and RepeatMasker (Smit et al., 2015b). Total interspersed repeats is 49.29%. Percentages are relative to the total genome length.

|  |  |  |  |  |
| --- | --- | --- | --- | --- |
| **Retroelements (17.32%)** | LINES | 15.44% | L2/CR1/Rex | 10.26% |
|  |  |  | R2/R4/NeSL | 0.01% |
|  |  |  | RTE/Bov-B | 2.61% |
|  |  |  | L1/CIN4 | 1.82% |
|  | SINEs | 0% |  |  |
|  | Penelope | 0.59% |  |  |
|  | LTR elements | 1.88% | Ty1/Copia | 0.27% |

|  |  |
| --- | --- |
| Gypsy/<br>DIRS1 | 1.03% |
| Retroviral | 0.15% |

**DNA transposons (0.27%)**
|  |  |
| --- | --- |
| hobo-Activator | 0.11% |
| Tc1-IS630-Pogo | 0.11% |
| PiggyBac | 0.02% |
| Tourist/<br>Harbinger | 0.03% |

After filtering and deduplication, the mean read depth of the assembly-generating reads was 36.92x (median 36.80x). Scaffold_14 had a mean read depth of 22.44x (median 20.67x), approximately half of that of other chromosome-length scaffolds, consistent with its tentative identification as a heterogametic sex chromosome (Fig S13, Table S14).

### 3.2 Synteny evolution in phrynosomatid lizards

In comparing genomic synteny in *U. stansburiana* to other phrynosomatids, we found a high level of synteny conservation in the six macrochromosomes with some short, inverted regions (Figure 2). We identified several rearrangements in the microchromosomes, especially scaffold_8, scaffold_12, scaffold_13, scaffold_15, scaffold_16, and scaffold_17. The likely X chromosome, scaffold_14 showed substantial conserved synteny with other identified X chromosomes (scaffold 14 in *Urosaurus nigricaudus*, scaffold 10 in *Sceloporus undulatus*, scaffold 9 in *Phrynosoma platyrhinos*, and the X chromosome in *Anolis carolinensis*), again suggesting that this scaffold represents the X chromosome in *U. stansburiana*.

**Figure 2.**
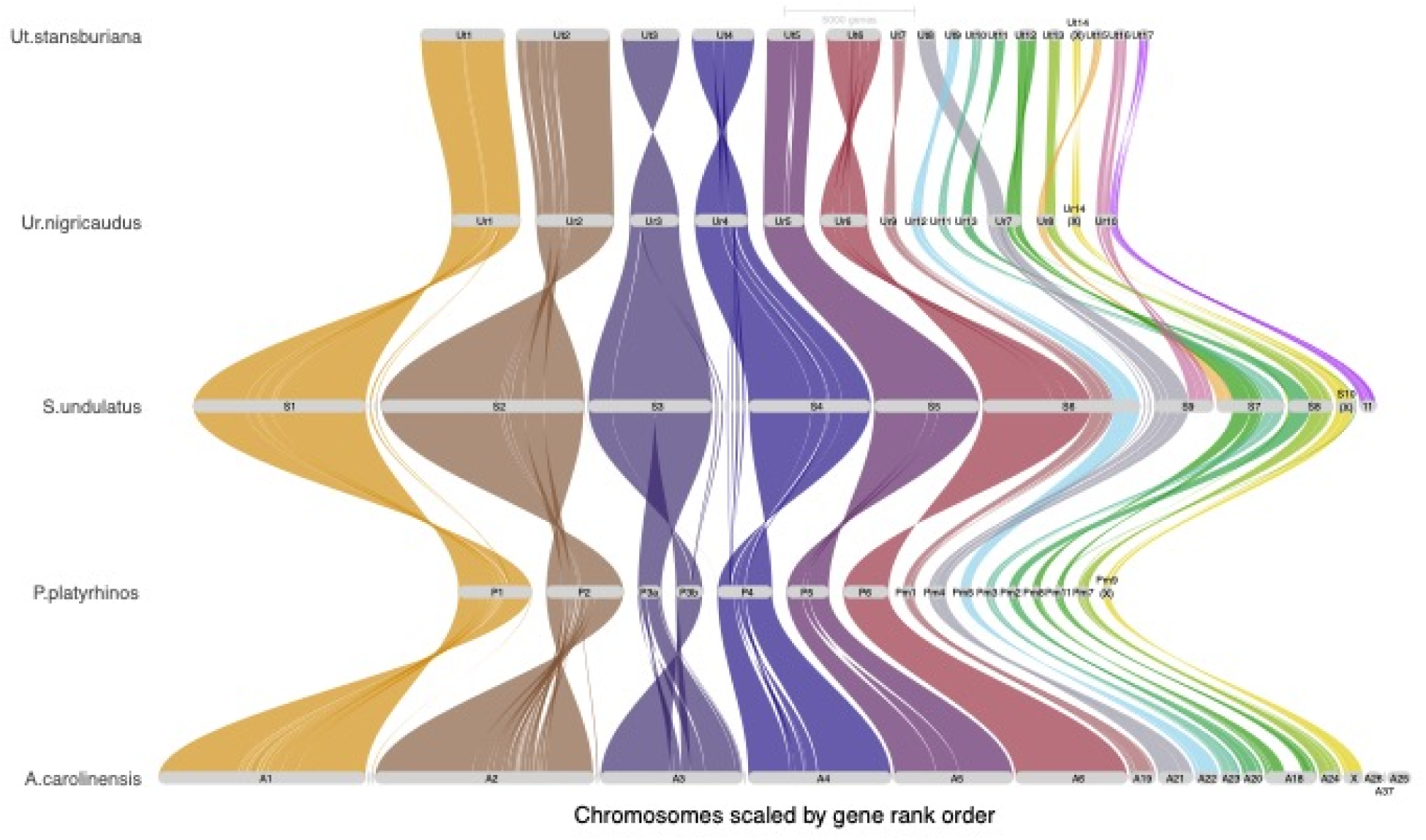
Riparian plot of genomic syntenic blocks in four phrynosomatid reference genomes (*Phrynosoma platirhinos*, *Sceloporus undulatus*, *Urosaurus nigricaudus*, and *Uta stansburiana*) and one dactyloid (*Anolis carolinensis*) reference genome. Chromosome-level scaffolds have been re-named to reflect the position of each scaffold when sorted by length and appended with the first letter or first two letters of the genus. Suspected or identified X chromosomes are noted.

### 3.3 RNA-seq bioinformatics

RNA sequencing resulted in 2.4 billion 101 bp paired-end reads, ranging from 24.2 million to 46.8 million raw reads per sample. 73.5% of raw reads had Q-scores of 35 or higher, while 92.8% of raw reads had Q-scores of 30 or higher. Raw read GC content ranged from 40-45% per sample. After trimming and filtering, average sequence length was 92 bp, ranged from 20-100 bp, and resulted in 2.35 billion paired-end reads. We retained 35.6 million paired-end reads per sample on average. Of these, 74.9% of read-pairs uniquely mapped to the genome and 2.7% multiply mapped; only uniquely mapped transcripts were retained. The RNA-seq data comprised reads mapping to 19,970 genes (i.e., those with non-zero counts) in total but averaged 14,830.5 genes per sample and ranged from 12,823 to 16,317 annotated genes per sample.

### 3.4 Differential expression by sex

DESeq2 analysis yielded 9 genes differentially expressed (Bonferroni-corrected adjusted p-value <0.025, Table S3) between female (reference level) and male lizards controlling for region and season, all of which had functional gene annotations. Moderated log2-fold change values for statistically significant gene annotations ranged from 1.50 (*gstm4*) to −1.40 (*shfl*). G:Profiler returned two enriched biological process GO terms (Table S8): *long-chain fatty acid metabolic process* (*p* = 1.51 x 10^-2^), and *xenobiotic catabolic process* (*p* = 2.19 x 10^-2^). Six enriched KEGG terms were returned: *drug metabolism cytochrome P450* (*p* = 2.92 x 10^-2^), *glutathione metabolism* (*p* = 3.00 x 10^-2^), *metabolism of xenobiotics by cytochrome P450* (*p* = 3.17 x 10^-2^), *platinum drug resistance* (*p* = 3.70 x 10^-2^), *chemical carcinogenesis-DNA adducts* (*p* = 4.08 x 10^-2^), and *drug metabolism-other enzymes* (*p* = 4.69 x 10^-2^). The STRING interaction network contained only a single edge, between *gstm4* and *gstm1*, did not contain significantly more interactions than expected (p = 0.332), and presented one additional enriched biological process (*nitrobenzene metabolic process,* FDR = 0.04) and one additional protein domain and feature (*glutathione s-transferase, Mu class*, FDR = 0.03).

### 3.5 Differential expression by throat color

DESeq2 analysis yielded one statistically significantly differentially expressed gene in orange-throated lizards compared to blue- or yellow-throated lizards (Bonferroni-corrected adjusted p-value 0.025), which was associated with an annotation (*slc12a7*; Table S4); this had a large moderated log2-fold change value of 7.21. g:Profiler returned a single enriched KEGG term, *collecting duct acid secretion* (*p* = 0.05). Because there was a single enriched gene, further exploration of GO terms was not pursued. Comparison between blue throated lizards and yellow- and orange-throated lizards found nine differentially expressed genes, all of which had functional annotations. Log2-fold change values for statistically significant annotated genes ranged from −4.80 (*phpt*) to −6.73 x 10^-7^ (*adam20*). g:Profiler did not return any biological process terms but found one enriched KEGG term (Table S10): *GnRH signaling pathway* (*p* = 4.68 x 10^-2^). The STRING network did not contain any edges and did not yield any significant enrichments. Finally, the comparison between yellow-throated and blue- and orange-throated lizards yielded two differentially expressed genes both associated with annotations. Log2-fold change values ranged from 2.78 (*plcd1*) to 3.88 (*cd74*). g:Profiler returned one enriched biological process term: *positive regulation of small molecule metabolic process* (*p* = 4.98 x 10^-2^) and no enriched KEGG processes.

### 3.6 Differential expression by geographic region, controlling for sex and season

DESeq2 analysis revealed 163 annotated genes differentially expressed (Bonferroni-corrected adjusted p-value < 0.025, Figure 3) between southern and northern lizards accounting for sex and season. Log2-fold change values for statistically significant annotated genes ranged from 11.43 (*adam20*) to −9.40 (*taar1*) (Figure 4, Table S7). g:Profiler results returned two statistically enriched biological processes: *regulation of response to stimulus* (*p* = 7.51 x 10^-3^) and *regulation of multicellular organism process* (*p* = 2.90 x 10^-2^) (Table S12, Fig. S6). There were four enriched KEGG terms: *endocrine resistance* (*p* = 6.03 x 10^-3^), *parathyroid hormone synthesis, secretion, and action* (*p* = 1.38 x 10^-3^), *chemical carcinogenesis-receptor activation* (*p* = 2.47 x 10^-2^), and *hepatocellular carcinoma* (*p* = 2.88 x 10^-2^). The STRING interaction network had 153 nodes, an average clustering coefficient of 0.35, and the network did not have significantly more interactions than expected by chance (*p* = 0.09). Network analysis yielded four hub genes with ≥ 10 connected edges: *egfr* (18), *ccnd1* (14), *fos* (14), and *kras* (10).

**Figure 3.**
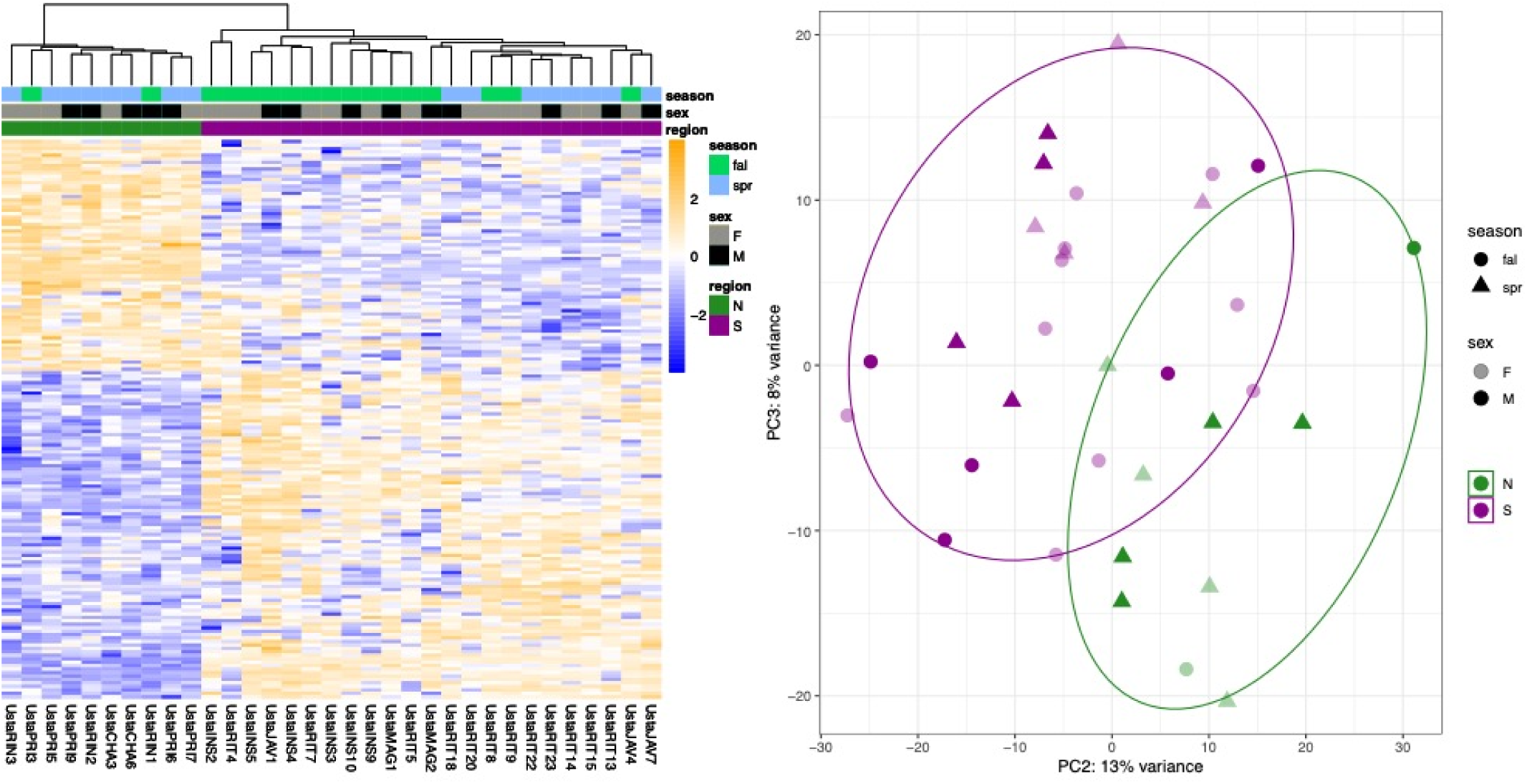
**A)** Heatmap of log2-transformed count data for genes with differential expression by region, controlling for season and sex. Samples are labeled by region, season, and sex. **B)** Principal components analysis of regular log-transformed count data. PC2 and PC3 are shown here to better illustrate the distinctions between groups; see Fig S4 for PC1 and PC2. Ellipses enclose the centroids of the northern and southern samples.

**Figure 4.**
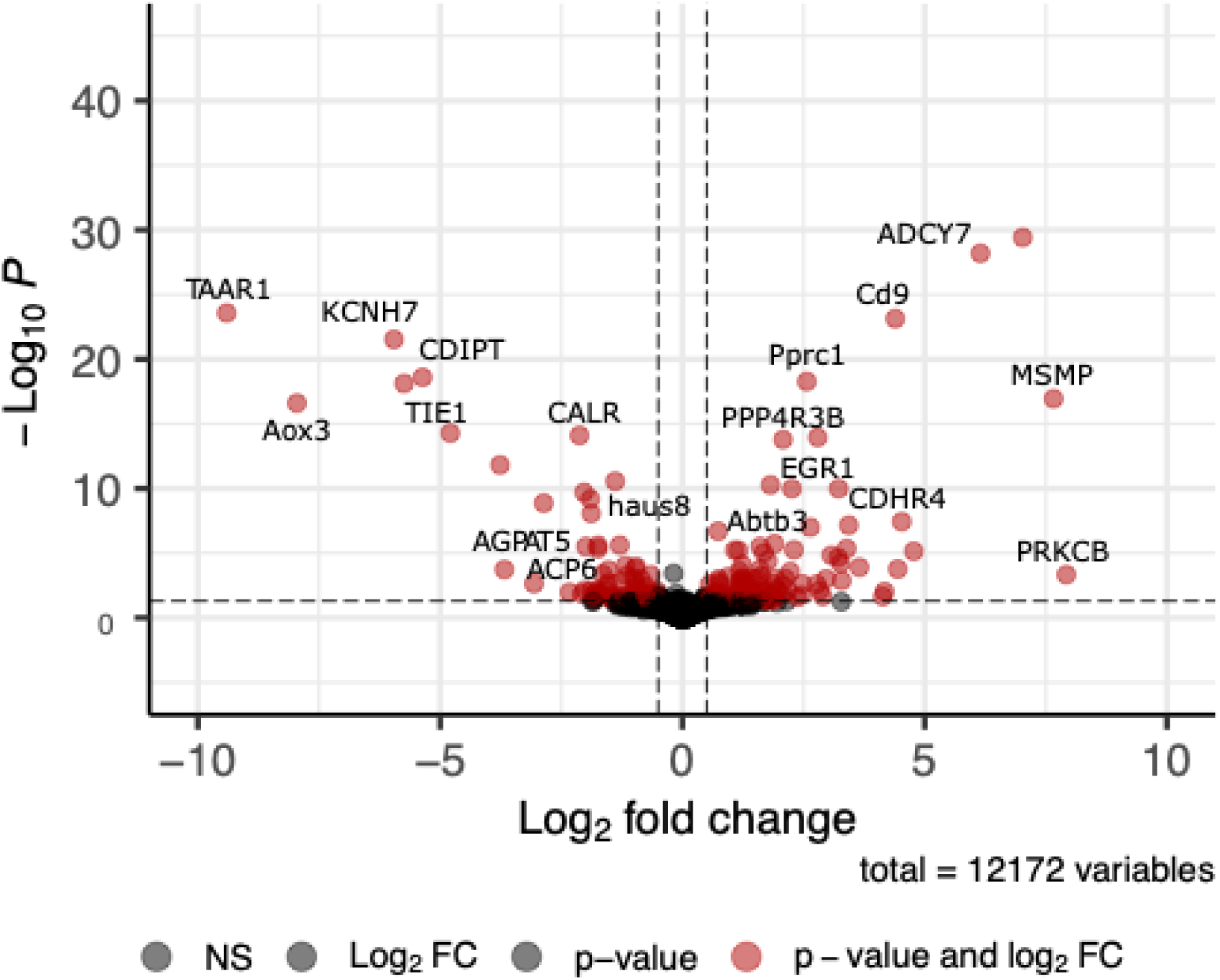
Volcano plot of genes differentially expressed between northern and southern groups, controlling for seasonal differences. Positive log2 fold change values correspond to genes differentially over-expressed in the southern group, while negative log2 fold change values correspond to genes differentially under-expressed in the southern group. ADAM20 is not plotted here, but see Fig. S5.

#### 3.6.1 Endocrine pathways

Two of the four enriched KEGG pathways in the region comparison were related to endocrine signaling, specifically *endocrine resistance* (hsa01522), *parathyroid synthesis, secretion, action* (hsa04928). Eight enriched genes were associated with this pathway in the STRING interaction network: *prkcb* (log2-fold change 7.93), *adcy7* (log2-fold change 6.15), *egr1* (log2-fold change 3.22), *egfr* (log2-fold change 1.19), *fos* (log2-fold change −1.59), and *casr* (log2-fold change −1.67). Endocrine resistance pathways (hsa01522) affect cell cycle and apoptosis; seven enriched genes are associated with this pathway in the STRING interaction network: *adcy7* (log2-fold change 6.15), *egfr* (log2-fold change 1.19), *dll1* (log2-fold change 1.17), *kras* (log2-fold change −0.65), *ccnd1* (log2-fold change −0.097), *fos* (log2-fold change −1.59), and *ncor1* (log2-fold change −1.88); g:Profiler did not identify *kras* as enriched for this pathway. Endocrine resistance and secretion are components of metabolism, cell proliferation, and sexual maturation which are predicted to occur with POLS differences.

#### 3.6.2 Proto-oncogenes and cancer pathways

One of the four enriched KEGG pathways (*hepatocellular carcinoma*) and a major keyword (proto-oncogene) in the analysis between regions were related to cancer or proto-oncogenes. The hepatocellular carcinoma KEGG pathway (hsa05225) describes several key signaling processes (genetic and epigenetic) that occur throughout liver cancer related to chromatin remodeling and oxidative stress. Nine enriched genes are associated with this pathway: *prkcb* (log2-fold change 9.60), *prkca* (log2-fold change 1.16), *smarcd3* (log2-fold change 1.66), *egfr* (log2-fold change 1.34), *kras* (log2-fold change −0.71), *ccnd1* (log2-fold change −1.07), *gsto1* (log2-fold change −1.22), *mgst1* (log2-fold change −1.25), and *dvl1* (log2-fold change −2.18). Chemical carcinogenesis-receptor activation (hsa05207) was identified as an enriched KEGG pathway in g:Profiler and describes cell signaling pathways which may control the pace at which cells replicate. g:Profiler identified eight enriched genes associated with this pathway: *fos* (log2-fold change −1.59), *gsto1* (log2-fold change −1.22), *mgst1* (log2-fold change −1.25), *ccnd1* (log2-fold change −1.07), *dll1* (log2-fold change 1.17), *egfr* (log2-fold change 1.34), *adcy7* (log2-fold change 6.15), and *prkcb* (log2-fold change 9.60).

## 4. Discussion

Pace-of-life traits may be influenced by environmental processes that are likely to impact sexual selection dynamics, yet little is known about the functional molecular mechanisms giving rise to such variation within or between species. Side-blotched lizards are an emerging model system exhibiting variation in several key traits that are known to predict POLS divergence; variation in SSD, longevity, polymorphism, population density, sexual signaling, immune response and growth, and other traits are all known to vary among populations experiencing alternate environmental conditions. Our study provides a high-quality annotated reference genome for *Uta stansburiana* as a resource for into the processes which underlie the variation across this widely distributed species and a template for studying the molecular mechanisms underlying POLS variation in other species. Our results identify pathways consistent with potential POLS divergence associated with differences in environmental stability across the lifetime and identify brain development and function, cell signaling, fertility, immunity, homeostasis, metal ion binding, and metabolism as candidate mechanisms underlying POLS variation.

### 4.1 Genome features and comparative genomics

This is the first genome assembly for the genus *Uta*, and the seventh chromosome-level published phrynosomatid genome, enabling comprehensive comparative genomic analyses. The genome assembly is chromosome-scale (L90 = 8, N50 = 320 Mb), and reveals a large number of microchromosomes (first described as such by (Pennock et al. (1968, 1969); but see Pinto et al. (2023)) implying substantial chromosome fission or fusion events relative to closely related lizards. This is consistent with the hypothesized ancestral karyotype of 2n = 34 for phrynosomatidae (Leaché and Sites, Jr. 2009). We identified the 17 longest scaffolds as likely chromosomes of *U. stansburiana nevadensis*. These comprise six macrochromosomes and 11 microchromosomes with a diploid count of 2n = 34. This is consistent with previous karyotype analysis in *Uta sp.* (Pennock et al. 1968) and the hypothesized ancestral karyotype of all phrynosomatids (2n = 34; Leaché and Sites, Jr. 2009). Further analysis is needed to confirm the identity of the sex chromosomes, but preliminary observations of pseudoautosomal regions suggest that scaffold_14 is the X chromosome, consistent with synteny analysis and read depth estimation, and that scaffold_18 is the Y.

At the macrochromosome level, the *U. stansburiana* genome shows high synteny and small regions of inversions relative to the other phrynosomatid genomes, consistent with previous findings in Iguania (Davalos-Dehullu et al. 2023). The microchromosomes also exhibit highly conserved syntenic blocks between these lizards, however, they also show clear evidence of substantial chromosome fission/fusion among phrynosomatids (Figure 4). In particular, scaffold_8 and scaffold_12 contain syntenic blocks that have undergone substantial fission/fusion. Finally, our hypothesized X chromosome (scaffold_14) was highly syntenic with identified X chromosomes in other phrynosomatids, including the closely related *Urosaurus nigricaudus*.

In general, microchromosomes contain conserved syntenic blocks of slowly evolving genes while undergoing dynamic fission/fusion (Srikulnath et al. 2021), and the consequences of these microchromosomes and their fission or fusion depend on their content. Increasing evidence shows that microchromosomes house genes under selection (Schield et al. 2019) and that the syntenic lability provided by microchromosomes could allow for flexibility in local adaptation via unique linkage disequilibrium regimes (Perry et al. 2021). This could contribute to plasticity and standing genetic variation for local adaptation in side-blotched lizards across their large range and highly disparate abiotic environments. However, previous karyotyping work in the genus *Uta* suggests that this may not be a hard rule; Pennock et al. (1968, 1969) found the same diploid number of chromosomes in all *Uta* species analyzed including insular taxa which presumably have very distinct local adaptations and recombination regimes. The consequences of microchromosome fission/fusion in side-blotched lizards relative to other phrynosomatid lizards will depend on the genic content of the microchromosomes.

### 4.2 Sexual differentiation in gene expression and POLS

While there are limited reasons to expect that male and female lizards would experience alternate POLS, many studies have detected large sexual differences in gene expression in other taxa (e.g., Lipshutz et al. 2025). Our gene expression data allowed us to assess the relative contribution of sex, throat color polymorphism, and environmental divergence among regions relative to predictions under the POLS hypothesis. Results demonstrate minimal differential expression between male and female *U. stansburiana*. Among differentially expressed genes, the main enriched processes deal with glutathione metabolism with associated genes having moderated log2-fold change values between 1.04 (*gstm1*) and 1.50 (*gstm4*). Our results indicate that there may be limited mechanisms for sex-specific POLS in this species. It is noteworthy to mention that other iguanian reptiles (*Anolis carolinensis,* Marin et al. 2017; *Sceloporus malachitichus,* Lisachov et al. 2021; the closely-related *Urosaurus nigricaudus,* Davalos-Dehullu et al. 2023) show evidence of dosage compensation systems, where males overexpress their single copy of the X chromosome to achieve expression levels of female XX. If present in side-blotched lizards, this could contribute to the low number of sex-biased expression observed. This also could be because genes expressed differentially by sex could vary among tissue type (Oliva et al. 2020) and we did not sample gonad tissue (Xu et al. 2020). However, *U. stansburiana* are predominantly sexually monomorphic for size across their range (Chelini et al. 2021) even while both sexes are polymorphic in throat color, suggesting limited sex differences. Finally, our tail samples were vascularized and included blood which allowed for detection of hormonal and reproductive gene expression differences in other comparisons.

### 4.3 Differential gene expression, polymorphism, and POLS

There have been some suggestions that alternate reproductive tactics may lead to divergence in POLS between morphs in polymorphic species (e.g., Friesen et al. 2017). Contrasts among throat colors, indicative of alternative reproductive tactics, returned few DEGs, few enriched GO terms, and no STRING interactions. The only DEG in orange-throated lizards was *slc12a7*, which produces a solute carrier protein maintaining renal homeostasis in the presence of potassium and chloride ions in humans. Relative to blue-throated lizards, orange- and yellow-throated lizards over-expressed genes related to gonadotropin releasing hormone pathways, however, moderated log2-fold change values for these DEGs were near zero. Blue-throated lizards did not over-express any genes relative to the other morphs, and the two DEGs with high moderated log2-fold change values (*phpt1*, a gene producing proteins involved in histidine phosphorylation, and *ppp4r3b*, which is involved in carbohydrate metabolism and DNA repair). Finally, yellow-throated lizards over-expressed genes involved in small molecule metabolic processes; *cd74* is a component of the major histocompatibility complex (MHC) and plays a role in regulation of prostaglandins (regulators of inflammation and fever response), while *plcd1* is involved in inositol triphosphate synthesis which mobilizes calcium ions from cellular stores.

Such few DEGs among color morphs could be explained by the fact that we did not sample tissue which might reveal differences in epidermal pigment or structural color. Previous studies have suggested that there may be difference among morphs within *U. stansburiana* (Hazard et al. 2019) and other polymorphic lizards (Friesen et al. 2017) in physiological traits associated with POLS variation. We find limited evidence of that, since only blue and yellow morphs show variation in one or two molecular pathways associated with metabolism, aging, and/or immunity. We note that the throat color analysis did not include the correction for region, season, or sex, meaning that while this should be a morph-specific expression signal, it could be driven by uneven sample sizes across other variables or small sample sizes within variables.

### 4.4 Environmental influences on gene expression and POLS

Our analyses revealed the environmentally divergent north and south is as a predominant force in differential expression of these lizards. We find significant differences in POLS-related gene pathways associated with brain development and cognition, temporal regulation, signaling, homeostasis, genome stability, sperm production, immune function, and ion binding. Southern populations overexpressed (log2-fold change > 2) eight genes related to brain development and function. These genes were associated with neuron maturation (*fbxo41*), aggregation (*sspo*), synapse assembly (*iqsec2*), synapse function (*slc9a6*, *chrdl1*, *fbxo41*), and post-synaptic organization (*arhgap39*) this is consistent with genes under differential selection in Araya-Donoso et al. (2024). Genes related to dopamine receptor binding (*clic6*) and GABA-based neurotransmitter signaling (*gabra5*) implicated in learning and behavioral fear responses were also enriched (Tan et al. 2011; Syding et al. 2023). In contrast, northern populations over-expressed (log2-fold change < −2) two related genes. These genes were related to neurotransmitter response and release (*kcnh7*) or reception (*taar1*) rather than neurogenesis.

Side-blotched lizards in populations with low predation and high sexual signaling intensity tend to have larger ventral posterior amygdalae (LaDage et al. 2022) and increased aggression and spatial processing needs, including in more complex spatial settings. This results in increased neurogenesis (LaDage et al. 2013, 2016). Our data indicate that southern region populations show male-biased SSD which is consistent with faster male growth and longer growing seasons (Chelini et al. 2021) which could lead to increased reproductive rates. Side-blotched lizard populations with increased reproductive rates often have increased rates of predation (Parker and Pianka 1975). Over-expression of genes related to neurogenesis in the southern populations could be an indication that southern lizards are signaling at higher rates in spite of predation risk consistent with a faster POLS and is also consistent with observations that most adult male lizards are removed from populations in southern portions of the species’ range by predators within their first breeding (Edwards D, pers. obs.).

Despite evidence for increased neurogenesis in side-blotched lizards, there exists mixed evidence for cognitive performance as a component of POLS. Goulet et al. (2018) found that skinks with fast POLS were more exploratory but slower learners, while De Meester et al. (2022) found no link between cognitive ability and ecology in lacertids. This could be because increased neurogenesis in side-blotched lizards is related to social behavior rather than memory and learning (e.g., Mettke-Hofmann 2014). This is supported by the over-expression of *gabra5* in southern populations. *gabra5* is both implicated in elevated stress responses and decreased learning in knockout studies and decreased short-term memory when up-regulated with drug treatment in mice (Tan et al. 2011; Syding et al. 2023). Over-expression of genes related to neurogenesis in southern populations is consistent with a fast POLS and male-biased SSD, but these results should be validated with careful assessment of population density, sexual signaling intensity, and cognitive ability in these populations.

Southern region populations of *U. stansburiana* showed higher expression (moderated lfc > 2) of genes related to transcriptional regulation (*egr1*, *prkcb*, and *pprc1*), genome repair (*egr1*, *ppp4r3b*, and *hmbox1*), and cellular and hormonal signaling especially in homeostasis (*prkcb*, *pprc1*, *aoc1*, and *glp2r*). Many of these genes have multiple roles in these functions and may be thematically linked by upregulation of *egr1*. *Egr1* is a multi-functional gene which regulates the biosynthesis of luteinizing hormone and mediates metabolism, reproduction, cell cycle regulation and DNA damage repair. Network analysis did not identify it as a major hub gene given our specified threshold of node degree > 10, but it was the sixth most connected DEG (seven connections). In northern populations, we found over-expression of four genes related to metabolism and homeostasis (*aox3*, *slc2a9*, *faah*, and *cdipt*). Transcriptional regulation, genome stability, and homeostasis are foundationally associated with a slower pace of life. These patterns may emphasize regulatory complexity and flexibility, consistent with a faster POLS, while northern populations prioritize metabolic stability and homeostasis, consistent with a slower pace-of-life.

Lucas and French (2012) demonstrated POLS shifts in *U. stansburiana* when environmental or urban stressors led to increased stress, decreased life maintenance processes, and increased reproductive output. We identified two enriched KEGG pathways related to endocrine processes and two related to cell cycle regulation associated with differential expression between southern and northern regions. Parathyroid synthesis and activation increases serum calcium and decreases bone density in other iguanid lizards (e.g., *Dipsosaurus dorsalis* and *Sceloporus grammicus*; McWhinnie and Cortelyou 1968). Four of the six genes related to this pathway were over-expressed in southern populations (*egfr*, *egr1*, *adcy7*, and *prkcb*). Parathyroid hormone (PTH) synthesis, secretion, and action (hsa04928) is responsible for regulating calcium and phosphorus ions. In cases of restricted dietary calcium, parathyroid hormone will stimulate bone turnover and prevent loss of phosphate ions in the kidney. In iguanian lizards, injection of PTH causes acute hypercalcemia and sometimes hyperphosphatemia (McWhinnie and Cortelyou 1968) and may elevate in response to metabolic or nutritional challenges and stress. This may reflect a deficit in resource allocation or increase in some stressor in southern populations resulting in quicker use of energy reserves. For example, predation rates might be higher in the south (but see Wilson 1991), higher population density or long periods between typical hurricanes (Parker 1974) could limit resources and increase stress (e.g., Malisch et al. 2020), or simply longer active seasons could result in higher generational turnover rates. Elevated reproduction, high competition for resources, and increased stress are all concordant with a faster POLS.

Cell signaling and proliferation are functional components of growth rates and aging which vary among POLS (Ujvari et al. 2022). The KEGG term *endocrine resistance* refers to cell cycle resistance to hormonal signaling, typically through cancer research in humans (i.e., Ribeiro and Freiman 2014). Three of the six genes related to this pathway were over-expressed in the south (*egfr*, *dll1*, and *adcy7*). Mixed expression of endocrine resistance genes is somewhat non-specific and may be due to many different processes such as decreased need for endocrine resistance, efficient cell proliferation, etc. Understanding which of these processes is responsible for generating this pattern will require detailed understanding of signaling pathways resulting in cell proliferation (or lack thereof), and consequences for POLS will depend on the effects of the DEGs.

Differences in longevity may also be associated with anticancer strategies (Tissier et al. 2022, Ujvari et al. 2022). The two enriched KEGG pathways related to cell cycle regulation were *hepatocellular carcinoma* and *chemical carcinogenesis-receptor activation*. Like *endocrine resistance*, these KEGG terms are primarily known through cancer research in humans. Three of the seven genes involved in the *hepatocellular carcinoma* pathway were over-expressed in the south (*egfr*, *prkcb*, and *smarcd3*). Four genes have functions specific to cell proliferation or carcinogenesis-*smarcd3* is a tumor suppressor gene, *gsto1* and *mgst1* are involved in antioxidant response, and *dvl1* is a cell proliferation regulator. The fact that a tumor suppressor gene is over-expressed in the south while antioxidant and cell proliferation control genes are under-expressed suggests that growth may be continuous and rapid in southern populations and is consistent with other results pointing toward environmental variation influencing differences in POLS between the regions.

The *chemical carcinogenesis-receptor activation* pathway had four of eight genes over-expressed in the south (*egfr*, *dll1*, *adcy7*, and *prkcb*). This KEGG pathway describes several cell receptor pathways which respond to chemical carcinogens to result in proliferation, chemoresistance, and apoptosis. GSTO1 and MGST1 are also involved in this pathway. The role of differential expression of genes related to cell proliferation and cancer pathways in *U. stansburiana* is likely dependent on the function of the individual genes implicated. Specifically, differences in biotic or abiotic conditions likely create alternate selection for POLS, echoing previous evidence for environmental drivers of POLS (e.g., Williams et al. 2010; Hämäläinen et al. 2021).

If, as we predict, southern region populations have higher densities and longer growing and activity seasons, POLS variation in reproductive rates could present as differential expression in molecular control of sexual function and fertility. Four genes over-expressed (moderated lfc > 2) in the southern population were directly related to sperm production or male fertility. *Adam20*, *topaz1*, and *piwil1* are directly involved in sperm maturation and spermatogenesis. *Adam20* is the single-most differentially expressed gene in the dataset (moderated log2 fold-change = 11.43). *Piwil1* which binds and suppresses TE-derived piwi interfering RNAs (piRNAs) to maintain germline integrity and is associated with infertility of certain crosses in canids (Stalker et al. 2016); it is thought to be a highly-conserved gene among amniotes (Lim et al. 2013). Elevated expression of a piRNA gene and genes related to spermatogenesis, especially those often differentially expressed in the testis, suggests that southern lizards have evolved sperm-related gene expression differences relative to northern males after controlling for seasonal effects. Direct analysis of sperm and cross-group sperm-egg fertilization rates would be helpful to resolve the consequences of these gene expression differences.

Northern populations over-expressed two calcium-binding genes, *wdr49* and *calr*, and one magnesium-binding gene, *ttll11*. *Calr* may be involved in oocyte maturation and the prevention of polyspermy, possibly implicating increased investment in offspring rather than total reproductive output. Southern populations expressed four calcium-binding genes, *pamr1*, *zbtb8os*, *cpne4*, and *cdhr4*, one magnesium-binding gene, *adcy7*, and one zinc-binding gene, *otud7b*. While these genes generally have diverse functions, the greater number of calcium ion binding genes expressed in southern lizards suggest that they have elevated stress hormone levels (Park et al. 2017) which can dampen immunocompetence (Hudson et al. 2020) and require physiological compensation. Taken together, these results suggest that northern region populations invest more in reproduction within a limited season, while southern region populations operate under higher levels of stress because of a faster POLS.

Side-blotched lizard populations with higher parasite load tend to have higher immunocompetence (Spence et al. 2017), and ectoparasite load was much higher in the south (G. Dolby, pers. comm.). Side-blotched lizard populations which more heavily invest in reproduction tend to heal wounds faster and have higher immunocompetence at the expense of body condition during the reproductive season (Smith et al. 2019), a sign of a fast POLS. Both northern and southern region populations differentially expressed genes related to immune function and ion binding, but the connections between these categories and among the individual genes was idiosyncratic. Immune genes expressed higher in the north are *zdhhc11* and *clec5a*, while southern region populations over-expressed *adcy7*, *otud7b*, *tmem45b*, and *msmp*. In general, *zdhhc11* and *clec5a* are components of the innate immune system involved in viral infections, while genes over-expressed in the south have a more general immune function. This may reflect that southern lizards are exposed to a greater diversity of pathogens (Clark and Bradford 1969; Quillfeldt et al. 2018), consistent with our hypothesis that side-blotched lizards in the south may be at higher density. These results suggest that northern region lizards are subject to fewer diseases while southern region lizards require a wider variety of immune responses to pathogens possibly due to higher prevalence of disease and a higher population density because of a faster POLS.

## 5. Conclusions

The molecular mechanisms controlling variation in POLS are relatively understudied. We used the high-quality chromosome-level reference genome presented here alongside gene expression data to test if populations living under different environmental conditions show expression patterns consistent with hypotheses relevant to POLS for the first time in a wild reptile. In a system which demonstrates range-wide shifts in longevity, polymorphic throat color and reproductive tactics, we find few genes related to sex- or throat color-based expression differences and limited to no evidence that sex or alternate reproductive tactics contribute to POLS differences. Our results indicate that environmental variation may be a major force driving gene expression differences that are consistent with POLS shifts. We found expression differences related to homeostasis in the face of resource stress, sexual function, and cell proliferation control. This points to variation in environments as mediating growth and reproduction opportunities, thereby affecting population dynamics and driving the evolution of longevity which all influence POLS variation in this emerging model system. These results represent early molecular insights into potential POLS-related processes, and we emphasize that functional validation and ecological data will be important to determine life history relevance. Future research should consider *U. stansburiana* to be a model system for studying the evolution of POLS, especially for understanding the relationships environmental variation and differences in alternative reproductive tactics, growth, reproduction, aging, senescence, telomere length, cancer suppression, and resource allocation.

## 6. Acknowledgments

We thank S. Baty and A. Biddy for help with bioinformatics and analysis, D. Ardell for insight into modeling, P. Zani for providing the specimen from which we generated the genome, B. Carlson and N. Whutthituntisil for feedback on the manuscript, O. Nguyen and L. Froenicke for help with long-read sequencing, K. Kusumi and D. Denardo for helpful discussions, and C. Pedraza-Marrón and M. Dawson for their advice and support.

## 7. Data Availability Statement

The authors affirm that the data necessary to confirm the conclusions of this article are available in the article, figures, table, or are available in public repositories. Raw Hi-C data are available on SRA (BioProject PRJNA512907, BioSample SAMN46443096). An interactive Hi-C contact map for the final assembly is available at https://www.dnazoo.org/post/a-top-notch-blotch, https://www.dnazoo.org/assemblies/uta_stansburiana. The assembled and annotated genome will be available on NCBI, submission number SUB15340363. The raw long-read and RNA-seq data are available on SRA (long-read data are under submission number SUB15364625, and RNA-seq data are under submission number SUB15372950; both are under BioProject PRJNA1268585).

## 8. Funding

This project was funded by NSF EAR #2305608 to G.A.D.; NSF EAR #1925771 to A.M.V.; NSF DBI-2021795, NSF PHY-2210291, and NIH CEGS RM1HG011016-01A1 to E.L.A., R.A.D. was supported by the doctoral fellowship 72200094 (ANID, Chile); E.D-D. was supported from CONAHCYT fellowship 162588; D.L.E. and S.R.F. acknowledge funding support from the University of California, Merced. The Hi-C guided assembly was done in association with the DNA Zoo consortium (dnazoo.org), which acknowledges support from Illumina, IBM, and Pawsey Supercomputing Center.

## 9. Contributions

SRF, DLE, AMV & GAD conceptualized and designed the study; SRF, RAD, EDD, AMV & OD collected tissues and data; SRF, RAD, RK & DW undertook analyses on collected data; SRF, DLE, AMV & GAD interpreted results; DLE, AMV & GAD provided resources necessary to execute the study; DLE, AMV, GAD & ELA acquired funding for the study; DLE & GAD supervised student researchers and oversaw administration for the work, SRF & DLE wrote the paper; All authors reviewed, edited and approved the paper.

## Notes

### Competing Interest Statement

The authors have declared no competing interest.

### Summary of Updates

We updated the text following some suggestions by a journal, added language surrounding methods and results of an additional analysis, and added a supplementary figure to display those results.

## References

Alonzo S.H., Sinervo B. 2001. Mate choice games, context-dependent good genes, and genetic cycles in the side-blotched lizard, Uta stansburiana. Behavioral Ecology and Sociobiology. 49:176–186.

Araya-Donoso R. 2024. Genomic and Ecological Factors Affecting Speciation Rate in Pleurodont Iguanian Lizards.

Araya-Donoso R., Biddy A., Munguía-Vega A., Lira-Noriega A., Dolby G.A. 2024. Habitat quality or quantity? Niche marginality across 21 plants and animals suggests differential responses between highland and lowland species to past climatic changes. Ecography. 2024:e07391.

Araya-Donoso R., Davalos-Dehullu E., Lukasik-Drescher Z. W., Moore D. G., Wilder, B. T., Lira-Noriega A., Munguía-Vega A., Kusumi K., Dolby G. A. 2025. Behavioral and phenotypic constraint belie deep genomic divergence and seasonal adaptation in a widespread desert lizard. bioRxiv.

Arnqvist G., Rowe L. 2023. Ecology, the pace-of-life, epistatic selection and the maintenance of genetic variation in life-history genes. Molecular Ecology. 32:4713–4724.

Baird S.F., Girard C. 1852. Characteristics of some new reptiles in the Museum of the Smithsonian Institution. Proc. Acad. Nat. Sci. Philadelphia. 6:68–70.

Bolstad G.H., Hansen T.F., Pélabon C., Falahati-Anbaran M., Pérez-Barrales R., Armbruster W.S. 2014. Genetic constraints predict evolutionary divergence in *Dalechampia* blossoms. Phil. Trans. R. Soc. B. 369:20130255.

Butlin R.K., Smadja C.M. 2018. Coupling, Reinforcement, and Speciation. The American Naturalist. 191:155–172.

Cabezas-Cartes F., Mariela Boretto J., Ruth Ibargüengoytía N. 2018. Effects of Climate and Latitude on Age at Maturity and Longevity of Lizards Studied by Skeletochronology. Integrative and Comparative Biology.

Campbell M.S., Holt C., Moore B., Yandell M. 2014. Genome Annotation and Curation Using MAKER and MAKER-P. CP in Bioinformatics. 48.

Chelini M.C., Brock K., Yeager J., Edwards D.L. 2021. Environmental drivers of sexual dimorphism in a lizard with alternative mating strategies. J of Evolutionary Biology. 34:1241–1255.

Cheng H., Concepcion G.T., Feng X., Zhang H., Li H. 2021. Haplotype-resolved de novo assembly using phased assembly graphs with hifiasm. Nat Methods. 18:170–175.

Clark G.W., Bradford J. 1969. Blood Parasites of Some Reptiles of the Pacific Northwest. The Journal of Protozoology. 16:578–581.

Corl A., Davis A.R., Kuchta S.R., Sinervo B. 2010. Selective loss of polymorphic mating types is associated with rapid phenotypic evolution during morphic speciation. Proceedings of the National Academy of Sciences. 107:4254–4259.

Corl A., Lancaster L.T., Sinervo B. 2012. Rapid Formation of Reproductive Isolation between Two Populations of Side-Blotched Lizards, Uta stansburiana. Copeia. 2012:593–602.

Da Veiga Leprevost F., Grüning B.A., Alves Aflitos S., Röst H.L., Uszkoreit J., Barsnes H., Vaudel M., Moreno P., Gatto L., Weber J., Bai M., Jimenez R.C., Sachsenberg T., Pfeuffer J., Vera Alvarez R., Griss J., Nesvizhskii A.I., Perez-Riverol Y. 2017. BioContainers: an open-source and community-driven framework for software standardization. Bioinformatics. 33:2580–2582.

Dammhahn M., Dingemanse N.J., Niemelä P.T., Réale D. 2018. Pace-of-life syndromes: a framework for the adaptive integration of behaviour, physiology and life history. Behav Ecol Sociobiol. 72:62, s00265-018-2473-y.

Davalos-Dehullu E., Baty S.M., Fisher R.N., Scott P.A., Dolby G.A., Munguia-Vega A., Cortez D. 2023. Chromosome-Level Genome Assembly of the Blacktail Brush Lizard, *Urosaurus nigricaudus*, Reveals Dosage Compensation in an Endemic Lizard. Genome Biology and Evolution. 15:evad210.

De Meester G., Van Linden L., Torfs J., Pafilis P., Šunje E., Steenssens D., Zulčić T., Sassalos A., Van Damme R. 2022. Learning with lacertids: Studying the link between ecology and cognition within a comparative framework. Evolution. 76:2531–2552.

Dobin A., Davis C.A., Schlesinger F., Drenkow J., Zaleski C., Jha S., Batut P., Chaisson M., Gingeras T.R. 2013. STAR: ultrafast universal RNA-seq aligner. Bioinformatics. 29:15–21.

Dolby G.A., Bennett S.E.K., Dorsey R.J., Stokes M.F., Riddle B.R., Lira-Noriega A., Munguia-Vega A., Wilder B.T. 2022. Integrating Earth–life systems: a geogenomic approach. Trends in Ecology & Evolution. 37:371–384.

Dolby G.A., Bennett S.E.K., Lira-Noriega A., Wilder B.T., Munguía-Vega A. 2015. Assessing the Geological and Climatic Forcing of Biodiversity and Evolution Surrounding the Gulf of California. jsw. 57:391–455.

Dudchenko O., Batra S.S., Omer A.D., Nyquist S.K., Hoeger M., Durand N.C., Shamim M.S., Machol I., Lander E.S., Aiden A.P., Aiden E.L. 2017. De novo assembly of the *Aedes aegypti* genome using Hi-C yields chromosome-length scaffolds. Science. 356:92–95.

Dudchenko O., Pham M., Lui C., Batra S.S., Hoeger M., Nyquist S.K., Durand N.C., Shamim M.S., Machol I., Erskine W., Aiden E.L., Kaur P. 2018. Hi-C yields chromosome-length scaffolds for a legume genome, Trifolium subterraneum.

Durand N.C., Shamim M.S., Machol I., Rao S.S.P., Huntley M.H., Lander E.S., Aiden E.L. 2016. Juicer Provides a One-Click System for Analyzing Loop-Resolution Hi-C Experiments. Cell Systems. 3:95–98.

Ewels P.A., Peltzer A., Fillinger S., Patel H., Alneberg J., Wilm A., Garcia M.U., Di Tommaso P., Nahnsen S. 2020. The nf-core framework for community-curated bioinformatics pipelines. Nat Biotechnol. 38:271–278.

Fick S.E., Hijmans R.J. 2017. WorldClim 2: new 1-km spatial resolution climate surfaces for global land areas. Int. J. Climatol. 37:4302–4315.

Friesen C.R., Johansson R., Olsson M. 2017. Morph-specific metabolic rate and the timing of reproductive senescence in a color polymorphic dragon. J Exp Zool Pt A. 327:433–443.

Gambón-Deza F. 2023. A new T Cell Receptor in Squamata Reptiles. bioRxiv.

Giraudeau M., Angelier F., Sepp T. 2019. Do Telomeres Influence Pace-of-Life-Strategies in Response to Environmental Conditions Over a Lifetime and Between Generations? BioEssays. 41:1800162.

Goulet C.T., Michelangeli M., Chung M., Riley J.L., Wong B.B.M., Thompson M.B., Chapple D.G. 2018. Evaluating cognition and thermal physiology as components of the pace-of-life syndrome. Evol Ecol. 32:469–488.

Grüning B., Dale R., Sjödin A., Chapman B.A., Rowe J., Tomkins-Tinch C.H., Valieris R., Köster J. 2018. Bioconda: sustainable and comprehensive software distribution for the life sciences. Nat Methods. 15:475–476.

Hämäläinen A.M., Guenther A., Patrick S.C., Schuett W. 2021. Environmental effects on the covariation among pace-of-life traits. Ethology. 127:32–44.

Hazard L.C., Nagy K.A., Miles D.B., Svensson E.I., Costa D., Sinervo B. 2019. Integration of Genotype, Physiological Performance, and Survival in a Lizard (*Uta stansburiana*) with Alternative Mating Strategies. Physiological and Biochemical Zoology. 92:303–315.

Hollingsworth B.D. 1999.The Molecular Systematics of the Side-blotched Lizards (IGUANIA: PHRYNOSOMATIDAE: UTA).

Hu C.K., York R.A., Metz H.C., Bedford N.L., Fraser H.B., Hoekstra H.E. 2022. cis-Regulatory changes in locomotor genes are associated with the evolution of burrowing behavior. Cell Reports. 38:110360.

Hudson S.B., Lidgard A.D., French S.S. 2020. Glucocorticoids, energy metabolites, and immunity vary across allostatic states for plateau side-blotched lizards (*Uta stansburiana uniformis*) residing in a heterogeneous thermal environment. J Exp Zool Pt A. 333:732–743.

Immonen E., Hämäläinen A., Schuett W., Tarka M. 2018. Evolution of sex-specific pace-of-life syndromes: genetic architecture and physiological mechanisms. Behav Ecol Sociobiol. 72:60.

Koochekian N., Ascanio A., Farleigh K., Card D.C., Schield D.R., Castoe T.A., Jezkova T. 2022. A chromosome-level genome assembly and annotation of the desert horned lizard, *Phrynosoma platyrhinos*, provides insight into chromosomal rearrangements among reptiles. GigaScience. 11:giab098.

Korf I. 2004. Gene finding in novel genomes. BMC Bioinformatics. 5:59.

Krueger F., James F., Ewels P., Afyounian E., Schuster-Boeckler B. 2021. Trim Galore.

LaDage L.D., Maged R.M., Forney M.V., Roth T.C., Sinervo B., Pravosudov V.V. 2013. Interaction between territoriality, spatial environment, and hippocampal neurogenesis in male side-blotched lizards. Behavioral Neuroscience. 127:555–565.

LaDage L.D., Roth T.C., Sinervo B., Pravosudov V.V. 2016. Environmental experiences influence cortical volume in territorial and nonterritorial side-blotched lizards, Uta stansburiana. Animal Behaviour. 115:11–18.

LaDage L.D., Yu T., Zani P.A. 2022. Higher Rate of Male Sexual Displays Correlates with Larger Ventral Posterior Amygdala Volume and Neuron Soma Volume in Wild-Caught Common Side-Blotched Lizards, *Uta stansburiana*. Brain Behav Evol. 97:298–308.

Leaché A.D., Sites, Jr. J.W. 2009. Chromosome Evolution and Diversification in North American Spiny Lizards (Genus *Sceloporus*). Cytogenet Genome Res. 127:166–181.

Li H. 2013. Aligning sequence reads, clone sequences and assembly contigs with BWA-MEM.

Li H. 2018. Minimap2: pairwise alignment for nucleotide sequences. Bioinformatics. 34:3094–3100.

Lim S.L., Tsend-Ayush E., Kortschak R.D., Jacob R., Ricciardelli C., Oehler M.K., Grützner F. 2013. Conservation and Expression of PIWI-Interacting RNA Pathway Genes in Male and Female Adult Gonad of Amniotes1. Biology of Reproduction. 89.

Lipshutz S.E., Hibbins M.S., Bentz A.B., Buechlein A.M., Empson T.A., George E.M., Hauber M.E., Rusch D.B., Schelsky W.M., Thomas Q.K., Torneo S.J., Turner A.M., Wolf S.E., Woodruff M.J., Hahn M.W., Rosvall K.A. 2025. Repeated behavioural evolution is associated with convergence of gene expression in cavity-nesting songbirds. Nat Ecol Evol. 9:845–856.

Lisachov A.P., Tishakova K.V., Romanenko S.A., Molodtseva A.S., Prokopov D.Yu., Pereira J.C., Ferguson-Smith M.A., Borodin P.M., Trifonov V.A. 2021. Whole-chromosome fusions in the karyotype evolution of *Sceloporus* (Iguania, Reptilia) are more frequent in sex chromosomes than autosomes. Phil. Trans. R. Soc. B. 376:20200099.

Love M.I., Huber W., Anders S. 2014. Moderated estimation of fold change and dispersion for RNA-seq data with DESeq2. Genome Biol. 15:550.

Lovell J.T., Sreedasyam A., Schranz M.E., Wilson M., Carlson J.W., Harkess A., Emms D., Goodstein D.M., Schmutz J. 2022. GENESPACE tracks regions of interest and gene copy number variation across multiple genomes. eLife. 11:e78526.

MacDonald A.J., Sarre S.D., FitzSimmons N.N., Aitken N. 2011. Determining microsatellite genotyping reliability and mutation detection ability: an approach using small-pool PCR from sperm DNA. Mol Genet Genomics. 285:1–18.

Maher P. 2005. The effects of stress and aging on glutathione metabolism. Ageing Research Reviews. 4:288–314.

Malisch J.L., Garland T., Claggett L., Stevenson L., Kohl E.A., John-Alder H.B. 2020. Living on the edge: Glucocorticoid physiology in desert iguanas (Dipsosaurus dorsalis) is predicted by distance from an anthropogenic disturbance, body condition, and population density. General and Comparative Endocrinology. 294:113468.

Marin R., Cortez D., Lamanna F., Pradeepa M.M., Leushkin E., Julien P., Liechti A., Halbert J., Brüning T., Mössinger K., Trefzer T., Conrad C., Kerver H.N., Wade J., Tschopp P., Kaessmann H. 2017. Convergent origination of a *Drosophila*-like dosage compensation mechanism in a reptile lineage. Genome Res. 27:1974–1987.

McKinney C.O. 1971. An Analysis of Zones of Intergradation in the Side-Blotched Lizard, Uta stansburiana (Sauria: Iguanidae). Copeia. 1971:596.

McWhinnie D.J., Cortelyou J.R. 1968. Influence of parathyroid extract on blood and urine mineral levels in iguanid lizards. General and Comparative Endocrinology. 11:78–87.

Mellor N.J., Webster T.H., Byrne H., Williams A.S., Edwards T., DeNardo D.F., Wilson M.A., Kusumi K., Dolby G.A. 2025. Divergence in Regulatory Regions and Gene Duplications May Underlie Chronobiological Adaptation in Desert Tortoises. Molecular Ecology. 34:e17600.

Mettke-Hofmann C. 2014. Cognitive ecology: ecological factors, life-styles, and cognition. WIRES Cognitive Science. 5:345–360.

Montiglio P.-O., Dammhahn M., Dubuc Messier G., Réale D. 2018. The pace-of-life syndrome revisited: the role of ecological conditions and natural history on the slow-fast continuum. Behav Ecol Sociobiol. 72:116.

Moore E., Arnscheidt A., Ger K. 2004. Simplified protocols for the preparation of genomic DNA from bacterial cultures. Molecular Microbial Ecology Manual. Kluwer Academic Publishers. p. 3–18.

Narum S.R., Di Genova A., Micheletti S.J., Maass A. 2018. Genomic variation underlying complex life-history traits revealed by genome sequencing in Chinook salmon. Proc. R. Soc. B. 285:20180935.

Oliva M., Muñoz-Aguirre M., Kim-Hellmuth S., Wucher V., Gewirtz A.D.H., Cotter D.J., Parsana P., Kasela S., Balliu B., Viñuela A., Castel S.E., Mohammadi P., Aguet F., Zou Y., Khramtsova E.A., Skol A.D., Garrido-Martín D., Reverter F., Brown A., Evans P., Gamazon E.R., Payne A., Bonazzola R., Barbeira A.N., Hamel A.R., Martinez-Perez A., Soria J.M., GTEx Consortium, Pierce B.L., Stephens M., Eskin E., Dermitzakis E.T., Segrè A.V., Im H.K., Engelhardt B.E., Ardlie K.G., Montgomery S.B., Battle A.J., Lappalainen T., Guigó R., Stranger B.E., Aguet F., Anand S., Ardlie K.G., Gabriel S., Getz G.A., Graubert A., Hadley K., Handsaker R.E., Huang K.H., Kashin S., Li X., MacArthur D.G., Meier S.R., Nedzel J.L., Nguyen D.T., Segrè A.V., Todres E., Balliu B., Barbeira A.N., Battle A., Bonazzola R., Brown A., Brown C.D., Castel S.E., Conrad D.F., Cotter D.J., Cox N., Das S., De Goede O.M., Dermitzakis E.T., Einson J., Engelhardt B.E., Eskin E., Eulalio T.Y., Ferraro N.M., Flynn E.D., Fresard L., Gamazon E.R., Garrido-Martín D., Gay N.R., Gloudemans M.J., Guigó R., Hame A.R., He Y., Hoffman P.J., Hormozdiari F., Hou L., Im H.K., Jo B., Kasela S., Kellis M., Kim-Hellmuth S., Kwong A., Lappalainen T., Li X., Liang Y., Mangul S., Mohammadi P., Montgomery S.B., Muñoz-Aguirre M., Nachun D.C., Nobel A.B., Oliva M., Park Y., Park Y., Parsana P., Rao A.S., Reverter F., Rouhana J.M., Sabatti C., Saha A., Stephens M., Stranger B.E., Strober B.J., Teran N.A., Viñuela A., Wang G., Wen X., Wright F., Wucher V., Zou Y., Ferreira P.G., Li G., Melé M., Yeger-Lotem E., Barcus M.E., Bradbury D., Krubit T., McLean J.A., Qi L., Robinson K., Roche N.V., Smith A.M., Sobin L., Tabor D.E., Undale A., Bridge J., Brigham L.E., Foster B.A., Gillard B.M., Hasz R., Hunter M., Johns C., Johnson M., Karasik E., Kopen G., Leinweber W.F., McDonald A., Moser M.T., Myer K., Ramsey K.D., Roe B., Shad S., Thomas J.A., Walters G., Washington M., Wheeler J., Jewell S.D., Rohrer D.C., Valley D.R., Davis D.A., Mash D.C., Branton P.A., Barker L.K., Gardiner H.M., Mosavel M., Siminoff L.A., Flicek P., Haeussler M., Juettemann T., Kent W.J., Lee C.M., Powell C.C., Rosenbloom K.R., Ruffier M., Sheppard D., Taylor K., Trevanion S.J., Zerbino D.R., Abell N.S., Akey J., Chen L., Demanelis K., Doherty J.A., Feinberg A.P., Hansen K.D., Hickey P.F., Jasmine F., Jiang L., Kaul R., Kibriya M.G., Li J.B., Li Q., Lin S., Linder S.E., Pierce B.L., Rizzardi L.F., Skol A.D., Smith K.S., Snyder M., Stamatoyannopoulos J., Tang H., Wang M., Carithers L.J., Guan P., Koester S.E., Little A.R., Moore H.M., Nierras C.R., Rao A.K., Vaught J.B., Volpi S. 2020. The impact of sex on gene expression across human tissues. Science. 369:eaba3066.

Park W., Rengaraj D., Kil D.-Y., Kim H., Lee H.-K., Song K.-D. 2017. RNA-seq analysis of the kidneys of broiler chickens fed diets containing different concentrations of calcium. Sci Rep. 7:11740.

Parker W.S. 1974. Home Range, Growth, and Population Density of Uta stansburiana in Arizona. Journal of Herpetology. 8:135.

Parker W.S., Pianka E.R. 1975. Comparative Ecology of Populations of the Lizard Uta stansburiana. Copeia. 1975:615.

Patel H., Ewels P., Peltzer A., Botvinnik O., Sturm G., Moreno D., Vemuri P., Garcia M.U., silviamorins, Pantano L., Binzer-Panchal M., bot nf-core, Syme R., Zepper M., Kelly G., Hanssen F., Yates J.A.F., Cheshire C., rfenouil, Espinosa-Carrasco J., marchoeppner, Miller E., Talbot A., Zhou P., Guinchard S., Hörtenhuber M., Gabernet G., Mertes C., Straub D., Tommaso P.D. 2023. nf-core/rnaseq: nf-core/rnaseq v3.12.0 - Osmium Octopus.

Patro R., Duggal G., Love M.I., Irizarry R.A., Kingsford C. 2017. Salmon provides fast and bias-aware quantification of transcript expression. Nat Methods. 14:417–419.

Pennock L.A., Tinkle D W, Shaw M W. 1969. Minute Y chromosome in the lizard genus Uta (family Iguanidae). Cytogenetics. 8:9–19.

Pennock LewisA., Tinkle DonaldW., Shaw MargeryW. 1968. Chromosome number in the lizard genus Uta (family Iguanidae). Chromosoma. 24.

Perry B.W., Schield D.R., Adams R.H., Castoe T.A. 2021. Microchromosomes Exhibit Distinct Features of Vertebrate Chromosome Structure and Function with Underappreciated Ramifications for Genome Evolution. Molecular Biology and Evolution. 38:904–910.

Pinto B.J., Gamble T., Smith C.H., Wilson M.A. 2023. A lizard is never late: Squamate genomics as a recent catalyst for understanding sex chromosome and microchromosome evolution. Journal of Heredity. 114:445–458.

Prabh N., Linnenbrink M., Jovicic M., Guenther A. 2023. Fast adjustment of pace-of-life and risk-taking to changes in food quality by altered gene expression in house mice. Ecology Letters. 26:99–110.

Quillfeldt P., Romeike T., Masello J.F., Reiner G., Willems H., Bedolla-Guzmán Y. 2018. Molecular survey of coccidian infections of the side-blotched lizard *Uta stansburiana* on San Benito Oeste Island, Mexico. Parasite. 25:43.

Quinlan A.R., Hall I.M. 2010. BEDTools: a flexible suite of utilities for comparing genomic features. Bioinformatics. 26:841–842.

Rao S.S.P., Huntley M.H., Durand N.C., Stamenova E.K., Bochkov I.D., Robinson J.T., Sanborn A.L., Machol I., Omer A.D., Lander E.S., Aiden E.L. 2014. A 3D Map of the Human Genome at Kilobase Resolution Reveals Principles of Chromatin Looping. Cell. 159:1665–1680.

Raudvere U., Kolberg L., Kuzmin I., Arak T., Adler P., Peterson H., Vilo J. 2019. g:Profiler: a web server for functional enrichment analysis and conversions of gene lists (2019 update). Nucleic Acids Research. 47:W191–W198.

Réale D., Garant D., Humphries M.M., Bergeron P., Careau V., Montiglio P.-O. 2010. Personality and the emergence of the pace-of-life syndrome concept at the population level. Phil. Trans. R. Soc. B. 365:4051–4063.

Réale D., Reader S.M., Sol D., McDougall P.T., Dingemanse N.J. 2007. Integrating animal temperament within ecology and evolution. Biological Reviews. 82:291–318.

Ribeiro J.R., Freiman R.N. 2014. Estrogen signaling crosstalk: Implications for endocrine resistance in ovarian cancer. The Journal of Steroid Biochemistry and Molecular Biology. 143:160–173.

Ricklefs R.E., Wikelski M. 2002. The physiology/life-history nexus. Trends in Ecology & Evolution. 17:462–468.

Royauté R., Berdal M.A., Garrison C.R., Dochtermann N.A. 2018. Paceless life? A meta-analysis of the pace-of-life syndrome hypothesis. Behav Ecol Sociobiol. 72:64.

Sampson J.M., Morrissey K.A., Mikolajova K.J., Zimmerly K.M., Gemmell N.J., Gardner M.G., Bertozzi T., Miller R.D. 2025. Squamate reptiles may have compensated for the lack of γδTCR with a duplication of the TRB locus. Front. Immunol. 15:1524471.

Schield D.R., Card D.C., Hales N.R., Perry B.W., Pasquesi G.M., Blackmon H., Adams R.H., Corbin A.B., Smith C.F., Ramesh B., Demuth J.P., Betrán E., Tollis M., Meik J.M., Mackessy S.P., Castoe T.A. 2019. The origins and evolution of chromosomes, dosage compensation, and mechanisms underlying venom regulation in snakes. Genome Res. 29:590–601.

Schlippe Justicia L., Mayer M., Lorioux-Chevalier U., Dittrich C., Rojas B., Chouteau M. 2024. Intraspecific divergence of sexual size dimorphism and reproductive strategies in a polytypic poison frog. Evol Ecol. 38:121–139.

Shannon P., Markiel A., Ozier O., Baliga N.S., Wang J.T., Ramage D., Amin N., Schwikowski B., Ideker T. 2003. Cytoscape: A Software Environment for Integrated Models of Biomolecular Interaction Networks. Genome Res. 13:2498–2504.

Simão F.A., Waterhouse R.M., Ioannidis P., Kriventseva E.V., Zdobnov E.M. 2015. BUSCO: assessing genome assembly and annotation completeness with single-copy orthologs. Bioinformatics. 31:3210–3212.

Sinervo B., Bleay C., Adamopoulou C. 2001. SOCIAL CAUSES OF CORRELATIONAL SELECTION AND THE RESOLUTION OF A HERITABLE THROAT COLOR POLYMORPHISM IN A LIZARD. Evolution. 55:2040–2052.

Sinervo B., Lively C.M. 1996. The rock–paper–scissors game and the evolution of alternative male strategies. Nature. 380:240–243.

Sinervo B., Svensson E., Comendant T. 2000. Density cycles and an offspring quantity and quality game driven by natural selection. Nature. 406:985–988.

Smit A.F.A., Hubley R., Green P. 2015a. RepeatModeler.

Smit A.F.A., Hubley R., Green P. 2015b. RepeatMasker.

Smith G.D., Zani P.A., French S.S. 2019. Life-history differences across latitude in common side-blotched lizards (*Uta stansburiana*). Ecology and Evolution. 9:5743–5751.

Soneson C., Love M.I., Robinson M.D. 2015. Differential analyses for RNA-seq: transcript-level estimates improve gene-level inferences. F1000Res. 4:1521.

Spence A.R., Durso A.M., Smith G.D., Skinner H.M., French S.S. 2017. Physiological Correlates of Multiple Parasitic Infections in Side-Blotched Lizards. Physiological and Biochemical Zoology. 90:321–327.

Srikulnath K., Ahmad S.F., Singchat W., Panthum T. 2021. Why Do Some Vertebrates Have Microchromosomes? Cells. 10:2182.

Stalker L., Russell S.J., Co C., Foster R.A., LaMarre J. 2016. PIWIL1 Is Expressed in the Canine Testis, Increases with Sexual Maturity, and Binds Small RNAs1. Biology of Reproduction. 94.

Stanke M., Keller O., Gunduz I., Hayes A., Waack S., Morgenstern B. 2006. AUGUSTUS: ab initio prediction of alternative transcripts. Nucleic Acids Research. 34:W435–W439.

Stebbins R.C. 2003. A Field Guide to Western Reptiles and Amphibians. Boston: Houghton Mifflin Company.

Supek F., Bošnjak M., Škunca N., Šmuc T. 2011. REVIGO Summarizes and Visualizes Long Lists of Gene Ontology Terms. PLoS ONE. 6:e21800.

Syding L.A., Kubik-Zahorodna A., Reguera D.P., Nickl P., Hruskova B., Kralikova M., Kopkanova J., Novosadova V., Kasparek P., Prochazka J., Rozman J., Turecek R., Sedlacek R. 2023. Ablation of Gabra5 Influences Corticosterone Levels and Anxiety-like Behavior in Mice. Genes. 14:285.

Szklarczyk D., Kirsch R., Koutrouli M., Nastou K., Mehryary F., Hachilif R., Gable A.L., Fang T., Doncheva N.T., Pyysalo S., Bork P., Jensen L.J., von Mering C. 2023. The STRING database in 2023: protein–protein association networks and functional enrichment analyses for any sequenced genome of interest. Nucleic Acids Research. 51:D638–D646.

Tan S., Rudd J.A., Yew D.T. 2011. Gene Expression Changes in GABAA Receptors and Cognition Following Chronic Ketamine Administration in Mice. PLoS ONE. 6:e21328.

Ujvari B., Raven N., Madsen T., Klaassen M., Dujon A.M., Schultz A.G., Nunney L., Lemaître J., Giraudeau M., Thomas F. 2022. Telomeres, the loop tying cancer to organismal life-histories. Molecular Ecology. 31:6273–6285.

Upton D.E., Murphy R.W. 1997. Phylogeny of the Side-Blotched Lizards (Phrynosomatidae:Uta) Based on mtDNA Sequences: Support for a Midpeninsular Seaway in Baja California. Molecular Phylogenetics and Evolution. 8:104–113.

Vasilieva N.A. 2022. Pace-of-Life Syndrome (POLS): Evolution of the Concept. Biol Bull Russ Acad Sci. 49:750–762.

Waits D.S., Simpson D.Y., Sparkman A.M., Bronikowski A.M., Schwartz T.S. 2020. The utility of reptile blood transcriptomes in molecular ecology. Molecular Ecology Resources. 20:308–317.

Williams J.B., Miller R.A., Harper J.M., Wiersma P. 2010. Functional Linkages for the Pace of Life, Life-history, and Environment in Birds. Integrative and Comparative Biology. 50:855–868.

Wilson B.S. 1991. Latitudinal Variation in Activity Season Mortality Rates of the Lizard Uta Stansburiana. Ecological Monographs. 61:393–414.

Wright J., Bolstad G.H., Araya-Ajoy Y.G., Dingemanse N.J. 2019. Life-history evolution under fluctuating density-dependent selection and the adaptive alignment of pace-of-life syndromes. Biological Reviews. 94:230–247.

Xu C., Dolby G.A., Drake K.K., Esque T.C., Kusumi K. 2020. Immune and sex-biased gene expression in the threatened Mojave desert tortoise, Gopherus agassizii. PLoS ONE. 15:e0238202.

Xue X., Thompson A.R., Adams B.J. 2024. An Antarctic worm and its soil ecosystem: A review of an emerging research program in ecological genomics. Applied Soil Ecology. 193:105110.

Young A.J. 2018. The role of telomeres in the mechanisms and evolution of life-history trade-offs and ageing. Phil. Trans. R. Soc. B. 373:20160452.

Zani P.A., Stein S.J. 2018. Field and laboratory responses to drought by Common Side-blotched Lizards (Uta stansburiana). Journal of Arid Environments. 154:15–23.

