## Supplementary material for "Chromosome-length genome assembly of *Uta stansburiana* and gene expression data reveal fast pace-of-life comes with environmental stability": SuppFigs.docx

**Supp. Mat. Fig 1. Sampling location with annual precipitation (mm)**

**Supp. Mat. Fig 2. Sampling location with average annual irradiation rate**

**Supp. Mat. Fig 3. Sampling locations with average annual UVB irradiation**

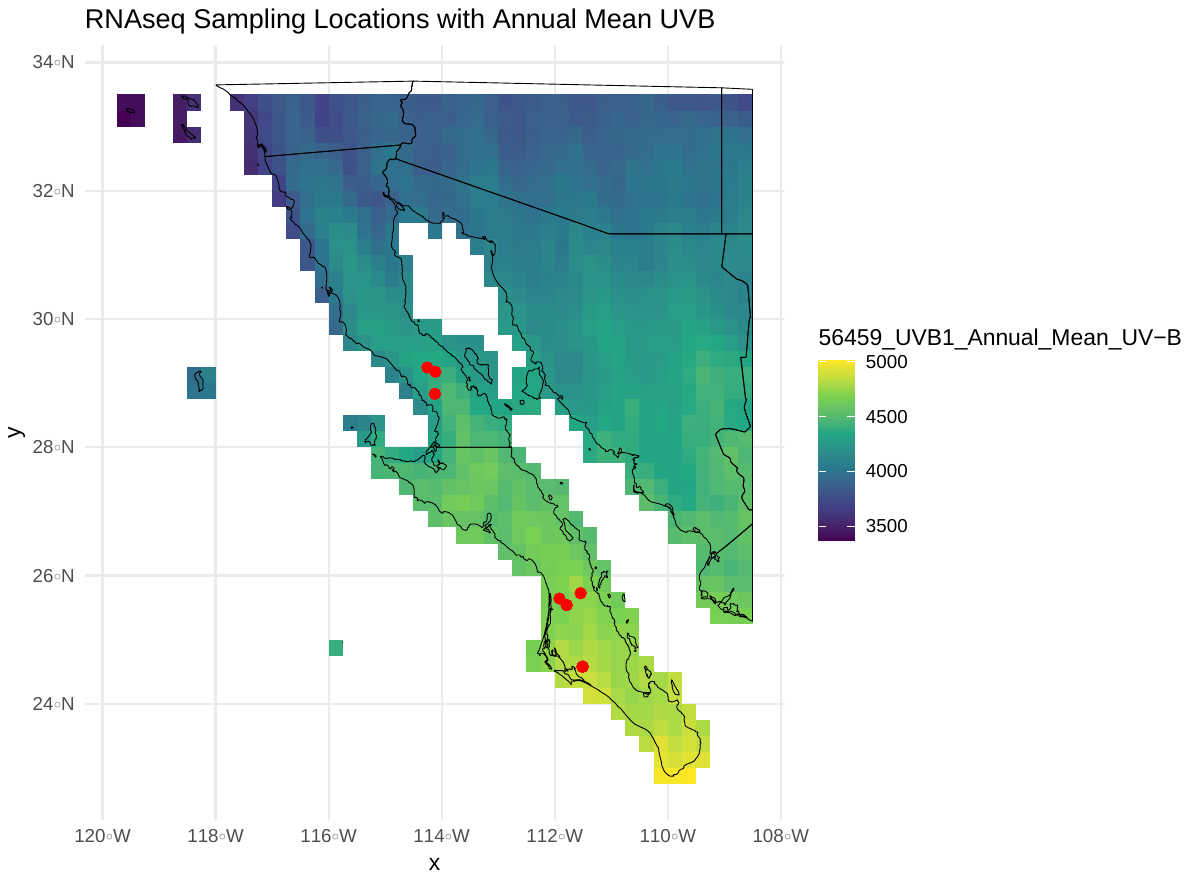

**Supp. Mat. Fig 4.** Heatmap of log2-transformed count data for genes with differential expression by region, controlling for season and sex. Samples are labeled by region, season, and sex. **B)** Principal components analysis (PC1, PC2) of regular log-transformed count data.

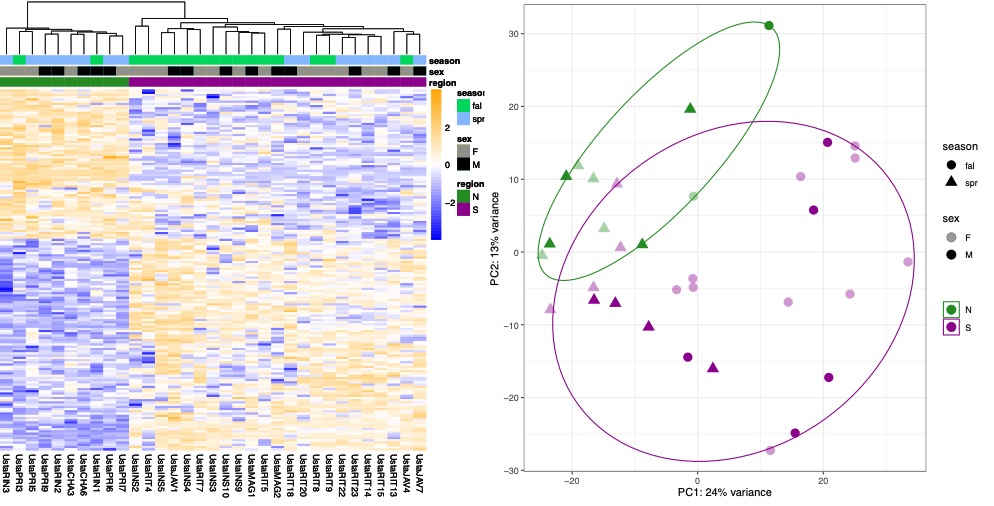

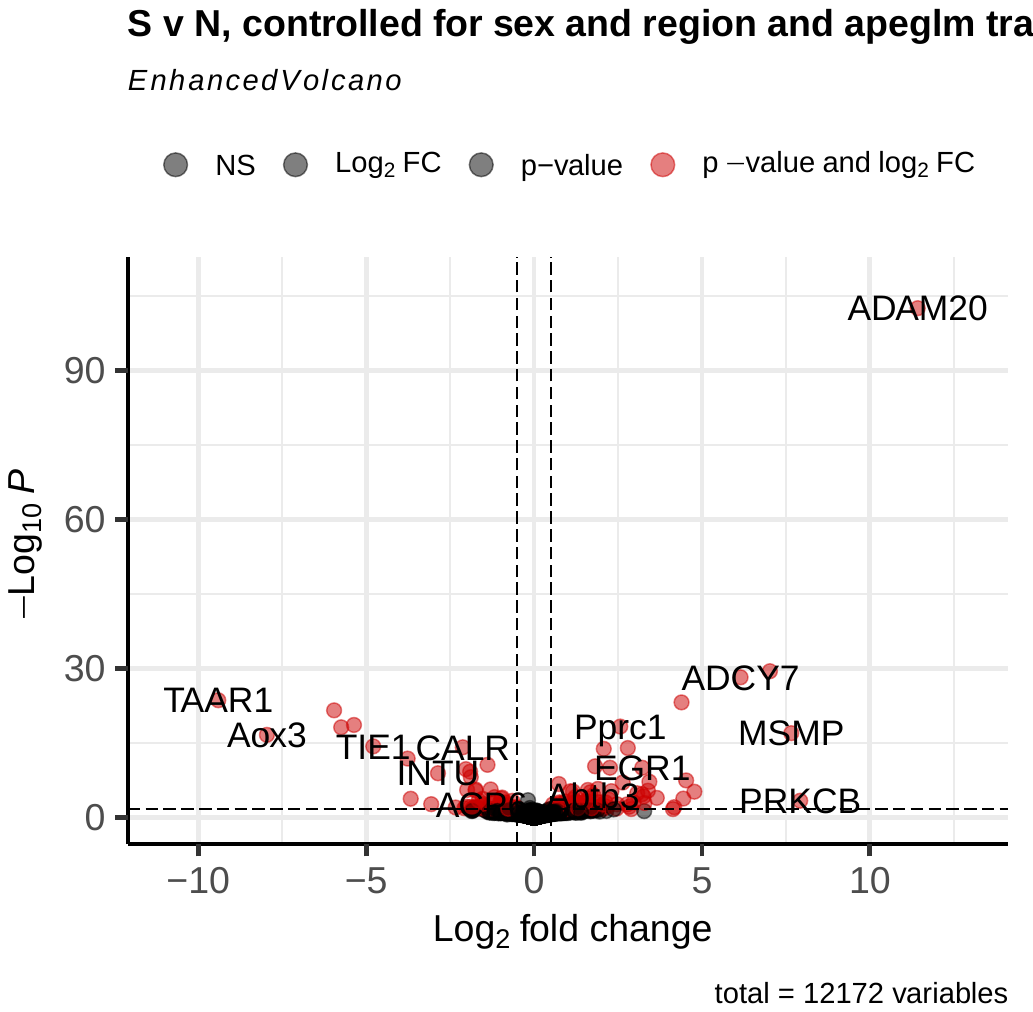

**Supp. Mat. Fig. 5. Volcano plot of genes differentially expressed between northern and southern groups, controlling for sex and seasonal differences. Positive log2 fold-change values correspond to genes differentially over-expressed in the southern group, while negative log2 fold-change values correspond to genes differentially under-expressed in the southern group.**

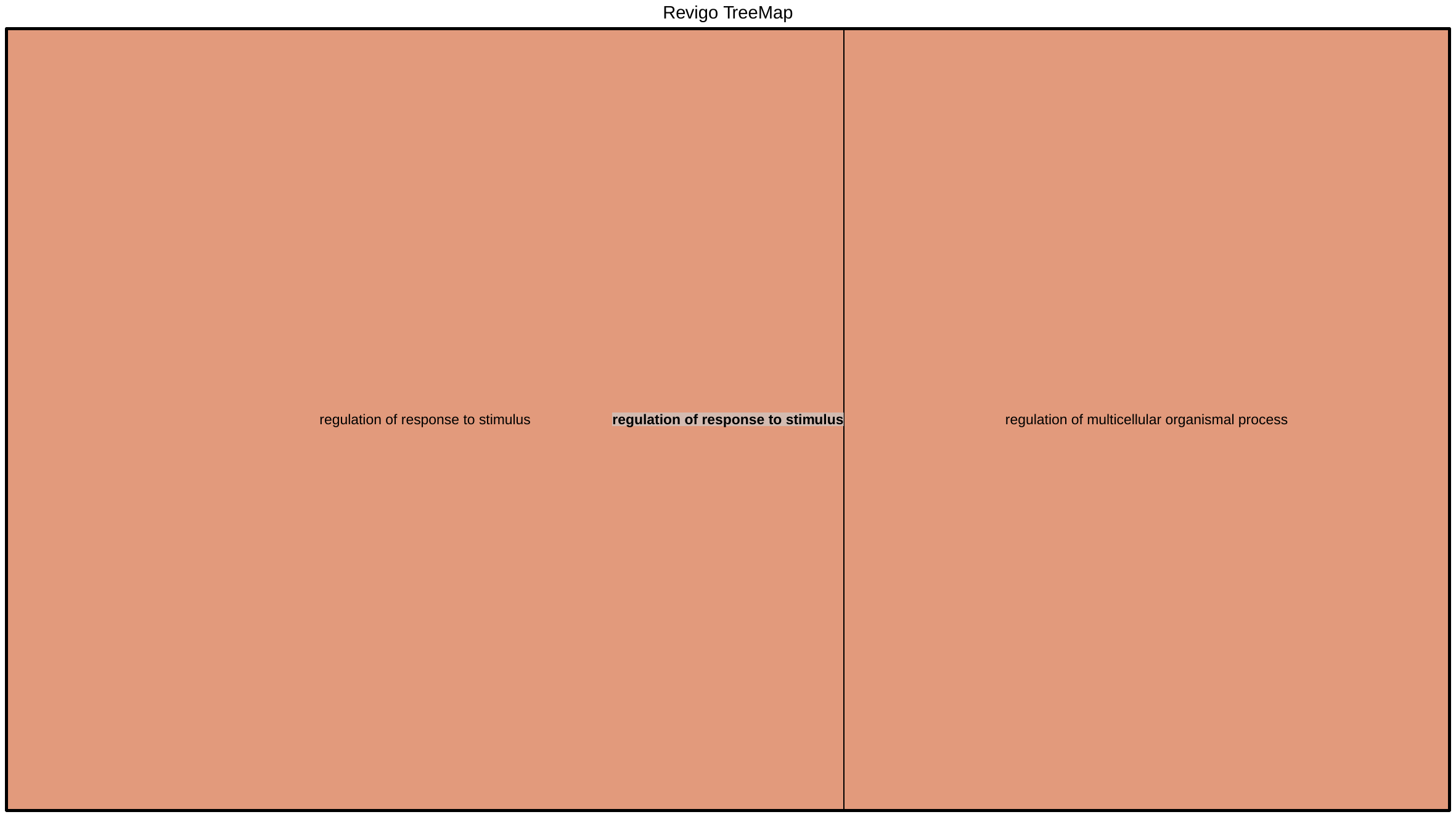

**Supp. Mat. Fig 6. Revigo treemap of enriched biological processes for the region controlling for season and sex contrast**

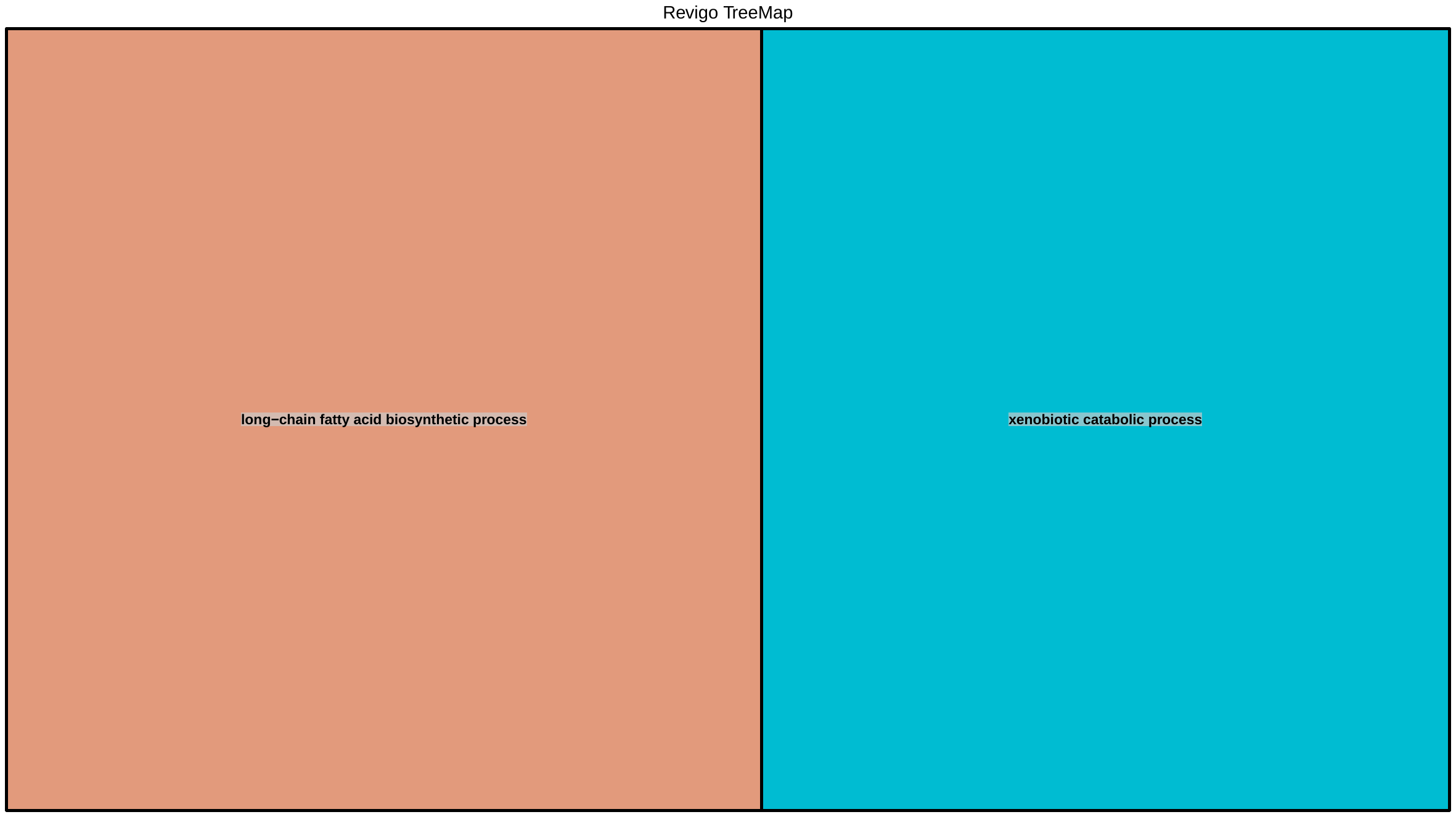

**Supp. Mat. 7. Revigo treemap of enriched biological processes for contrast by sex**

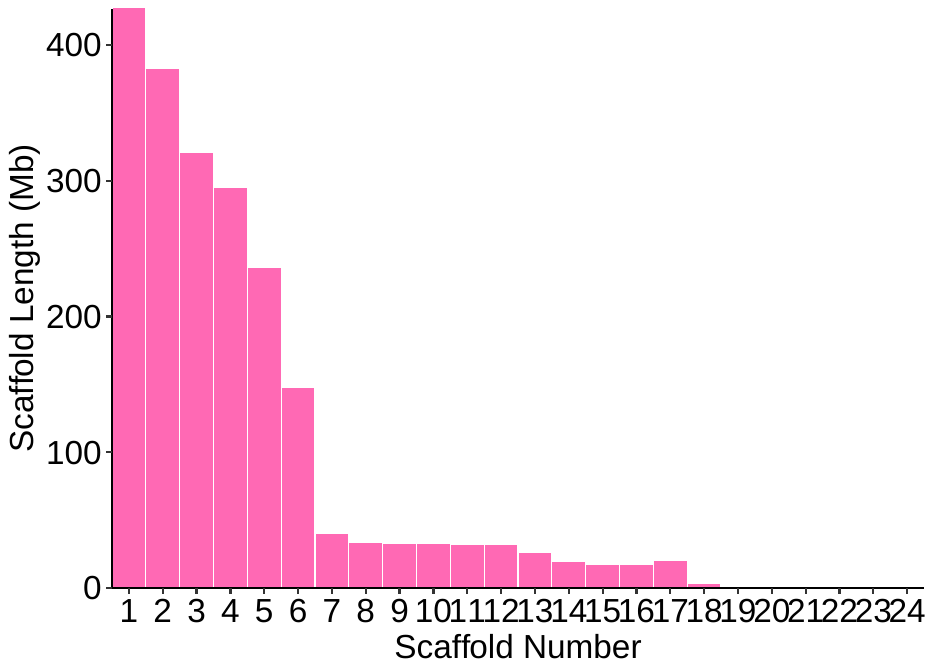

**Supp. Mat. 8. Distribution of scaffold size. Scaffold 1-17 correspond to HiC_scaffolds_1-17. HiC_scaffold_18 is likely the Y chromosome.**

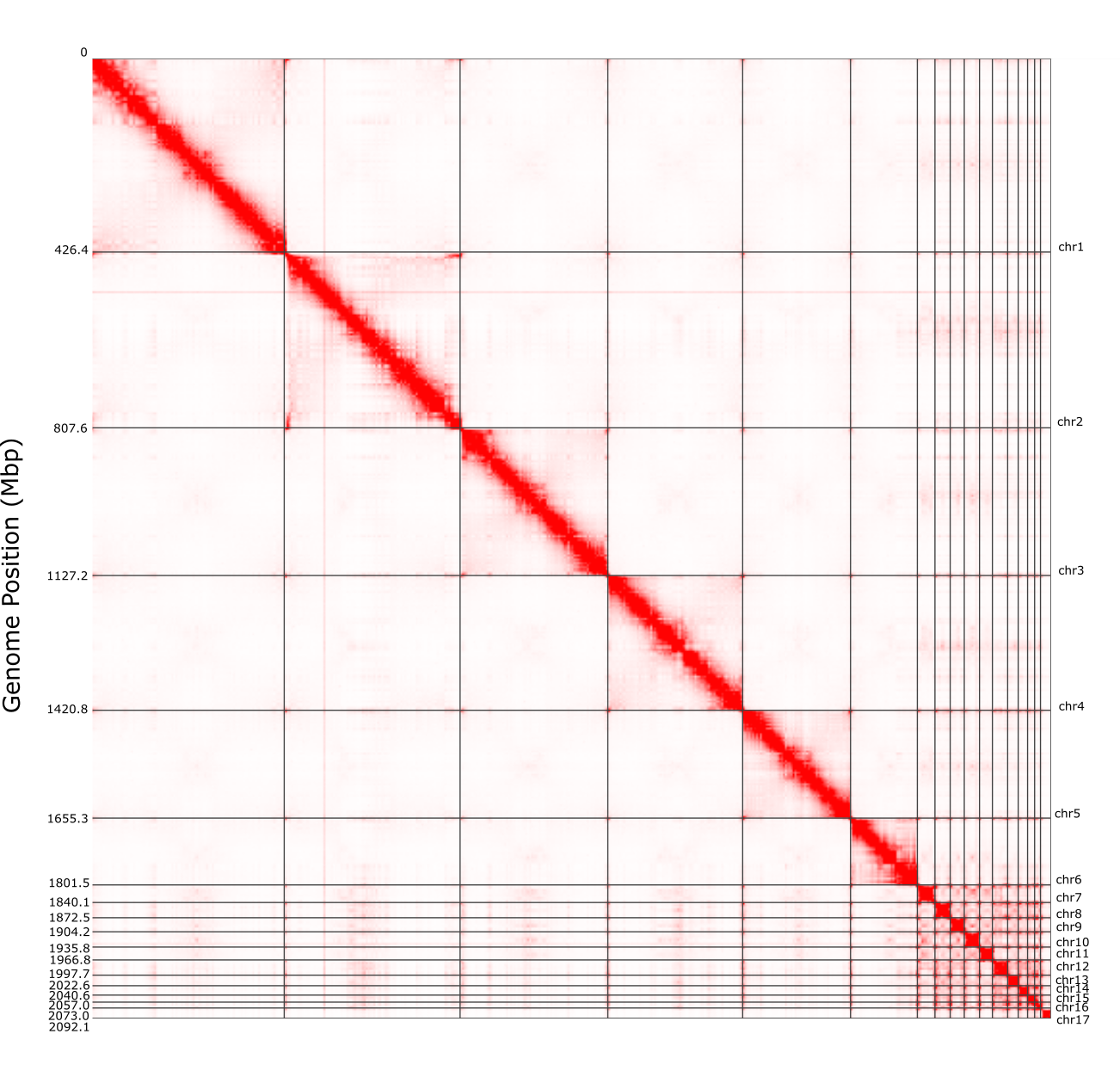

**Supp. Mat. 9. Interaction map of Hi-C chromosome-level data.**

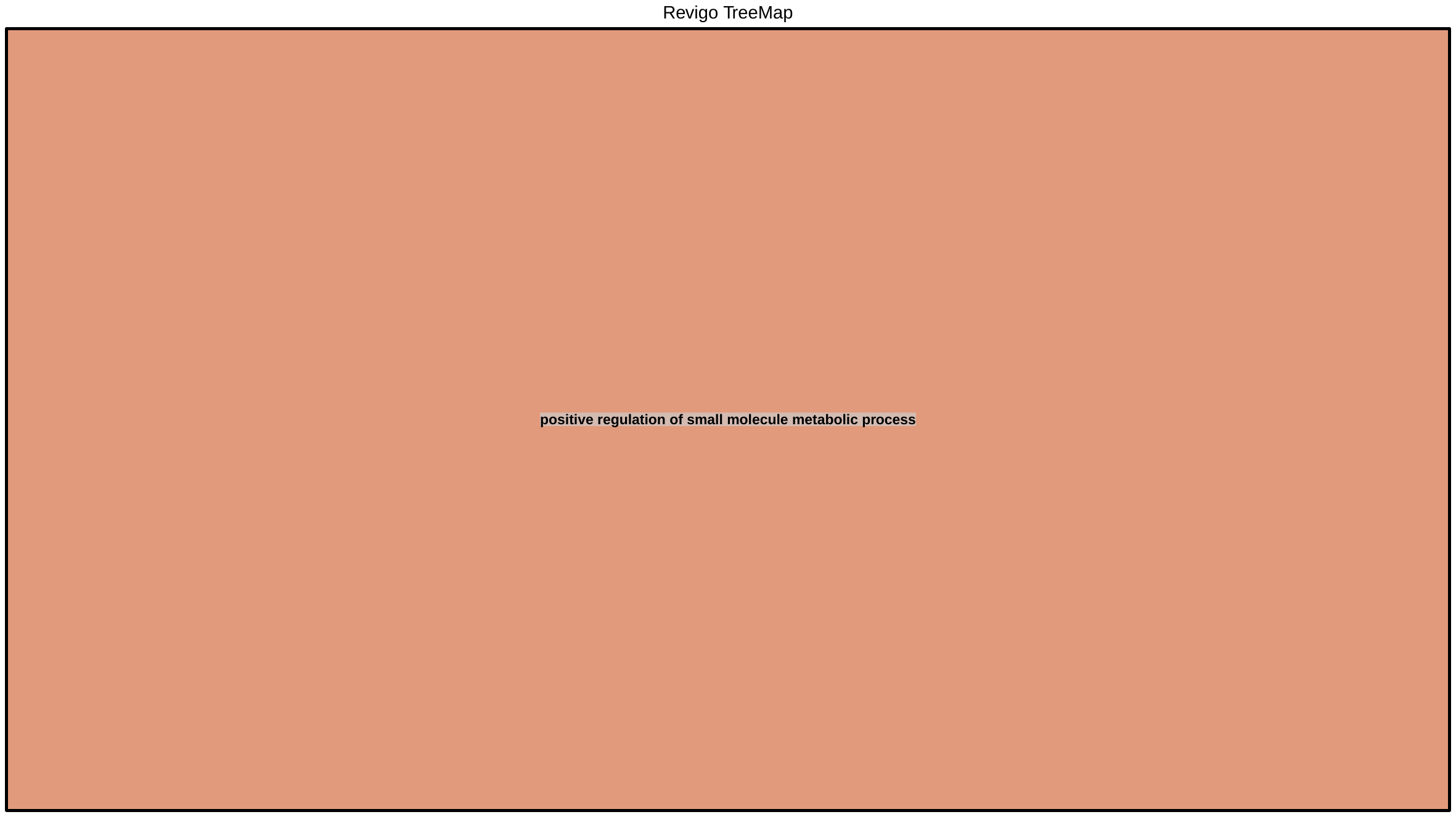

**Supp. Mat. 10. Revigo treemap of enriched biological processes for yellow vs. orange- and blue-throated lizard contrast**

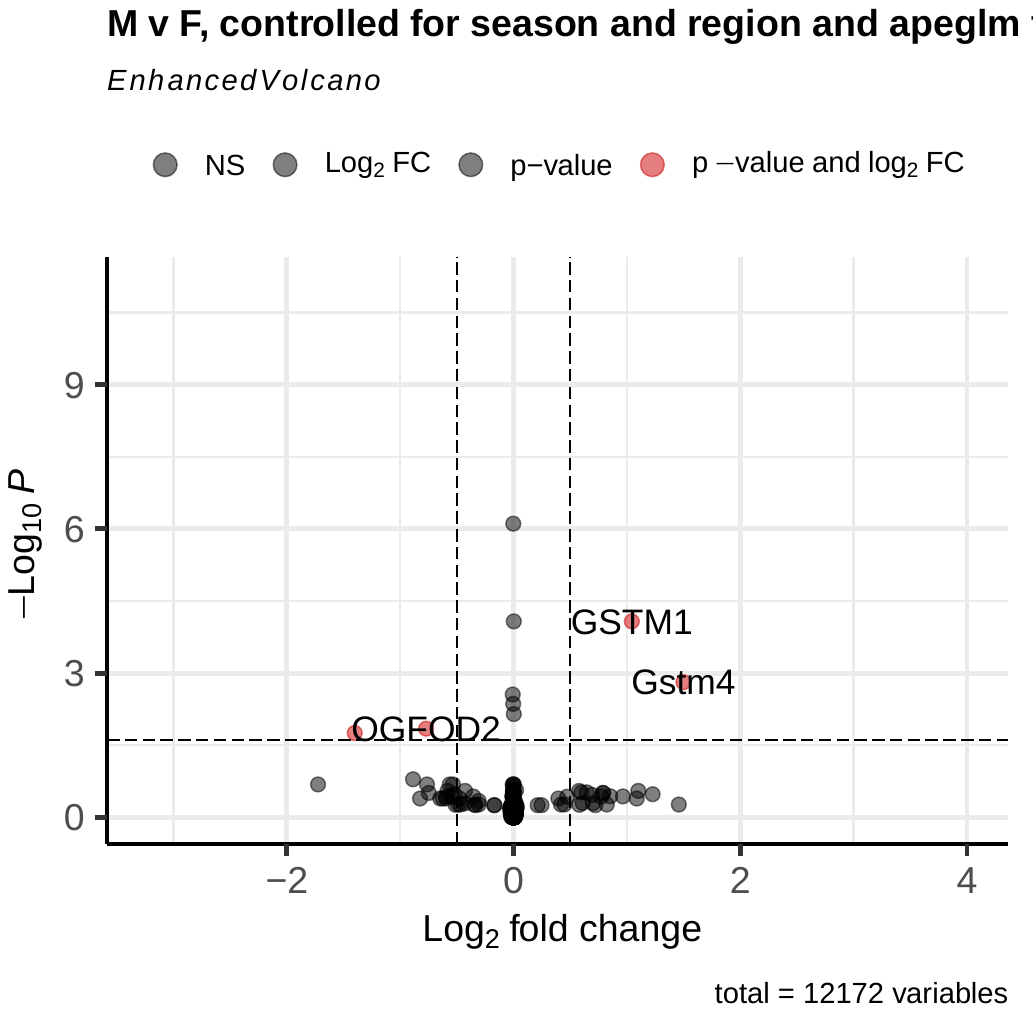

**Supp. Mat. 11. Volcano plot of genes differentially expressed between male and female lizards, controlling for regional and seasonal differences. Positive log2 fold-change values correspond to genes differentially over-expressed in males, while negative log2 fold-change values correspond to genes differentially under-expressed in the southern group (over-expressed in females).**

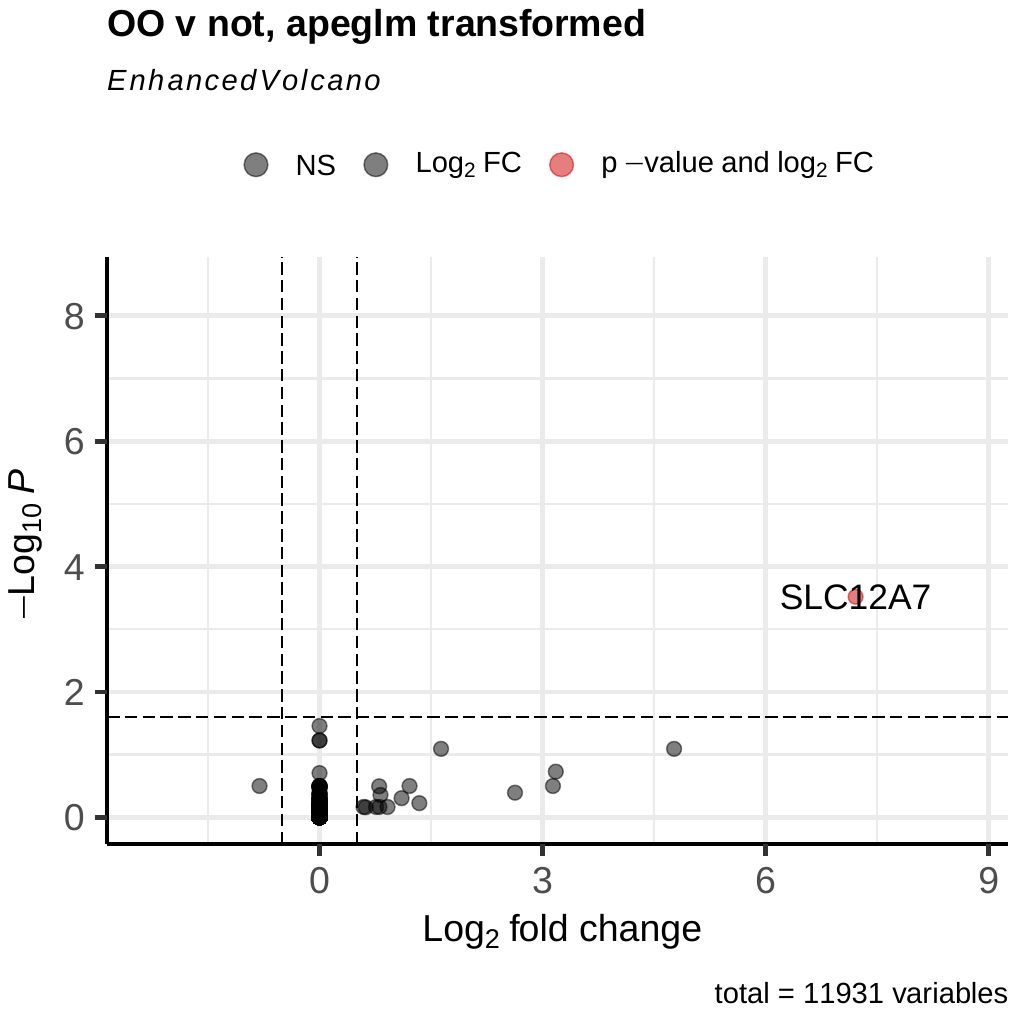

**Supp. Mat. 12. Volcano plot of genes differentially expressed between orange-throated and blue- and yellow-throated lizards. Positive log2 fold-change values correspond to genes differentially over-expressed in orange-throated lizards, while negative log2 fold-change values correspond to genes differentially under-expressed in the orange-throated lizards (over-expressed in blue- and yellow-throated lizards).**

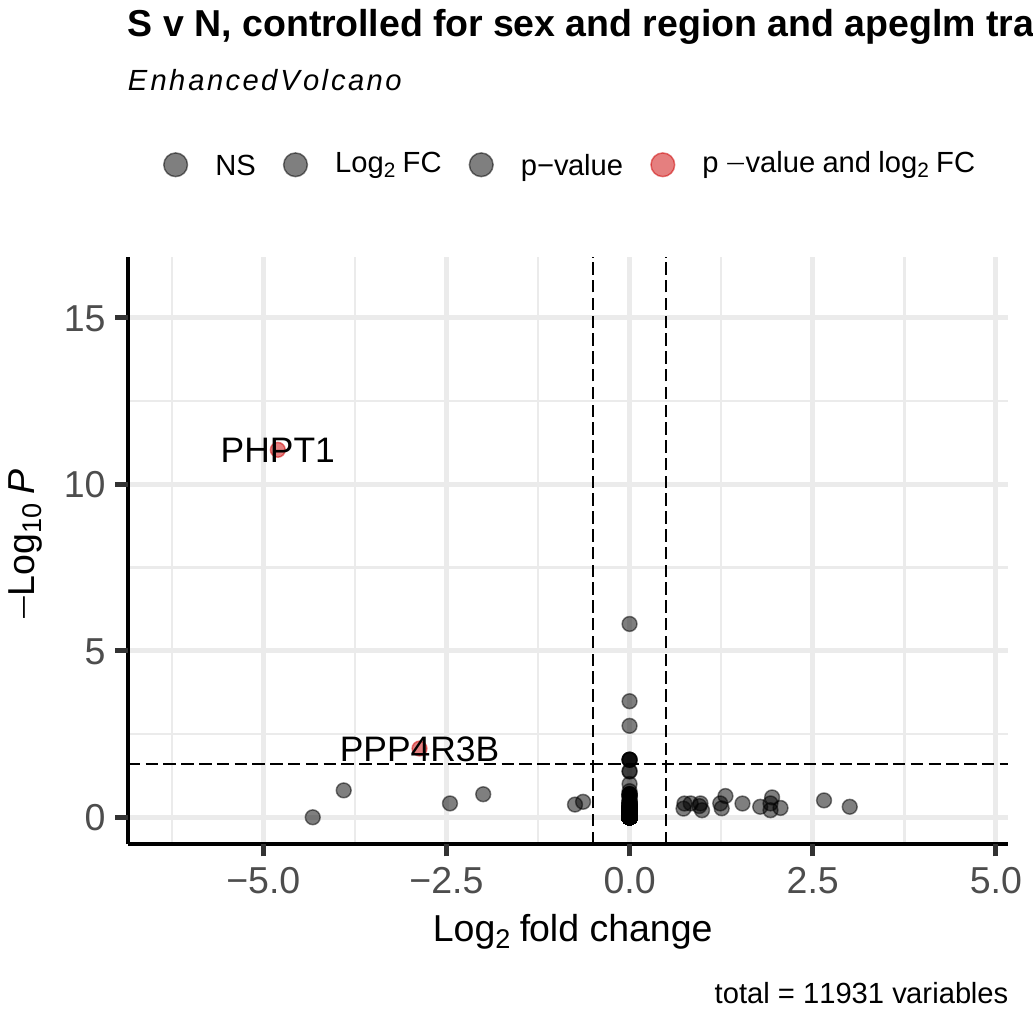

**Supp. Mat. 12. Volcano plot of genes differentially expressed between blue-throated and orange- and yellow-throated lizards. Positive log2 fold-change values correspond to genes differentially over-expressed in blue-throated lizards, while negative log2 fold-change values correspond to genes differentially under-expressed in the blue-throated lizards (over-expressed in orange- and yellow-throated lizards).**

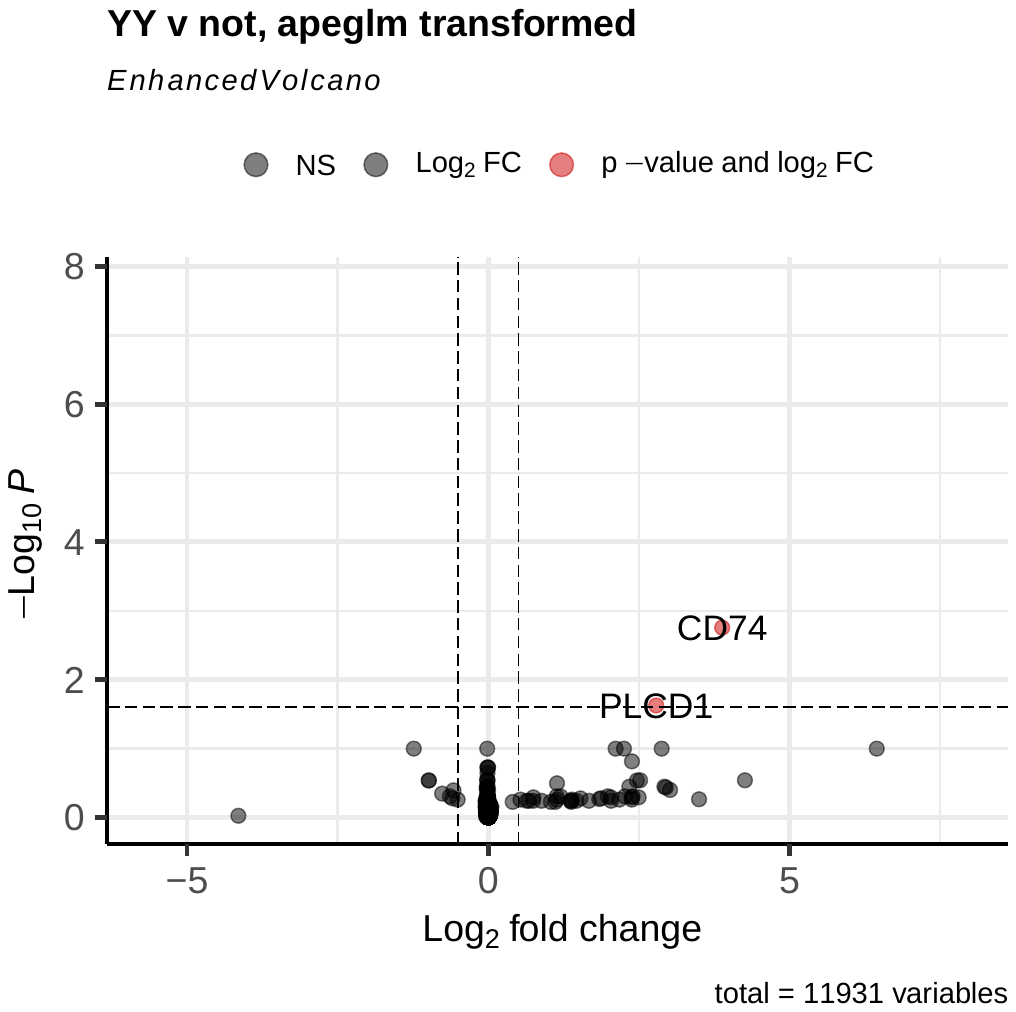

**Supp. Mat. 12. Volcano plot of genes differentially expressed between yellow-throated and blue- and orange-throated lizards. Positive log2 fold-change values correspond to genes differentially over-expressed in yellow-throated lizards, while negative log2 fold-change values correspond to genes differentially under-expressed in the yellow-throated lizards (over-expressed in blue- and orange-throated lizards).**

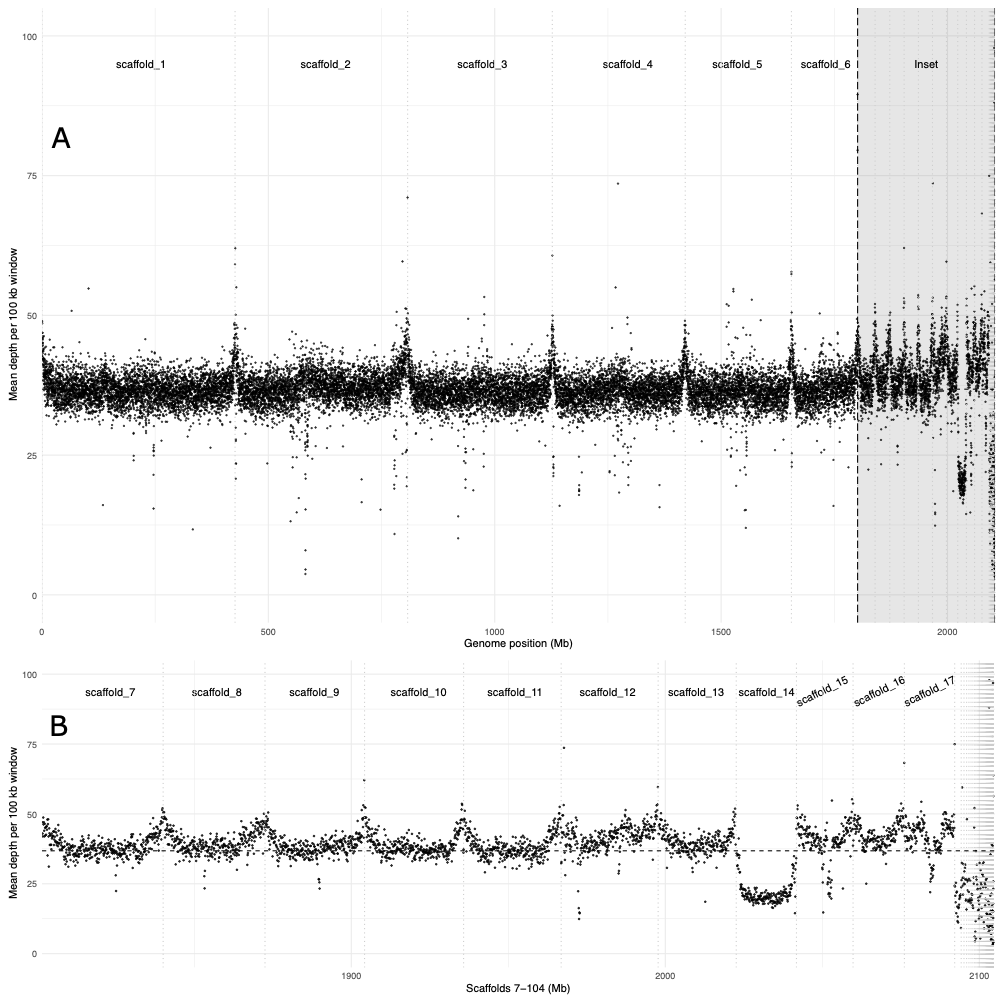

**Supp. Mat. 13. A) Plot of mean read depth of assembly-generating reads per 100 kb window across the entire genome. The genome-wide median read depth (36.80x) is plotted as a dashed line. The six chromosome-length scaffolds corresponding to the six macrochromosomes are labeled. Microchromosomes and unplaced scaffolds are highlighted in gray and labeled as an inset. B) Plot of mean read depth of assembly-generating reads per 100 kb window in microchromosomes and unplaced scaffolds. The chromosome-length scaffolds corresponding to the 11 microchromosomes (scaffold_7 through scaffold 17) are labeled, the remaining scaffolds are not. Vertical dashed lines indicate scaffold boundaries, and unlabeled scaffolds appear in descending order according to length. The genome-wide median read depth (36.80x) is plotted as a dashed line.**
