## Supplementary material for "Chromosome-length genome assembly of *Uta stansburiana* and gene expression data reveal fast pace-of-life comes with environmental stability": SuppTables.docx

**Table S1 Body size data for SSD estimation**

**Table S2 Sample list and metadata**

**Table S3 DEG list, sex contrast**

**Table S4 DEG list, OO v BB and YY contrast**

**Table S5 DEG list, BB v OO and YY contrast**

**Table S6 DEG list, YY v OO and BB contrast**

**Table S7 DEG list, region controlling for sex and season contrast**

**Table S8 GProfiler results, sex contrast**

**Table S9 GProfiler results, OO v BB and YY contrast**

**Table S10 GProfiler results, BB v OO and YY contrast**

**Table S11 GProfiler results, YY v OO and YY contrast**

**Table S12 GProfiler results, region controlling for sex and season contrast**

**Table S13 Notes regarding scaffold nomenclature differences between the annotation, DNA Zoo assembly, and GENESPACE riparian plot**

**Table S14 Mean and median read depth per 100 kb window per scaffold**

**Table S1. Body size data for SSD estimation**

| Region | Population | Latitude | Longitude | Sex | SVL |
| --- | --- | --- | --- | --- | --- |
| North | FEL | 30.731326 | -114.72227 | Female | 40 |
| North | FEL | 30.729157 | -114.72912 | Female | 37 |
| North | FEL | 30.72849 | -114.73046 | Female | 37.1 |
| North | FEL | 30.728194 | -114.73008 | Female | 39.7 |
| North | FEL | 30.731435 | -114.72192 | Male | 41.2 |
| North | FEL | 30.729036 | -114.72983 | Male | 42.1 |
| North | IGN | 27.329147 | -112.77595 | Female | 44.6 |
| North | IGN | 27.370913 | -112.6544 | Female | 40 |
| North | IGN | 27.328852 | -112.77555 | Male | 40.2 |
| North | IGN | 27.378977 | -112.67866 | Male | 45.3 |
| North | IGN | 27.379152 | -112.67977 | Male | 46 |
| North | IGN | 27.370913 | -112.6544 | Male | 43.4 |
| North | PRI | 28.833422 | -114.12165 | Female | 45.7 |
| North | PRI | 28.833282 | -114.1208 | Female | 47 |
| North | PRI | 28.833539 | -114.11696 | Male | 47.6 |
| North | PRI | 28.833282 | -114.1208 | Male | 44.9 |
| North | PRI | 28.83356 | -114.11839 | Male | 50.4 |
| North | PRI | 28.833538 | -114.11775 | Male | 45.6 |
| South | INS | 25.540114 | -111.78852 | Female | 42 |
| South | INS | 25.540114 | -111.78852 | Female | 39.6 |
| South | INS | 25.540114 | -111.78852 | Female | 44.3 |
| South | INS | 25.540079 | -111.78833 | Female | 41.2 |
| South | INS | 25.540066 | -111.78829 | Female | 45.3 |
| South | INS | 25.540066 | -111.78829 | Female | 36.3 |
| South | INS | 25.540066 | -111.78829 | Male | 46.6 |
| South | INS | 25.540095 | -111.78836 | Male | 46.3 |
| South | INS | 25.540066 | -111.78829 | Male | 51.2 |
| South | RIT | 24.579703 | -111.50666 | Female | 44 |
| South | RIT | 24.579329 | -111.50878 | Female | 38.2 |
| South | RIT | 24.579272 | -111.50832 | Female | 39.2 |
| South | RIT | 24.578802 | -111.50687 | Female | 36.4 |
| South | RIT | 24.582117 | -111.50502 | Female | 38 |
| South | RIT | 24.580201 | -111.50551 | Female | 40.6 |
| South | RIT | 24.580151 | -111.50423 | Female | 39 |
| South | RIT | 24.580231 | -111.50416 | Female | 44.5 |
| South | RIT | 24.579999 | -111.50753 | Male | 40.3 |
| South | RIT | 24.578802 | -111.50687 | Male | 43.8 |
| South | RIT | 24.579891 | -111.50652 | Male | 48.4 |

**Table S2. Sample list and associated metadata**

| Sample | region | sex | genotype | season | year |
| --- | --- | --- | --- | --- | --- |
| UstaCHA3 | N | F | BB | spr | 2022 |
| UstaCHA6 | N | M | OB | spr | 2022 |
| UstaINS2 | S | F | UNK | fal | 2021 |
| UstaINS3 | S | F | OO | fal | 2021 |
| UstaINS4 | S | M | BY | fal | 2021 |
| UstaINS5 | S | F | UNK | fal | 2021 |
| UstaINS9 | S | F | UNK | fal | 2021 |
| UstaINS10 | S | M | OO | fal | 2021 |
| UstaJAV1 | S | M | OB | fal | 2021 |
| UstaJAV4 | S | F | OO | fal | 2021 |
| UstaJAV7 | S | M | UNK | spr | 2022 |
| UstaMAG1 | S | M | OB | fal | 2021 |
| UstaMAG2 | S | M | OO | fal | 2021 |
| UstaPRI3 | N | F | UNK | fal | 2021 |
| UstaPRI5 | N | F | YY | spr | 2022 |
| UstaPRI6 | N | M | BB | spr | 2022 |
| UstaPRI7 | N | F | OO | spr | 2022 |
| UstaPRI9 | N | M | OB | spr | 2022 |
| UstaRIN1 | N | M | BB | fal | 2021 |
| UstaRIN2 | N | M | BB | spr | 2022 |
| UstaRIN3 | N | F | OO | spr | 2022 |
| UstaRIT4 | S | F | UNK | fal | 2021 |
| UstaRIT5 | S | F | UNK | fal | 2021 |
| UstaRIT7 | S | F | UNK | fal | 2021 |
| UstaRIT8 | S | F | UNK | fal | 2021 |
| UstaRIT9 | S | F | UNK | fal | 2021 |
| UstaRIT13 | S | M | YY | spr | 2022 |
| UstaRIT14 | S | F | OY | spr | 2022 |
| UstaRIT15 | S | F | UNK | spr | 2022 |
| UstaRIT18 | S | M | YY | spr | 2022 |
| UstaRIT20 | S | F | OO | spr | 2022 |
| UstaRIT22 | S | F | OY | spr | 2022 |
| UstaRIT23 | S | M | YY | spr | 2022 |

**Table S3. Differentially expressed gene list, sex contrast**

|  | baseMean | log2FoldChange | lfcSE | pvalue | padj |
| --- | --- | --- | --- | --- | --- |
| GSTM1 | 14392.4234 | 1.04265412 | 0.21032819 | 1.69E-08 | 8.40E-05 |
| Gstm4 | 474.87165 | 1.49813028 | 0.35481992 | 5.10E-07 | 0.00154451 |
| Il5ra | 79.0122722 | -0.0082699 | 0.03441599 | 1.15E-06 | 0.00278927 |
| LYZ | 53.756372 | -0.0038882 | 0.0332085 | 2.16E-06 | 0.00436778 |
| OGFOD2 | 127.471657 | -0.7724481 | 0.21630928 | 9.47E-06 | 0.01433086 |
| P2RX4 | 6016.80834 | -0.0031595 | 0.03308655 | 6.42E-11 | 7.78E-07 |
| RPL6 | 46.3665049 | 0.00133787 | 0.032903 | 2.08E-08 | 8.40E-05 |
| shfl | 38.7699912 | -1.4019534 | 0.42027434 | 1.31E-05 | 0.01762252 |
| Tmed2 | 26.0587853 | 0.0015504 | 0.03291613 | 4.11E-06 | 0.00711709 |

**Table S4. Differentially expressed gene list, OO v BB and YY contrast**

|  | baseMean | log2FoldChange | lfcSE | pvalue | padj |
| --- | --- | --- | --- | --- | --- |
| SLC12A7 | 855.5119 | 7.21016319 | 1.46289808 | 2.55E-08 | 0.00030277 |

**Table S5. Differentially expressed gene list, BB v OO and YY contrast**

|  | baseMean | log2FoldChange | lfcSE | pvalue | padj |
| --- | --- | --- | --- | --- | --- |
| ADAM20 | 3539.63293 | -6.73E-07 | 0.0014427 | 2.73E-10 | 1.56E-06 |
| ADCY7 | 2133.25371 | -1.42E-06 | 0.0014427 | 8.57E-08 | 0.00032645 |
| Cd9 | 478.004553 | -2.23E-06 | 0.0014427 | 6.27E-07 | 0.00179083 |
| EGR1 | 197.664866 | -2.99E-06 | 0.0014427 | 1.23E-05 | 0.01863681 |
| GLP2R | 26.1609325 | -2.08E-06 | 0.0014427 | 1.47E-05 | 0.01863681 |
| MSMP | 57.4443049 | -8.40E-07 | 0.0014427 | 1.39E-05 | 0.01863681 |
| PAMR1 | 1216.81426 | -1.54E-06 | 0.0014427 | 1.05E-05 | 0.01863681 |
| PHPT1 | 3281.42417 | -4.8018199 | 0.63758729 | 8.01E-16 | 9.15E-12 |
| PPP4R3B | 1116.08192 | -2.8671831 | 0.77614419 | 3.76E-06 | 0.00859917 |

**Table S6. Differentially expressed gene list, YY v OO and BB contrast**

|  | baseMean | log2FoldChange | lfcSE | pvalue | padj |
| --- | --- | --- | --- | --- | --- |
| CD74 | 277.570594 | 3.88234289 | 0.81176419 | 1.48E-07 | 0.00175927 |
| PLCD1 | 440.367279 | 2.7863508 | 0.69487073 | 4.00E-06 | 0.02378549 |

**Table S7. Differentially expressed gene list, region controlling for sex and season contrast**

|  | baseMean | log2FoldChange | lfcSE | pvalue | padj |
| --- | --- | --- | --- | --- | --- |
| TAAR1 | 146.440915 | -9.4043138 | 0.85803401 | 8.81E-28 | 2.67E-24 |
| Aox3 | 97.4167356 | -7.955788 | 1.08715271 | 2.28E-20 | 2.51E-17 |
| KCNH7 | 34.6725813 | -5.954778 | 0.56879186 | 1.44E-25 | 2.90E-22 |
| slc7a2 | 61.9329029 | -5.7437452 | 0.58137606 | 5.61E-22 | 7.55E-19 |
| CDIPT | 52.1787425 | -5.3629823 | 0.51601061 | 1.45E-22 | 2.51E-19 |
| TIE1 | 50.7161164 | -4.7896045 | 0.55360924 | 5.15E-18 | 5.20E-15 |
| zdhhc11 | 30.2306054 | -3.7647667 | 0.50349372 | 1.99E-15 | 1.50E-12 |
| Esm1 | 13.2544953 | -3.6736418 | 1.05702605 | 7.92E-07 | 0.00018435 |
| SLC2A9 | 8.70214501 | -3.0633413 | 0.96801801 | 1.53E-05 | 0.00227956 |
| INTU | 222.485295 | -2.8618944 | 0.42616295 | 2.58E-12 | 1.36E-09 |
| WDR49 | 23.2511141 | -2.3334819 | 0.67609425 | 0.00010306 | 0.01075591 |
| CALR | 1790.78138 | -2.1221975 | 0.25274922 | 8.57E-18 | 7.98E-15 |
| Faah | 28.8733317 | -2.0767318 | 0.78501793 | 0.00015715 | 0.01486405 |
| TTLL11 | 104.243468 | -2.0300526 | 0.31561262 | 3.50E-13 | 2.02E-10 |
| Clec5a | 24.6217356 | -2.0214817 | 0.68635015 | 6.03E-05 | 0.00722669 |
| Dvl1 | 69.4267872 | -1.9931571 | 0.41624875 | 8.57E-09 | 3.14E-06 |
| haus8 | 417.835963 | -1.9028686 | 0.28955838 | 1.18E-12 | 6.49E-10 |
| Slc25a22 | 629.121476 | -1.892598 | 0.70651321 | 6.96E-05 | 0.00802504 |
| ncor1 | 58.7942869 | -1.8835791 | 0.2911791 | 1.67E-11 | 8.41E-09 |
| Tap2 | 23.2857837 | -1.8454789 | 0.6468936 | 0.0002234 | 0.0191824 |
| ATp6v1fnb | 25.0592466 | -1.793344 | 0.58628946 | 0.00012009 | 0.01221781 |
| Elavl2 | 38.7364648 | -1.7665095 | 0.74880426 | 0.00025639 | 0.02087596 |
| Hspg2 | 11.8006699 | -1.7517556 | 0.5752375 | 6.25E-05 | 0.00741567 |
| CA2 | 1813.03367 | -1.7411886 | 0.31058495 | 6.43E-09 | 2.51E-06 |
| Tbc1d7 | 399.130532 | -1.7397146 | 0.37199356 | 1.27E-08 | 4.38E-06 |
| CHI3L1 | 182.546483 | -1.7278756 | 0.57392961 | 0.00013849 | 0.01359148 |
| CPN2 | 57.2072321 | -1.72275 | 0.52405236 | 0.00015132 | 0.01454014 |
| CASR | 9.42042731 | -1.6719887 | 0.54721815 | 0.00020526 | 0.018139 |
| WASF3 | 324.890734 | -1.6463795 | 0.39857898 | 2.02E-06 | 0.00042983 |
| FOS | 14.9442844 | -1.5931293 | 0.57442229 | 6.33E-05 | 0.00743514 |
| ACP6 | 87.2550911 | -1.5636053 | 0.45708719 | 2.26E-05 | 0.003081 |
| MTHFD2 | 69.0165819 | -1.5307645 | 0.35455636 | 8.54E-07 | 0.00019512 |
| SPTBN5 | 79.4631748 | -1.5173149 | 0.41851336 | 1.23E-05 | 0.00186214 |
| H2B-VII | 85.1167496 | -1.4807928 | 0.44780989 | 0.0002033 | 0.01809845 |
| PSMA4 | 3110.46602 | -1.3864444 | 0.18810528 | 3.81E-14 | 2.71E-11 |
| AGPAT5 | 137.849552 | -1.2926247 | 0.24329983 | 6.07E-09 | 2.45E-06 |
| FAM131C | 95.7579827 | -1.2789989 | 0.42786896 | 0.00025669 | 0.02087596 |
| Cdk5rap1 | 68.8609731 | -1.2725557 | 0.34370626 | 1.07E-05 | 0.00166279 |
| RNF213 | 122.819621 | -1.219472 | 0.43999541 | 0.0002497 | 0.02081978 |
| METTL9 | 103.266982 | -1.176161 | 0.27308906 | 3.40E-07 | 8.95E-05 |
| Gkap1 | 766.123896 | -1.1492647 | 0.29783675 | 2.67E-06 | 0.00053853 |
| GSTO1 | 331.916055 | -1.0113518 | 0.37203654 | 0.00015433 | 0.01471221 |
| GATAD1 | 168.456088 | -1.0088545 | 0.25988098 | 4.65E-06 | 0.00086668 |
| MGST1 | 133.950014 | -0.9987764 | 0.41211902 | 0.00025692 | 0.02087596 |
| TRIM27 | 408.243678 | -0.9743642 | 0.2108217 | 6.54E-07 | 0.00016153 |
| Cdk2ap2 | 177.225396 | -0.9734087 | 0.29194242 | 3.27E-05 | 0.00416635 |
| CCND1 | 227.277269 | -0.9668017 | 0.2560641 | 1.01E-05 | 0.00162323 |
| brcc3 | 119.515139 | -0.9519643 | 0.21488268 | 4.15E-07 | 0.00010697 |
| Eef1akmt2 | 258.37261 | -0.9329358 | 0.27778405 | 3.44E-05 | 0.00433532 |
| CCNL1 | 297.69512 | -0.9066402 | 0.31060835 | 5.30E-05 | 0.00648152 |
| Prkrip1 | 58.4336932 | -0.8922307 | 0.35749079 | 0.00026263 | 0.02105721 |
| TMEM165 | 1074.25159 | -0.8494724 | 0.2327666 | 5.88E-06 | 0.00098913 |
| YBX1 | 80396.2126 | -0.7699991 | 0.30648345 | 0.00023482 | 0.01979863 |
| Rpf2 | 271.70571 | -0.7685355 | 0.21069169 | 2.45E-05 | 0.00330155 |
| mcts1-a | 1966.27545 | -0.7414905 | 0.26415827 | 0.00017055 | 0.01576215 |
| Amacr | 214.541428 | -0.7297279 | 0.26749799 | 0.00026986 | 0.02149483 |
| SGK3 | 536.074049 | -0.6990374 | 0.27272521 | 0.00028873 | 0.02240819 |
| K-RAS | 793.833421 | -0.6483743 | 0.15605639 | 2.35E-06 | 0.00048222 |
| Spcs2 | 514.565255 | -0.6423889 | 0.22600954 | 0.00029982 | 0.02283815 |
| TFAM | 383.344049 | -0.6115011 | 0.19467369 | 0.00012999 | 0.0130062 |
| GTF2H3 | 246.341292 | -0.5810318 | 0.19858478 | 0.00030182 | 0.02283815 |
| TMEM33 | 240.362066 | -0.5796343 | 0.20452983 | 0.0002171 | 0.01890957 |
| PSIP1 | 863.431426 | -0.573218 | 0.18479945 | 0.00012607 | 0.01271888 |
| ALDH7A1 | 1506.70414 | -0.5198379 | 0.16130651 | 7.77E-05 | 0.00888 |
| VPS25 | 496.758613 | -0.4642629 | 0.15106439 | 0.00023548 | 0.01979863 |
| Foxa3 | 3.23573786 | -0.1812697 | 0.36346625 | 1.68E-06 | 0.00036233 |
| ADGRE3 | 5.12363145 | -0.1245084 | 0.25021033 | 0.00014033 | 0.01359148 |
| Morn4 | 423.118898 | 0.45688679 | 0.15469384 | 0.00030037 | 0.02283815 |
| Sf3a3 | 1385.0882 | 0.55740382 | 0.19257176 | 0.00026041 | 0.02101876 |
| MRPL16 | 429.849578 | 0.5613028 | 0.15560307 | 1.79E-05 | 0.00251839 |
| DUSP16 | 354.23292 | 0.65087215 | 0.24067136 | 0.00031221 | 0.0233602 |
| VCL | 1335.49261 | 0.66820638 | 0.21307338 | 0.00011534 | 0.01183392 |
| DYRK1A | 305.927372 | 0.68010875 | 0.23169018 | 0.00017511 | 0.01606124 |
| HELZ | 223.160595 | 0.69074066 | 0.22205485 | 8.91E-05 | 0.00954674 |
| Anp32e | 1703.656 | 0.73785943 | 0.13006357 | 4.66E-10 | 2.01E-07 |
| tars3 | 2216.18255 | 0.75106886 | 0.20453144 | 1.10E-05 | 0.00168678 |
| TIPRL | 443.70731 | 0.75183369 | 0.1992019 | 5.80E-06 | 0.00098913 |
| Mydgf | 959.361836 | 0.77407716 | 0.21868885 | 1.85E-05 | 0.00257474 |
| FRA10AC1 | 82.1863494 | 0.82195732 | 0.29167287 | 0.00023278 | 0.01979863 |
| RAPGEF4 | 136.016777 | 0.86230168 | 0.30113695 | 0.00013968 | 0.01359148 |
| PBDC1 | 224.308024 | 0.93671383 | 0.23583599 | 3.52E-06 | 0.00068683 |
| SNX29 | 63.3509731 | 0.96749574 | 0.34503114 | 0.00027918 | 0.02209188 |
| MTIF2 | 275.929714 | 1.07876687 | 0.21493785 | 1.77E-08 | 5.80E-06 |
| NMRK2 | 610.817866 | 1.08153725 | 0.41289568 | 0.00019517 | 0.01776667 |
| SPR | 182.194053 | 1.10037919 | 0.43088288 | 0.00022183 | 0.0191824 |
| RIOK2 | 331.89391 | 1.15911955 | 0.23025669 | 1.90E-08 | 6.07E-06 |
| hoxc11a | 89.7467087 | 1.17295336 | 0.33451799 | 9.17E-06 | 0.00150067 |
| DLL1 | 122.698087 | 1.17363891 | 0.36517697 | 2.92E-05 | 0.00380038 |
| LMNA | 1759.49806 | 1.17813536 | 0.28063786 | 1.10E-06 | 0.00024603 |
| EGFR | 141.445284 | 1.19108373 | 0.33268325 | 1.02E-05 | 0.00162471 |
| sepp1b | 245.847991 | 1.2213214 | 0.47618006 | 0.00031258 | 0.0233602 |
| METTL27 | 39.8554432 | 1.22403766 | 0.40485959 | 8.88E-05 | 0.00954674 |
| ATXN10 | 169.979111 | 1.23249741 | 0.34134617 | 4.29E-06 | 0.00082026 |
| MRTO4 | 56.6482728 | 1.23611694 | 0.27901537 | 2.91E-07 | 7.82E-05 |
| Notum | 226.682869 | 1.23875935 | 0.37024285 | 2.53E-05 | 0.00337147 |
| SVEP1 | 44.198134 | 1.244284 | 0.37345576 | 2.86E-05 | 0.00376578 |
| CREM | 679.510718 | 1.26364184 | 0.34442429 | 5.41E-06 | 0.00094845 |
| Scn2b | 155.980751 | 1.27816662 | 0.34065614 | 1.06E-05 | 0.00166279 |
| nr2f1a | 32.25482 | 1.30034662 | 0.44499128 | 8.34E-05 | 0.00935129 |
| Xirp1 | 1805.09794 | 1.3066781 | 0.47092479 | 0.00016013 | 0.01502852 |
| SIAH1 | 88.0050211 | 1.32735366 | 0.50205142 | 8.64E-05 | 0.00953838 |
| RC3H2 | 53.1919645 | 1.34293523 | 0.39032009 | 1.58E-05 | 0.00233564 |
| Smarcd3 | 76.8499554 | 1.39083527 | 0.39992356 | 4.00E-05 | 0.00499116 |
| CCDC136 | 28.1905526 | 1.39969071 | 0.51244191 | 0.00013374 | 0.0132718 |
| ANKRD37 | 60.6111849 | 1.42922175 | 0.58227921 | 0.00025107 | 0.02081978 |
| CSTPP1 | 171.290019 | 1.4684545 | 0.38196089 | 5.36E-06 | 0.00094845 |
| Asb12 | 202.607276 | 1.48237957 | 0.50996718 | 0.00010858 | 0.01123556 |
| ALPK3 | 67.9313285 | 1.50736621 | 0.46413125 | 9.09E-05 | 0.00965509 |
| WHAMM | 323.236567 | 1.55204636 | 0.43487631 | 4.34E-06 | 0.00082026 |
| ZZEF1 | 235.504122 | 1.60327548 | 0.3201028 | 8.21E-09 | 3.11E-06 |
| GPRC5A | 153.503183 | 1.60401295 | 0.37720596 | 7.36E-07 | 0.00017464 |
| Ralgps1 | 60.2948423 | 1.62666969 | 0.46841179 | 2.97E-06 | 0.00058896 |
| JPH4 | 19.6142851 | 1.63032322 | 0.49816593 | 3.24E-05 | 0.00416635 |
| AGO3 | 185.542116 | 1.65137261 | 0.49706489 | 9.26E-05 | 0.00974664 |
| S100A10 | 202.89253 | 1.67820748 | 0.32325366 | 2.99E-08 | 9.06E-06 |
| PYROXD2 | 84.7173316 | 1.7103591 | 0.65778269 | 8.78E-05 | 0.00954674 |
| Zcchc3 | 64.3709865 | 1.71254223 | 0.69815552 | 0.00016537 | 0.01540084 |
| Abtb3 | 28.831793 | 1.76288658 | 0.36795629 | 1.36E-07 | 3.83E-05 |
| PFKFB3 | 53.754765 | 1.80469788 | 0.7019387 | 0.00028434 | 0.02220947 |
| FXYD1 | 1546.59461 | 1.81730743 | 0.26369241 | 7.99E-14 | 5.37E-11 |
| FAM110B | 19.7349748 | 1.86267943 | 0.82574955 | 0.00019717 | 0.01781465 |
| ACSBG2 | 166.82028 | 1.88062246 | 0.54688078 | 1.69E-05 | 0.00242877 |
| EP400 | 48.1027246 | 1.90214321 | 0.3515322 | 4.55E-09 | 1.90E-06 |
| ATF3 | 151.640117 | 1.91661353 | 0.58871486 | 4.83E-06 | 0.00088547 |
| NR4A3 | 252.231857 | 1.94732588 | 0.73561537 | 4.95E-05 | 0.00611564 |
| LRCH3 | 13.2796095 | 1.96616406 | 0.73123725 | 8.67E-05 | 0.00953838 |
| Tspan7 | 36.1003387 | 2.05697544 | 0.59442871 | 2.00E-05 | 0.00274586 |
| Aoc1 | 16.5728818 | 2.06533129 | 0.83647952 | 0.00021567 | 0.01890957 |
| SLC9A6 | 128.001 | 2.07589839 | 0.2675367 | 2.02E-17 | 1.63E-14 |
| OTUD7B | 28.5083399 | 2.12579316 | 0.57541882 | 5.29E-06 | 0.00094845 |
| Fbxo41 | 16.693956 | 2.17566676 | 0.74817703 | 6.00E-05 | 0.00722669 |
| CEP19 | 61.3803932 | 2.22630868 | 0.55770026 | 1.21E-06 | 0.00026592 |
| NEU3 | 610.973897 | 2.26086464 | 0.33943496 | 1.68E-13 | 1.05E-10 |
| tmem45b | 1061.06149 | 2.30386613 | 0.44309841 | 1.62E-08 | 5.45E-06 |
| ARHGAP39 | 5.24294251 | 2.45590078 | 0.93587199 | 0.00019956 | 0.01789649 |
| Iqsec2 | 17.3159673 | 2.47359713 | 0.75844659 | 1.64E-05 | 0.00239263 |
| Pprc1 | 668.599826 | 2.56736673 | 0.26906812 | 3.52E-22 | 5.32E-19 |
| CHRDL1 | 162.493401 | 2.63954569 | 0.44546577 | 2.25E-10 | 1.01E-07 |
| HMBOX1 | 138.632904 | 2.77053646 | 1.11153493 | 1.78E-05 | 0.00251839 |
| PPP4R3B | 1288.49152 | 2.79223624 | 0.38253439 | 1.30E-17 | 1.12E-14 |
| CLIC6 | 19.4714686 | 2.83030902 | 1.00685748 | 6.57E-05 | 0.00764708 |
| PDE6C | 4.35213116 | 2.89580338 | 1.08370039 | 0.00028183 | 0.02215638 |
| SERINC4 | 13.2363237 | 2.96687404 | 0.77834409 | 5.79E-06 | 0.00098913 |
| CPNE4 | 27.9428017 | 3.07246856 | 0.60790966 | 4.92E-08 | 1.45E-05 |
| LRRC74B | 15.1789112 | 3.21496449 | 0.67375006 | 7.74E-08 | 2.23E-05 |
| EGR1 | 194.248242 | 3.22244239 | 0.50726924 | 1.73E-13 | 1.05E-10 |
| ZC3H12C | 28.4022878 | 3.26408944 | 0.79404521 | 2.20E-07 | 6.06E-05 |
| SSPO | 71.3608048 | 3.28803359 | 1.10013274 | 8.14E-06 | 0.00134976 |
| GLP2R | 23.3830242 | 3.39066993 | 0.85530221 | 1.23E-08 | 4.38E-06 |
| PIWIL1 | 66.775691 | 3.43445703 | 0.64137124 | 1.51E-10 | 7.04E-08 |
| ZBTB8OS | 8.77523175 | 3.65490453 | 0.83182372 | 4.92E-07 | 0.00012409 |
| UROC1 | 7.58116618 | 4.13695897 | 1.51064733 | 0.00029821 | 0.02283815 |
| GABRA5 | 3.73813409 | 4.17334966 | 1.4651219 | 8.24E-05 | 0.00932659 |
| Cd9 | 541.013111 | 4.39187503 | 0.47942741 | 2.97E-27 | 7.19E-24 |
| LGSN | 9.46394719 | 4.44502313 | 1.15208255 | 7.02E-07 | 0.0001699 |
| CDHR4 | 167.132883 | 4.5290633 | 0.87132026 | 7.67E-11 | 3.72E-08 |
| TOPAZ1 | 69.3836859 | 4.7749413 | 0.95309559 | 2.25E-08 | 7.00E-06 |
| ADCY7 | 2642.72348 | 6.14966727 | 0.55824695 | 1.58E-32 | 6.37E-29 |
| PAMR1 | 1225.26648 | 7.02395582 | 0.61683412 | 6.29E-34 | 3.81E-30 |
| MSMP | 55.80676 | 7.66068533 | 0.91042971 | 9.47E-21 | 1.15E-17 |
| PRKCB | 174.047426 | 7.92580785 | 2.30843059 | 2.23E-06 | 0.00046604 |
| ADAM20 | 3297.78569 | 11.4258141 | 0.54408946 | 2.36E-107 | 2.86E-103 |

**Table S8. GProfiler results, sex contrast**

| source | term_name | term_id | highlighted | adjusted_p_value | negative_log10_of_adjusted_p_value | term_size | query_size | intersection_size | effective_domain_size | intersections |
| --- | --- | --- | --- | --- | --- | --- | --- | --- | --- | --- |
| GO:MF | glutathione binding | GO:0043295 | TRUE | 0.00263747 | 2.5788118674926075 | 20 | 8 | 2 | 25071 | GSTM1,GSTM4 |
| GO:MF | oligopeptide binding | GO:1900750 | FALSE | 0.00291464 | 2.5354154875195913 | 21 | 8 | 2 | 25071 | GSTM1,GSTM4 |
| GO:MF | glutathione transferase activity | GO:0004364 | TRUE | 0.00687202 | 2.162915603093053 | 32 | 8 | 2 | 25071 | GSTM1,GSTM4 |
| GO:MF | transferase activity, transferring alkyl or aryl (other than methyl) groups | GO:0016765 | FALSE | 0.028666723976079424 | 1.542621935250391 | 65 | 8 | 2 | 25071 | GSTM1,GSTM4 |
| GO:MF | interleukin-5 receptor activity | GO:0004914 | TRUE | 0.0498584 | 1.302261643878731 | 1 | 8 | 1 | 25071 | IL5RA |
| GO:BP | long-chain fatty acid biosynthetic process | GO:0042759 | TRUE | 0.015160382814495365 | 1.8192898322050293 | 25 | 7 | 2 | 26963 | GSTM1,GSTM4 |
| GO:BP | xenobiotic catabolic process | GO:0042178 | TRUE | 0.021968965139618043 | 1.6581904002819785 | 30 | 7 | 2 | 26963 | GSTM1,GSTM4 |
| KEGG | Drug metabolism - cytochrome P450 | KEGG:00982 | FALSE | 0.02918067 | 1.534904813890528 | 71 | 5 | 2 | 9330 | GSTM1,GSTM4 |
| KEGG | Glutathione metabolism | KEGG:00480 | FALSE | 0.03000794 | 1.5227638286973917 | 72 | 5 | 2 | 9330 | GSTM1,GSTM4 |
| KEGG | Metabolism of xenobiotics by cytochrome P450 | KEGG:00980 | FALSE | 0.03169662 | 1.4989870514630435 | 74 | 5 | 2 | 9330 | GSTM1,GSTM4 |
| KEGG | Platinum drug resistance | KEGG:01524 | FALSE | 0.037035179542163815 | 1.4313855455990745 | 80 | 5 | 2 | 9330 | GSTM1,GSTM4 |
| KEGG | Chemical carcinogenesis - DNA adducts | KEGG:05204 | FALSE | 0.04082072 | 1.3891193406435982 | 84 | 5 | 2 | 9330 | GSTM1,GSTM4 |
| KEGG | Drug metabolism - other enzymes | KEGG:00983 | FALSE | 0.046837586044417734 | 1.3294054960634727 | 90 | 5 | 2 | 9330 | GSTM1,GSTM4 |
| REAC | Glutathione conjugation | REAC:R-MMU-156590 | FALSE | 0.026189891608298214 | 1.5818662989789443 | 37 | 5 | 2 | 8429 | GSTM1,GSTM4 |
| WP | Oxidative stress and redox pathway | WP:WP4466 | FALSE | 0.044941555638649676 | 1.3473518986817248 | 92 | 4 | 2 | 4532 | GSTM1,GSTM4 |

**Table S9. GProfiler results, OO v BB and YY contrast**

| source | term_name | term_id | highlighted | adjusted_p_value | negative_log10_of_adjusted_p_value | term_size | query_size | intersection_size | effective_domain_size | intersections |
| --- | --- | --- | --- | --- | --- | --- | --- | --- | --- | --- |
| KEGG | Collecting duct acid secretion | KEGG:04966 | FALSE | 0.04989467 | 1.30194587 | 27 | 1 | 1 | 9330 | SLC12A7 |

**Table S10. GProfiler results, BB v OO and YY contrast**

| source | term_name | term_id | highlighted | adjusted_p_value | negative_log10_of_adjusted_p_value | term_size | query_size | intersection_size | effective_domain_size | intersections |
| --- | --- | --- | --- | --- | --- | --- | --- | --- | --- | --- |
| GO:MF | protein histidine phosphatase activity | GO:0101006 | TRUE | 0.04999728 | 1.30105359 | 1 | 9 | 1 | 25071 | PHPT1 |
| KEGG | GnRH signaling pathway | KEGG:04912 | FALSE | 0.04683759 | 1.3294055 | 90 | 5 | 2 | 9330 | ADCY7,EGR1 |
| MIRNA | mmu-miR-7578 | MIRNA:mmu-miR-7578 | FALSE | 0.04998765 | 1.30113725 | 1 | 2 | 1 | 7222 | EGR1 |

**Table S11. GProfiler results, YY v OO and BB contrast**

| source | term_name | term_id | highlighted | adjusted_p_value | negative_log10_of_adjusted_p_value | term_size | query_size | intersection_size | effective_domain_size | intersections |
| --- | --- | --- | --- | --- | --- | --- | --- | --- | --- | --- |
| GO:MF | MHC class II protein binding, via antigen binding groove | GO:0042658 | TRUE | 0.0498584 | 1.30226164 | 1 | 2 | 1 | 25071 | CD74 |
| GO:BP | positive regulation of small molecule metabolic process | GO:0062013 | TRUE | 0.03020914 | 1.51986161 | 182 | 2 | 2 | 26963 | CD74,PLCD1 |
| GO:CC | NOS2-CD74 complex | GO:0035693 | TRUE | 0.01663867 | 1.77888134 | 1 | 2 | 1 | 27195 | CD74 |
| GO:CC | macrophage migration inhibitory factor receptor complex | GO:0035692 | TRUE | 0.03327673 | 1.47785933 | 2 | 2 | 1 | 27195 | CD74 |
| TF | Factor: NF-kappaB; motif: NGGGANTTYCCMNNNN; match class: 1 | TF:M00774_1 | FALSE | 0.00695586 | 2.15764887 | 135 | 2 | 2 | 21628 | CD74,PLCD1 |
| TF | Factor: NF-kappaB; motif: NGGGGAMTTTCCNN; match class: 1 | TF:M00194_1 | FALSE | 0.04328094 | 1.36370335 | 336 | 2 | 2 | 21628 | CD74,PLCD1 |
| CORUM | Cd74-Cd44 receptor complex | CORUM:3226 | FALSE | 0.02497877 | 1.60242898 | 1 | 1 | 1 | 1082 | CD74 |

**Table S12. GProfiler results, region controlling for sex and season contrast**

| source | term_name | term_id | highlighted | adjusted_p_value | negative_log10_of_adjusted_p_value | term_size | query_size | intersection_size | effective_domain_size | intersections |
| --- | --- | --- | --- | --- | --- | --- | --- | --- | --- | --- |
| GO:MF | protein binding | GO:0005515 | TRUE | 0.00243045 | 2.61431254 | 10472 | 144 | 87 | 25071 | AOX3,FOXA3,TIE1,ESM1,INTU,WDR49,FAAH,CLEC5A,TAP2,CALR,DVL1,TTLL11,HSPG2,CHI3L1,CPN2,HAUS8,CASR,NCOR1,FOS,TBC1D7,CA2,WASF3,SPTBN5,GKAP1,MGST1,CDK2AP2,PRKRIP1,CCND1,CCNL1,TRIM27,BRCC3,YBX1,AMACR,TFAM,PSIP1,ALDH7A1,VPS25,MORN4,VCL,ANP32E,DUSP16,DYRK1A,HELZ,RAPGEF4,LMNA,NMRK2,EGFR,DLL1,ATXN10,CREM,SVEP1,RC3H2,XIRP1,SMARCD3,ZZEF1,ASB12,WHAMM,ANKRD37,S100A10,FXYD1,AGO3,ABTB3,EP400,SLC9A6,ATF3,NR4A3,LRCH3,OTUD7B,FBXO41,PPRC1,CHRDL1,IQSEC2,PPP4R3B,ARHGAP39,CPNE4,EGR1,CLIC6,HMBOX1,LRRC74B,PDE6C,PIWIL1,SSPO,LGSN,GABRA5,CD9,MSMP,PRKCB |
| GO:MF | catalytic activity | GO:0003824 | TRUE | 0.00564479 | 2.24835247 | 5731 | 144 | 56 | 25071 | AOX3,CDIPT,TIE1,ZDHHC11,FAAH,TAP2,TTLL11,CA2,ACP6,MTHFD2,RNF213,CDK5RAP1,AGPAT5,METTL9,MGST1,GSTO1,CCND1,EEF1AKMT2,TRIM27,BRCC3,AMACR,SGK3,SPCS2,ALDH7A1,DUSP16,DYRK1A,HELZ,TARS3,FRA10AC1,MTIF2,RIOK2,NMRK2,EGFR,SPR,NOTUM,RC3H2,ZZEF1,ALPK3,ASB12,AGO3,EP400,PYROXD2,ACSBG2,PFKFB3,NEU3,OTUD7B,AOC1,PDE6C,ZC3H12C,PIWIL1,UROC1,LGSN,ADCY7,PAMR1,PRKCB,ADAM20 |
| GO:BP | regulation of response to stimulus | GO:0048583 | TRUE | 0.00751164 | 2.12426541 | 4015 | 150 | 44 | 26963 | TAAR1,SLC7A2,ESM1,ZDHHC11,INTU,TAP2,CALR,DVL1,CHI3L1,CASR,NCOR1,TBC1D7,GKAP1,TRIM27,BRCC3,RPF2,SGK3,TMEM33,DUSP16,DYRK1A,MYDGF,LMNA,EGFR,DLL1,NOTUM,SVEP1,RC3H2,SMARCD3,RALGPS1,AGO3,EP400,ZCCHC3,SLC9A6,ATF3,NEU3,NR4A3,OTUD7B,CHRDL1,IQSEC2,PPP4R3B,EGR1,CD9,ADCY7,PRKCB |
| GO:BP | regulation of multicellular organismal process | GO:0051239 | TRUE | 0.02903715 | 1.53704596 | 3145 | 150 | 36 | 26963 | SLC7A2,FOXA3,TIE1,INTU,FAAH,CLEC5A,CALR,DVL1,CHI3L1,CASR,FOS,CA2,WASF3,CCND1,TRIM27,VCL,MYDGF,RAPGEF4,LMNA,EGFR,DLL1,NOTUM,SVEP1,SCN2B,RC3H2,SMARCD3,S100A10,JPH4,FXYD1,ZCCHC3,NR4A3,OTUD7B,EGR1,CD9,ADCY7,PRKCB |
| GO:CC | cytoplasm | GO:0005737 | TRUE | 2.42E-07 | 6.61696404 | 11701 | 149 | 101 | 27195 | AOX3,CDIPT,FOXA3,ZDHHC11,INTU,FAAH,CLEC5A,SLC25A22,TAP2,CALR,DVL1,TTLL11,CHI3L1,HAUS8,NCOR1,FOS,TBC1D7,CA2,WASF3,ACP6,SPTBN5,MTHFD2,RNF213,CDK5RAP1,PSMA4,AGPAT5,METTL9,GKAP1,MGST1,GSTO1,CDK2AP2,CCND1,CCNL1,EEF1AKMT2,TRIM27,BRCC3,TMEM165,YBX1,AMACR,SGK3,SPCS2,TFAM,TMEM33,ALDH7A1,VPS25,MORN4,MRPL16,VCL,ANP32E,DUSP16,DYRK1A,HELZ,TIPRL,TARS3,MYDGF,RAPGEF4,MTIF2,RIOK2,LMNA,NMRK2,EGFR,SPR,MRTO4,DLL1,ATXN10,CREM,SVEP1,RC3H2,SMARCD3,CSTPP1,WHAMM,ANKRD37,S100A10,RALGPS1,JPH4,AGO3,PYROXD2,ZCCHC3,SLC9A6,ACSBG2,PFKFB3,NEU3,FAM110B,LRCH3,OTUD7B,TMEM45B,CEP19,IQSEC2,PPP4R3B,ARHGAP39,EGR1,CLIC6,HMBOX1,ZC3H12C,PIWIL1,UROC1,LGSN,GABRA5,TOPAZ1,MSMP,PRKCB |
| GO:CC | cellular anatomical structure | GO:0110165 | FALSE | 0.00080893 | 3.09209042 | 22771 | 149 | 143 | 27195 | TAAR1,AOX3,KCNH7,SLC7A2,CDIPT,FOXA3,TIE1,ESM1,ZDHHC11,SLC2A9,INTU,FAAH,CLEC5A,SLC25A22,ELAVL2,TAP2,CALR,DVL1,TTLL11,HSPG2,CHI3L1,CPN2,HAUS8,CASR,NCOR1,FOS,TBC1D7,CA2,WASF3,ACP6,SPTBN5,MTHFD2,RNF213,CDK5RAP1,PSMA4,AGPAT5,METTL9,GKAP1,MGST1,GSTO1,GATAD1,CDK2AP2,PRKRIP1,CCND1,CCNL1,EEF1AKMT2,TRIM27,BRCC3,TMEM165,YBX1,AMACR,RPF2,SGK3,SPCS2,TFAM,TMEM33,GTF2H3,PSIP1,ALDH7A1,VPS25,MORN4,MRPL16,SF3A3,VCL,ANP32E,DUSP16,DYRK1A,HELZ,TIPRL,TARS3,MYDGF,FRA10AC1,RAPGEF4,MTIF2,RIOK2,LMNA,NMRK2,EGFR,SPR,MRTO4,DLL1,ATXN10,NOTUM,CREM,SVEP1,SCN2B,RC3H2,XIRP1,SMARCD3,CSTPP1,CCDC136,ZZEF1,ALPK3,WHAMM,GPRC5A,ANKRD37,S100A10,RALGPS1,JPH4,FXYD1,AGO3,ABTB3,EP400,PYROXD2,ZCCHC3,SLC9A6,ACSBG2,PFKFB3,ATF3,TSPAN7,NEU3,NR4A3,FAM110B,LRCH3,OTUD7B,TMEM45B,CEP19,PPRC1,CHRDL1,IQSEC2,PPP4R3B,ARHGAP39,CPNE4,SERINC4,EGR1,CLIC6,HMBOX1,PDE6C,ZC3H12C,PIWIL1,GLP2R,SSPO,UROC1,LGSN,GABRA5,CD9,CDHR4,TOPAZ1,ADCY7,PAMR1,MSMP,PRKCB,ADAM20 |
| KEGG | Endocrine resistance | KEGG:01522 | FALSE | 0.00603033 | 2.21965873 | 93 | 71 | 6 | 9330 | NCOR1,FOS,CCND1,EGFR,DLL1,ADCY7 |
| KEGG | Parathyroid hormone synthesis, secretion and action | KEGG:04928 | FALSE | 0.01384183 | 1.8588065 | 108 | 71 | 6 | 9330 | CASR,FOS,EGFR,EGR1,ADCY7,PRKCB |
| KEGG | Chemical carcinogenesis - receptor activation | KEGG:05207 | FALSE | 0.02471002 | 1.60712688 | 225 | 71 | 8 | 9330 | FOS,MGST1,GSTO1,CCND1,EGFR,DLL1,ADCY7,PRKCB |
| KEGG | Hepatocellular carcinoma | KEGG:05225 | FALSE | 0.02879547 | 1.54067578 | 174 | 71 | 7 | 9330 | DVL1,MGST1,GSTO1,CCND1,EGFR,SMARCD3,PRKCB |
| TF | Factor: Kaiso; motif: GCMGGGRGCRGS | TF:M03876 | FALSE | 0.0027102 | 2.56699863 | 11106 | 153 | 107 | 21628 | AOX3,KCNH7,CDIPT,FOXA3,ESM1,INTU,FAAH,SLC25A22,DVL1,TTLL11,HSPG2,CHI3L1,HAUS8,NCOR1,FOS,CA2,WASF3,MTHFD2,FAM131C,CDK5RAP1,AGPAT5,METTL9,GKAP1,GSTO1,GATAD1,CDK2AP2,PRKRIP1,CCND1,CCNL1,EEF1AKMT2,TRIM27,BRCC3,TMEM165,YBX1,AMACR,TFAM,TMEM33,PSIP1,ALDH7A1,VPS25,SF3A3,VCL,ANP32E,DUSP16,DYRK1A,HELZ,TIPRL,TARS3,MYDGF,FRA10AC1,RAPGEF4,MTIF2,SNX29,LMNA,EGFR,MRTO4,DLL1,ATXN10,NOTUM,CREM,SCN2B,METTL27,RC3H2,SMARCD3,CCDC136,ALPK3,ASB12,WHAMM,GPRC5A,ANKRD37,S100A10,RALGPS1,JPH4,EP400,PYROXD2,ZCCHC3,SLC9A6,ACSBG2,PFKFB3,TSPAN7,NR4A3,LRCH3,OTUD7B,TMEM45B,CEP19,FBXO41,PPRC1,IQSEC2,PPP4R3B,ARHGAP39,CPNE4,SERINC4,EGR1,CLIC6,HMBOX1,PDE6C,ZC3H12C,GLP2R,SSPO,UROC1,ZBTB8OS,GABRA5,CD9,TOPAZ1,ADCY7,PAMR1,PRKCB |
| TF | Factor: Pax-5; motif: RRMSWGANWYCTNRAGCGKRACSRYNSM | TF:M00144 | FALSE | 0.00470868 | 2.32710038 | 11967 | 153 | 112 | 21628 | TAAR1,KCNH7,SLC7A2,CDIPT,FOXA3,ZDHHC11,SLC2A9,INTU,WDR49,FAAH,CLEC5A,SLC25A22,CALR,DVL1,TTLL11,HSPG2,HAUS8,NCOR1,CA2,ACP6,SPTBN5,MTHFD2,FAM131C,RNF213,AGPAT5,METTL9,GKAP1,GSTO1,CDK2AP2,PRKRIP1,CCND1,CCNL1,TRIM27,TMEM165,YBX1,AMACR,SGK3,TFAM,TMEM33,PSIP1,VPS25,MORN4,MRPL16,SF3A3,VCL,ANP32E,DUSP16,HELZ,TIPRL,TARS3,MYDGF,FRA10AC1,RAPGEF4,PBDC1,MTIF2,SNX29,RIOK2,NMRK2,EGFR,SPR,MRTO4,DLL1,ATXN10,NOTUM,CREM,SVEP1,SCN2B,METTL27,RC3H2,XIRP1,SMARCD3,CSTPP1,CCDC136,ZZEF1,GPRC5A,ANKRD37,S100A10,RALGPS1,JPH4,AGO3,ABTB3,EP400,ZCCHC3,ACSBG2,PFKFB3,TSPAN7,NEU3,NR4A3,LRCH3,OTUD7B,TMEM45B,CEP19,FBXO41,PPRC1,CHRDL1,IQSEC2,PPP4R3B,ARHGAP39,CPNE4,SERINC4,EGR1,CLIC6,HMBOX1,ZC3H12C,PIWIL1,SSPO,ZBTB8OS,GABRA5,CD9,CDHR4,MSMP,PRKCB |
| TF | Factor: ZF5; motif: NRNGNGCGCGCWN | TF:M00333 | FALSE | 0.00496272 | 2.30428005 | 14397 | 153 | 127 | 21628 | KCNH7,SLC7A2,CDIPT,FOXA3,ZDHHC11,INTU,FAAH,SLC25A22,TAP2,CALR,DVL1,TTLL11,HSPG2,HAUS8,CASR,NCOR1,FOS,TBC1D7,CA2,WASF3,SPTBN5,MTHFD2,FAM131C,RNF213,CDK5RAP1,PSMA4,AGPAT5,METTL9,GKAP1,MGST1,GSTO1,GATAD1,CDK2AP2,PRKRIP1,CCND1,CCNL1,EEF1AKMT2,TRIM27,BRCC3,TMEM165,YBX1,AMACR,RPF2,SGK3,TFAM,TMEM33,GTF2H3,PSIP1,ALDH7A1,VPS25,MORN4,MRPL16,SF3A3,VCL,ANP32E,DUSP16,DYRK1A,HELZ,TIPRL,TARS3,MYDGF,FRA10AC1,RAPGEF4,PBDC1,MTIF2,SNX29,RIOK2,LMNA,NMRK2,EGFR,SPR,MRTO4,DLL1,ATXN10,NOTUM,CREM,SVEP1,SCN2B,RC3H2,SMARCD3,CCDC136,ZZEF1,ALPK3,ASB12,WHAMM,GPRC5A,ANKRD37,S100A10,RALGPS1,JPH4,FXYD1,AGO3,ABTB3,EP400,PYROXD2,ZCCHC3,SLC9A6,ACSBG2,PFKFB3,TSPAN7,NEU3,NR4A3,LRCH3,OTUD7B,TMEM45B,CEP19,FBXO41,PPRC1,CHRDL1,IQSEC2,PPP4R3B,ARHGAP39,CPNE4,EGR1,CLIC6,HMBOX1,LRRC74B,ZC3H12C,PIWIL1,GLP2R,ZBTB8OS,LGSN,GABRA5,CD9,CDHR4,PAMR1,PRKCB |
| TF | Factor: FOXN4; motif: NNWANNCGWMCGCGTCNNNNMT; match class: 1 | TF:M04662_1 | FALSE | 0.0251654 | 1.59919612 | 12761 | 153 | 115 | 21628 | TAAR1,KCNH7,SLC7A2,CDIPT,FOXA3,TIE1,INTU,FAAH,CLEC5A,SLC25A22,CALR,DVL1,TTLL11,HSPG2,HAUS8,CASR,NCOR1,FOS,TBC1D7,CA2,WASF3,ACP6,SPTBN5,MTHFD2,FAM131C,CDK5RAP1,PSMA4,AGPAT5,METTL9,GKAP1,MGST1,GSTO1,GATAD1,CDK2AP2,PRKRIP1,CCND1,CCNL1,EEF1AKMT2,TRIM27,BRCC3,TMEM165,YBX1,AMACR,RPF2,SGK3,TFAM,TMEM33,GTF2H3,PSIP1,ALDH7A1,VPS25,MORN4,MRPL16,SF3A3,VCL,ANP32E,DUSP16,DYRK1A,HELZ,TIPRL,TARS3,MYDGF,FRA10AC1,RAPGEF4,PBDC1,MTIF2,SNX29,RIOK2,LMNA,NMRK2,EGFR,MRTO4,DLL1,ATXN10,NOTUM,CREM,SVEP1,METTL27,RC3H2,SMARCD3,CCDC136,ZZEF1,WHAMM,GPRC5A,ANKRD37,RALGPS1,JPH4,AGO3,ABTB3,EP400,ZCCHC3,SLC9A6,PFKFB3,TSPAN7,NR4A3,LRCH3,OTUD7B,TMEM45B,CEP19,FBXO41,PPRC1,IQSEC2,PPP4R3B,ARHGAP39,EGR1,CLIC6,LRRC74B,ZC3H12C,PIWIL1,ZBTB8OS,GABRA5,CD9,TOPAZ1,ADCY7,PRKCB |
| TF | Factor: Kaiso; motif: GCMGGGRGCRGS; match class: 1 | TF:M03876_1 | FALSE | 0.03032765 | 1.51816129 | 5835 | 153 | 65 | 21628 | KCNH7,TTLL11,FOS,CA2,WASF3,MTHFD2,FAM131C,AGPAT5,METTL9,GKAP1,GSTO1,GATAD1,CDK2AP2,CCND1,CCNL1,TRIM27,YBX1,AMACR,TFAM,TMEM33,PSIP1,ALDH7A1,VPS25,VCL,DUSP16,DYRK1A,HELZ,TIPRL,TARS3,MYDGF,FRA10AC1,RAPGEF4,LMNA,MRTO4,DLL1,ATXN10,NOTUM,CREM,SMARCD3,CCDC136,WHAMM,GPRC5A,ANKRD37,RALGPS1,PYROXD2,ZCCHC3,SLC9A6,ACSBG2,PFKFB3,TSPAN7,LRCH3,CEP19,FBXO41,IQSEC2,PPP4R3B,ARHGAP39,CPNE4,SERINC4,PDE6C,ZC3H12C,ZBTB8OS,GABRA5,ADCY7,PAMR1,PRKCB |
| TF | Factor: ZF5; motif: NRNGNGCGCGCWN; match class: 1 | TF:M00333_1 | FALSE | 0.03948453 | 1.40357298 | 11183 | 153 | 104 | 21628 | KCNH7,SLC7A2,CDIPT,FOXA3,INTU,FAAH,SLC25A22,CALR,DVL1,TTLL11,HSPG2,HAUS8,CASR,NCOR1,FOS,TBC1D7,CA2,WASF3,MTHFD2,FAM131C,CDK5RAP1,PSMA4,AGPAT5,METTL9,GKAP1,MGST1,GATAD1,CDK2AP2,PRKRIP1,CCND1,CCNL1,TRIM27,BRCC3,TMEM165,YBX1,AMACR,RPF2,SGK3,TFAM,TMEM33,GTF2H3,PSIP1,ALDH7A1,SF3A3,VCL,ANP32E,DUSP16,DYRK1A,HELZ,TIPRL,TARS3,MYDGF,RAPGEF4,PBDC1,MTIF2,SNX29,RIOK2,LMNA,NMRK2,EGFR,SPR,MRTO4,DLL1,ATXN10,NOTUM,CREM,SVEP1,SCN2B,SMARCD3,CCDC136,ZZEF1,ASB12,WHAMM,GPRC5A,ANKRD37,S100A10,JPH4,FXYD1,AGO3,ABTB3,EP400,ZCCHC3,SLC9A6,PFKFB3,TSPAN7,NR4A3,LRCH3,OTUD7B,CEP19,FBXO41,PPRC1,CHRDL1,IQSEC2,PPP4R3B,ARHGAP39,EGR1,CLIC6,HMBOX1,LRRC74B,ZC3H12C,ZBTB8OS,GABRA5,CD9,PRKCB |
| TF | Factor: Sp5; motif: RNGGRGGNGGRGNNGGGGGAGGRG; match class: 1 | TF:M10376_1 | FALSE | 0.04390022 | 1.35753325 | 1940 | 153 | 30 | 21628 | KCNH7,FOS,FAM131C,METTL9,GATAD1,CCND1,TRIM27,SGK3,TFAM,PSIP1,VPS25,DUSP16,DYRK1A,HELZ,NMRK2,EGFR,DLL1,ATXN10,SCN2B,RC3H2,CCDC136,ASB12,RALGPS1,PYROXD2,SLC9A6,TSPAN7,OTUD7B,FBXO41,PPP4R3B,EGR1 |

**Table S13. Notes regarding scaffold nomenclature differences between the annotation, DNA Zoo assembly, and GENESPACE riparian plot**

| Annotation | DNA Zoo Assembly | Synteny Riparian Plot |
| --- | --- | --- |
| scaffold_1 | HiC-scaffold_1 | Ut1 |
| scaffold_2 | HiC-scaffold_2 | Ut2 |
| scaffold_3 | HiC-scaffold_3 | Ut3 |
| scaffold_4 | HiC-scaffold_4 | Ut4 |
| scaffold_5 | HiC-scaffold_5 | Ut5 |
| scaffold_6 | HiC-scaffold_6 | Ut6 |
| scaffold_7 | HiC-scaffold_7 | Ut7 |
| scaffold_8 | HiC-scaffold_8 | Ut8 |
| scaffold_9 | HiC-scaffold_9 | Ut9 |
| scaffold_10 | HiC-scaffold_10 | Ut10 |
| scaffold_11 | HiC-scaffold_11 | Ut11 |
| scaffold_12 | HiC-scaffold_12 | Ut12 |
| scaffold_13 | HiC-scaffold_13 | Ut13 |
| scaffold_15 | HiC-scaffold_14 | Ut15 |
| scaffold_16 | HiC-scaffold_15 | Ut16 |
| scaffold_17 | HiC-scaffold_16 | Ut17 |
| scaffold_14 | HiC-scaffold_17 | Ut14 |

**Table S14. Mean and median read depth per 100 kb window per scaffold**

| **scaffold** | **mean_depth** | **median_depth** |
| --- | --- | --- |
| **scaffold_1** | 36.6953966552005 | 36.6187 |
| **scaffold_2** | 37.2214642041527 | 37.05684 |
| **scaffold_3** | 36.4837125191489 | 36.39988 |
| **scaffold_4** | 36.4718529198978 | 36.451625 |
| **scaffold_5** | 36.4473928639599 | 36.33496 |
| **scaffold_6** | 37.1343920877307 | 36.93281 |
| **scaffold_7** | 38.5927446163212 | 38.189 |
| **scaffold_8** | 40.2849980190769 | 39.20494 |
| **scaffold_9** | 38.9180558503145 | 38.428605 |
| **scaffold_10** | 39.0527818645886 | 37.933165 |
| **scaffold_11** | 38.2924916340514 | 37.54396 |
| **scaffold_12** | 41.4452302555663 | 41.72173 |
| **scaffold_13** | 40.5486973983936 | 39.64977 |
| **scaffold_14** | 22.4397078819271 | 20.67563 |
| **scaffold_15** | 40.4157093030939 | 41.06396 |
| **scaffold_16** | 42.6244697720122 | 41.962585 |
| **scaffold_17** | 42.1356757220497 | 42.93401 |
| **scaffold_18** | 18.3826739418095 | 19.55602 |
| **scaffold_19** | 30.4109528677778 | 27.12306 |
| **scaffold_20** | 24.7388552777778 | 25.08798 |
| **scaffold_21** | 26.1226470675 | 24.243955 |
| **scaffold_22** | 21.0663636971429 | 19.14762 |
| **scaffold_23** | 19.4760706016667 | 19.46188 |
| **scaffold_24** | 30.3861950633333 | 26.13911 |
| **scaffold_25** | 7.0845373152 | 6.09847 |
| **scaffold_26** | 30.5960536 | 29.6181722 |
| **scaffold_27** | 21.5685232575 | 23.06591 |
| **scaffold_28** | 18.2298841133333 | 18.97875 |
| **scaffold_29** | 14.5751709166667 | 13.49967275 |
| **scaffold_30** | 19.4475000903333 | 20.4486 |
| **scaffold_31** | 26.1929641466667 | 25.40657244 |
| **scaffold_32** | 19.7082626466667 | 21.26293 |
| **scaffold_33** | 18.872474465 | 18.872474465 |
| **scaffold_34** | 28.4066433 | 28.4066433 |
| **scaffold_35** | 27.909071415 | 27.909071415 |
| **scaffold_36** | 30.07644715 | 30.07644715 |
| **scaffold_37** | 10.1435365705 | 10.1435365705 |
| **scaffold_38** | 100.71581695 | 100.71581695 |
| **scaffold_39** | 21.27733582 | 21.27733582 |
| **scaffold_40** | 22.099726185 | 22.099726185 |
| **scaffold_41** | 23.77906292 | 23.77906292 |
| **scaffold_42** | 26.4106489 | 26.4106489 |
| **scaffold_43** | 19.44516478 | 19.44516478 |
| **scaffold_44** | 14.34421293 | 14.34421293 |
| **scaffold_45** | 26.86524533 | 26.86524533 |
| **scaffold_46** | 102.5931953 | 102.5931953 |
| **scaffold_47** | 14.22878844 | 14.22878844 |
| **scaffold_48** | 9.971294307 | 9.971294307 |
| **scaffold_49** | 13.85298124 | 13.85298124 |
| **scaffold_50** | 12.15940991 | 12.15940991 |
| **scaffold_51** | 22.87152999 | 22.87152999 |
| **scaffold_52** | 18.60140325 | 18.60140325 |
| **scaffold_53** | 7.689215654 | 7.689215654 |
| **scaffold_54** | 9.038570505 | 9.038570505 |
| **scaffold_55** | 9.43532264 | 9.43532264 |
| **scaffold_56** | 17.6584289 | 17.6584289 |
| **scaffold_57** | 88.08545872 | 88.08545872 |
| **scaffold_58** | 97.84252768 | 97.84252768 |
| **scaffold_59** | 8.324634156 | 8.324634156 |
| **scaffold_60** | 18.16961563 | 18.16961563 |
| **scaffold_61** | 103.8928077 | 103.8928077 |
| **scaffold_62** | 14.08884835 | 14.08884835 |
| **scaffold_63** | 9.989305171 | 9.989305171 |
| **scaffold_64** | 9.95546228 | 9.95546228 |
| **scaffold_65** | 20.70776777 | 20.70776777 |
| **scaffold_66** | 35.49093555 | 35.49093555 |
| **scaffold_67** | 13.89117633 | 13.89117633 |
| **scaffold_68** | 15.3164091 | 15.3164091 |
| **scaffold_69** | 7.133077989 | 7.133077989 |
| **scaffold_70** | 5.08479399 | 5.08479399 |
| **scaffold_71** | 13.31076255 | 13.31076255 |
| **scaffold_72** | 13.41065948 | 13.41065948 |
| **scaffold_73** | 5.094368208 | 5.094368208 |
| **scaffold_74** | 4.703864037 | 4.703864037 |
| **scaffold_75** | 21.46220188 | 21.46220188 |
| **scaffold_76** | 7.191967257 | 7.191967257 |
| **scaffold_77** | 10.04763333 | 10.04763333 |
| **scaffold_78** | 38.95240651 | 38.95240651 |
| **scaffold_79** | 15.15218294 | 15.15218294 |
| **scaffold_80** | 17.15115098 | 17.15115098 |
| **scaffold_81** | 8.78327345 | 8.78327345 |
| **scaffold_82** | 8.728627638 | 8.728627638 |
| **scaffold_83** | 5.910064753 | 5.910064753 |
| **scaffold_84** | 235.7077181 | 235.7077181 |
| **scaffold_85** | 15.12450823 | 15.12450823 |
| **scaffold_86** | 15.42397113 | 15.42397113 |
| **scaffold_87** | 14.89737492 | 14.89737492 |
| **scaffold_88** | 123.1906485 | 123.1906485 |
| **scaffold_89** | 10.44847133 | 10.44847133 |
| **scaffold_90** | 38.63128687 | 38.63128687 |
| **scaffold_91** | 9.291073412 | 9.291073412 |
| **scaffold_92** | 3.282769895 | 3.282769895 |
| **scaffold_93** | 3.434877516 | 3.434877516 |
| **scaffold_94** | 4.724008607 | 4.724008607 |
| **scaffold_95** | 96.82834208 | 96.82834208 |
| **scaffold_96** | 3.953235146 | 3.953235146 |
| **scaffold_97** | 17.90641673 | 17.90641673 |
| **scaffold_98** | 15.04204484 | 15.04204484 |
| **scaffold_99** | 12.7434 | 12.7434 |
| **scaffold_100** | 63.84554096 | 63.84554096 |
| **scaffold_101** | 4.03 | 4.03 |
| **scaffold_102** | 25.83466667 | 25.83466667 |
| **scaffold_103** | 23.1945 | 23.1945 |
| **scaffold_104** | 56.364 | 56.364 |
